# Allopatric speciation is more prevalent than parapatric ecological divergence in a recent high-Andean diversification (Asteraceae: *Linochilus*)

**DOI:** 10.1101/868216

**Authors:** Oscar M. Vargas, Santiago Madriñán, Beryl B. Simpson

## Abstract

Elucidating how species accumulate in diversity hotspots is an ongoing debate in evolutionary biology. The páramo, in the Northern Andes, has remarkably high indices of plant diversity, endemicity, and diversification rates. A hypothesis for explaining such indices is that allopatric speciation is high in the páramo given its island-like distribution; an alternative hypothesis is that the altitudinal gradient of the Andean topography provides a variety of niches that drive vertical parapatric ecological speciation. A formal test for evaluating the relative roles of allopatric speciation and parapatric ecological divergence is lacking. The main aim of our study is to test which kind of speciation is more common in an endemic páramo genus. We developed a framework incorporating phylogenetics, species’ distributions, and a morpho-ecological trait (leaf area) to compare sister species and infer whether allopatric or parapatric ecological divergence caused their speciation. We applied our framework to the species-rich genus *Linochilus* (63 spp.) and found that the majority of recent speciation events in it (12, 80%) have been driven by allopatric speciation, while a smaller fraction (1, 6.6%) is attributed to parapatric ecological divergence; two pairs produced inconclusive results (13.3%). We conclude that páramo autochthonous diversification is primarily driven by allopatric speciation.

## INTRODUCTION

Alexander von Humboldt’s and Aimé Bonpland’s *Tableau Physique des Andes et Pays Voisins* illustrated how plant species are assembled from lowlands to high altitudes in the tropical Andes (von Humboldt and Bonpland 1805). This representation visualized for the first time how diversity changes locally with elevation, setting a seminal starting point for biodiversity studies. Today, 217 years after the first publication of the *Essay on the Geography of Plants* (von Humboldt and Bonpland 1805), scientists have mapped species richness around the globe and have identified numerous hot-spots of biodiversity. Understanding how species accumulation occurs in such hotspots is pivotal for the fields of evolutionary biology and biogeography.

Because biodiversity hotspots often coincide with areas of topographic complexity (Barthlott et al. 1996; Myers et al. 2000; Mutke and Barthlott 2005; Jenkins et al. 2013) geographical isolation and ecological opportunity are typically cited to explain species richness. In mountain systems, valleys and canyons act as landscape barriers for organisms (Janzen 1967; van der Hammen and Cleef 1986; Muños-Ortiz et al. 2015) promoting allopatric speciation (vicariant or peripatric) via geographical isolation, while slopes and ecological gradients can drive parapatric speciation via ecological divergence (Hughes and Atchison 2015; Pyron et al. 2015). Vicariant speciation occurs when a mother species that is distributed broadly is divided into two daughter populations, which speciate via subsequent independent evolution. Glacial and interglacial cycles are often assumed to have caused these vicariant events (van der Hammen and Cleef 1986; Carstens and Knowles 2007, Flantua et al. 2019). Peripatric speciation occurs when a dispersal event from a source lineage to a new, previously uncolonized area takes place (founder speciation event). If the newly colonized area is geographically isolated enough, then lineages become reproductively isolated with one another, resulting in progenitor and derivative lineage as different species (Coyne and Orr 2004). Parapatric speciation takes place when a continuous population that is distributed along an ecological gradient (e.g. an altitudinal gradient) is subdivided in two or more subpopulations that locally adapt to different niches (e.g. lower vs. higher elevations); the initial subpopulations become independent species by means of ecological differentiation, non-random mating, and the formation of reproductive barriers (Givnish 1997; Schluter 2000; Simpson 1953).

Under a scenario of allopatric speciation, recently diverged species should be similar to each other thanks to phylogenetic conservatism (Fig. 1A). In the context of sky-islands, sister species that arose because of geographic speciation should occupy similar niches in different islands, presenting similar ecological characteristics (Pyron et al. 2015). Alternatively, under a scenario of parapatric ecological speciation, sister species are expected to occupy different niches within an island, presenting a signature of ecological divergence (Fig. 1D) (Pyron et al. 2015). Therefore, it is possible to test the predictions of allopatric speciation and parapatric ecological divergence in sky-islands by studying the distribution and ecological traits of sister taxa.

**FIGURE 1.**
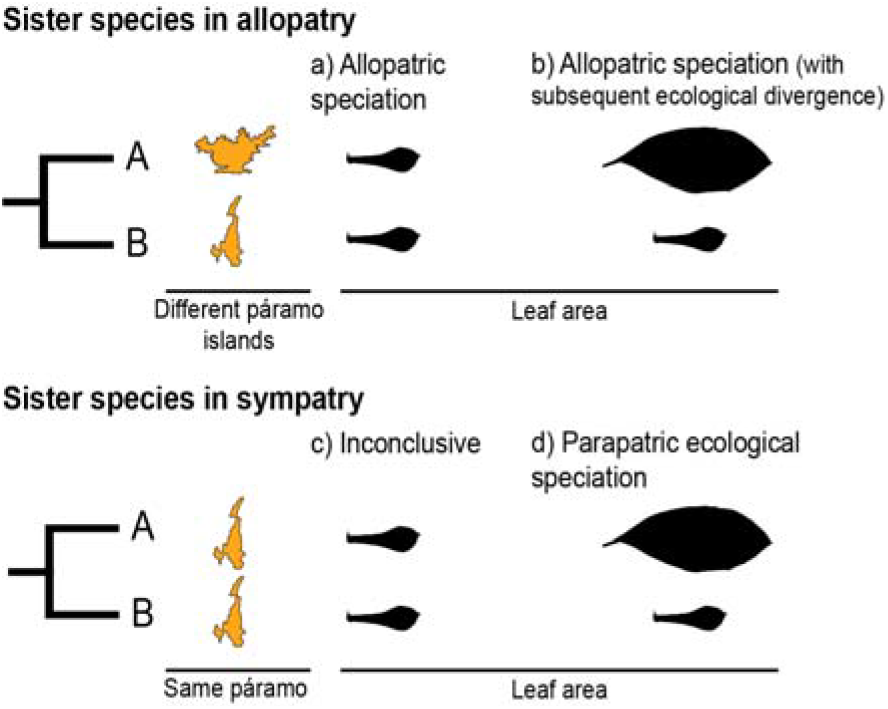
Hypothetical scenarios for speciation in sister species. a) Allopatric speciation event in which geographical isolation resulted in two species living in separate páramo islands (indicated by island shape and color) and occupying similar niches (as indicated by similar leaf shapes). b) Allopatric speciation event in which geographical isolation resulted in two species living in separate páramo islands occupying different niches as indicated by different leaf types. c) A speciation event in which the reason for divergence is inconclusive, sister species inhabit the same island and have similar leaf shapes. d) A parapatric ecological speciation event in which sister species evolved different leaf areas in response to selection to different niches on the same páramo island.

The Páramo, a high-altitude ecosystem found above the timberline in the Northern Andes (the Andes of Ecuador, Colombia, and Venezuela) (Cuatrecasas 1968; Luebert and Weigend 2014; Weigend 2002), provides an ideal scenario to test how speciation happens in high elevation and island-like biodiversity hotspots (Fig. 2). With ca. ~3,400 species of vascular plants (Luteyn 1999), of which 60% to 100% are estimated to be endemic (Luteyn 1992; Madriñán et al. 2013), and particularly high diversification rates (Madriñán et al. 2013), the páramo is considered the most species rich ecosystem of the world’s tropical montane regions (Sklenář et al. 2014). Unlike other South American high-altitude areas farther south (i.e. Perú and Chile), the páramo is characterized by abundant precipitation throughout the year, and is therefore defined by both high altitude and humidity. It has been estimated that the páramo originated when the high altitudes (>3,000 m) of the Northern Andes emerged as a result of rapid uplift 2–4 Ma (van der Hammen and Cleef 1986; Gregory-Wodzicki 2000). The fragmented coverage of the paramo (Fig. 2) and its altitudinal gradient (~3000–4500 m) are hypothesized to have acted as drivers of diversification by promoting both allopatric speciation (vicariant or peripatric) via geographical isolation (van der Hammen and Cleef 1986) and parapatric speciation via ecological divergence (Hughes and Atchison 2015, Nevado et al. 2018). Despite efforts documenting Andean diversification (e.g. Nürk et al. 2015; Uribe-Convers and Tank 2015; Lagomarsino et al. 2016; Pérez-Escobar et al. 2017) a formal test to quantify the relative prevalence of allopatric versus parapatric ecological speciation of taxa in the region is lacking.

**FIGURE 2.**
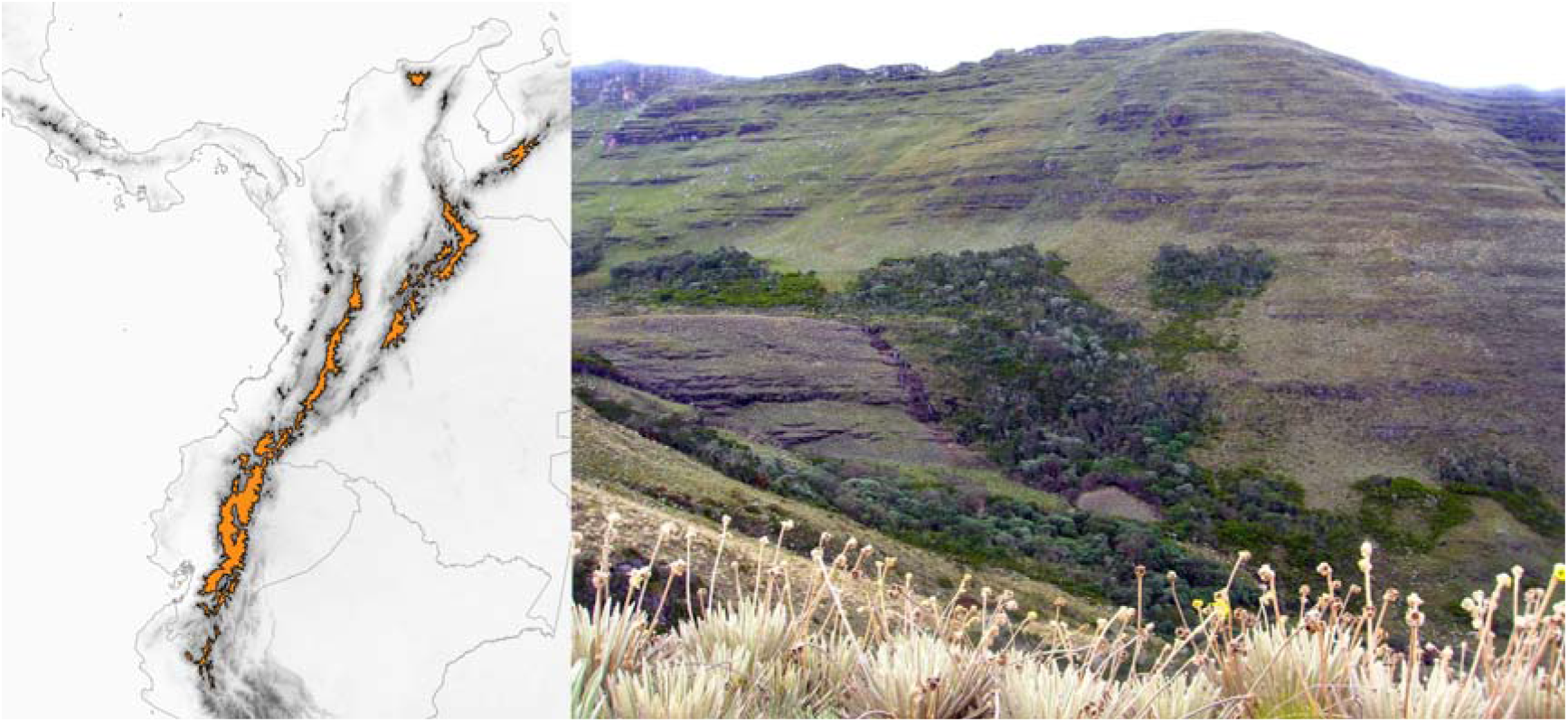
Approximate geographical extent of the Páramo ecosystem (left) on the Northern Andes. Páramo la Rusia, Boyacá, Colombia (right); notice how high the Andean forest interdigitates with the Páramo along the creek.

To quantify the relative contributions of allopatric and parapatric speciation in the páramo, we used a comparative framework that combines phylogenetic, geographical, and ecological information. We applied this approach to *Linochilus*, a genus restricted to the páramo and the upper boundary of the cloud forest (Blake 1928; Cuatrecasas 1969; Vargas 2011, 2018; Vargas et al. 2017; Saldivia et al. 2019). *Lichochilus’* phylogeny was recently inferred using high-throughput sequencing (Vargas et al. 2017), and a taxonomic monograph is almost complete for the genus by OMV.

The main aim of our study is to ascertain which kind of speciation, allopatric versus parapatric ecological divergence, is more common in this this plant genus that inhabits an island-like system and its lower boundary. We specifically aim to 1) quantify the relative contribution of allopatric and parapatric ecological speciation in the divergence of sister taxa in *Linochilus*, and 2) to pinpoint the geographical origin of the *Linochilus* genus, since its presence in the páramo and the upper boundary of the cloud forest suggest and ecological transition from paramo to the forest (Vargas & Madriñán 2012).

## MATERIALS AND METHODS

R Code, input and control files, and bioregionalization maps are available at https://bitbucket.org/XXXXXXXX/linochilus

### FOCAL CLADE

We used *Linochilus* (Asteraceae) because of our recent comprehensive knowledge about its phylogeny (Vargas et al. 2017), distribution, and taxonomy of its species (Cuatrecasas 1969; Vargas 2011; Vargas 2018). Additionally, it is almost entirely restricted to the páramo, although some species dwell in the upper boundary of the cloud forest due to a downslope colonization event (Vargas and Madriñán 2012). The genus contains 63 species distributed in the disjunct mountains of the Talamanca Cordillera (Costa Rica), the Sierra Nevada de Santa Marta (Colombia), and the Northern Andes (Colombia, Venezuela, and Ecuador), exhibiting a variety of woody habits from decumbent subshrubs only 10 cm tall to small trees 6 m tall (Cuatrecasas 1969; Vargas 2018). Growth form and leaf area of *Linochilus* species are associated with the habitat they occupy—shrubs and decumbent subshrubs with microphyllous leaves inhabit the open páramo, while small trees with broad leaves reside at lower altitudes in the upper, more humid, edge of Andean forest (Vargas and Madriñán 2012). *Linochilus* was recently segregated from *Diplostephium*, a genus with similar morphology and ecology that primarily inhabits the Central Andes (Vargas 2018, Saldivia et al. 2019).

### SISTER SPECIES COMPARISONS

To measure the relative contribution of allopatric speciation and parapatric ecological divergence on recent speciation events, we compared the geographical distribution (Appendix S1) and the leaf areas (Appendix S2) of species shown to be sister in the phylogeny of *Linochilus* (Vargas et al. 2017). We used leaf area as a proxy to evaluate ecological divergence between sister species. Leaf area is a functional character that varies with the eco-physiological pressures of a species’ niche, reflecting its adaptation to water availably, irradiance, and elevation (Givnish 1987), therefore providing a proxy for ecological niche. For example, *L. antioquensis* inhabits the upper Andean forest and has an average leaf area of 848.4 mm^2^. In contrast, *L. phylicoides*, dwells in the physiologically dry open páramo with plenty of access to sunlight, having an average leaf area of 4.0 mm^2^. When possible, we measured the area of 30 leaves from six different individuals for each species (Appendix S3). We scanned the leaves at 600 dpi from herbarium material belonging to ANDES, TEX, and US herbaria. Each leaf was outlined using PHOTOSHOP CS4 (Adobe Systems, San Jose, California) as a single white and black image. We then used the R package MOMOCS (Bonhomme et al. 2014) to calculate the area of each leaf from the images created in the previous step. We performed a Wilcoxon signed-rank tests of the log-transformed leaf areas between sister species using R (R Core Team 2016). We also used the Wilcoxon signed-rank test to compare the leaf area between clades. It is important to note that this test considers the distribution of values for leaf areas in each species (instead of just the average) and it is designed to compare non-independent samples.

We scored a pair as allopatric when they inhabited non-overlapping páramo islands (mountaintops) and when they inhabited different slopes of the same mountain (this pattern is possible for species that inhabit the upper boundaries of the Andean forest). We scored a pair as sympatric when their distribution overlapped at least one páramo island— in other words, the presence of the two sisters in a single páramo island will make them sympatric. We followed the paramo delineation of Londoño et al. 2014. In addition to using paramo islands for scoring species as allopatric or sympatric, we calculated range overlap and range asymmetry between sisters by overlaying occurrences in a grid following Vargas et al. (2020). We used two grid sizes, 0.05 and 0.1 decimal degrees, corresponding to 33 and 131 Km2 respectively. With the grid-ranges of sister species, we calculated the sisters’ overlap as the area occupied by both sisters divided by the summed area of the smaller ranged sister; an overlap of 0 indicates full allopatry while an overlap of 1 indicates that the smaller-ranged sister is found solely within the range of its sister (Barraclough and Vogler 2000; Fitzpatrick and Turelli 2006). Range asymmetry was calculated as the area occupied by the larger-ranged sister divided by the area of its sister (Fitzpatrick and Turelli 2006). The aforementioned calculations were made on R using the “raster” package v. 3.5-21 (Hijmans 2022). Distributional data were extracted from curated COL and US herbaria specimens.

We interpreted the results in the following fashion (Fig. 1):

- If a sister-species pair is allopatric and there is no significant difference between their leaf areas, we interpreted this scenario as an event of allopatric speciation driven by geographical isolation, with the reasoning that leaf area had remained similar because either relatively little time had passed since divergence and/or because of niche conservatism. (Fig. 1a; Wiens 2004; Pyron et al. 2015).
- If a sister-species pair is allopatric and their leaf areas are significantly different, we interpreted this scenario as an event of allopatric speciation following geographical isolation (Fig. 1b) in which there was subsequent ecological divergence driven by local adaption (Rundell and Price 2009; Pyron et al. 2015)
- If a sister-species pair is scored as sympatric and their leaf areas are different, we interpreted this scenario as an event of parapatric speciation with ecological divergence (Rundle and Nosil 2005; Rundell and Price 2009) (Fig. 1d).
- If a sister-species pair has overlapping geographical distributions and there is no significant difference between their leaf areas, we interpreted this scenario as inconclusive (Fig. 1c). This pattern could be the result of different processes, e.g., allopatric speciation with no ecological divergence followed by secondary contact (Rundell and Price 2009; Hopkins 2013) or a parapatric speciation event driven by ecological divergence in a trait other than leaf area (e.g., Snaydon and Davies 1976; Silvertown et al. 2005).

We acknowledge that the role of parapatric ecological divergence may be underestimated in this study because we only measured leaf area as an ecological proxy. Ecological traits independent from leaf area can confer the ability to colonize different páramo niches (e.g. Cortés et al. 2018), such as underground eco-physiological adaptations to soils with different water saturation, or physiological adaptations at the anatomical and cellular level. Studying alternative physio-ecological variables in sister species comparisons could shed light on other types of ecological divergence (e.g. adaptations to different moisture in soils, microclimatic preferences).

Our framework assumes that:

1. Leaf area represents a good proxy for the organism’s niche, which is likely true in our focal genus. *Linochilus* leaf size is associated with their habitat, broad and large leaves are present in species that dwell in the upper limit of the Andean forest, medium to small leaves are characteristic of taxa inhabiting the páramo, and microphyllous leaves are found in lineages that inhabit the upper boundary of the páramo (a.k.a. superpáramo) (Cuatrecasas 1969, Vargas & Madriñán 2012).
2. The phylogeny employed represents the true species tree and contains all taxa. The best hypothesis to date for the species tree in *Linochilus* (Vargas et al. 2017, based on double digest restriction associated DNA data) contains only a subset of documented *Linochilus* species (36 of 63). To account for the potential effects of missing species in our phylogeny, we searched the taxonomic literature for the most morphologically similar species (assuming this is most likely its sister species) to those species not sampled in the phylogeny and compared their distributional ranges (Appendix S4). Specifically, we searched for new descriptions of species where authors indicate the most similar taxon to the new species based on key traits. Diagnostic characters in *Linochilus* include, but are not limited to, habit, the size and shape of the leaf, the number of capitula per inflorescence, and several measurements in the corolla of both disk and ray flowers. Alternative sisters based on morphology were added only to species included in the first sister species analysis, aiming to maintain a similar number of pairs and make the results comparable. We provide two sets of results: a) considering only the pairs derived from the phylogeny and b) pairing non-sampled species to their most likely sister species present on the phylogeny (we provide a discussion explaining why we believe this second set of results is more reliable).
3. The páramo developed only in the last 2–4 Ma (van der Hammen and Cleef 1986; Gregory-Wodzicki 2000) and it is therefore reasonable to assume that its carrying capacity for the number of species has not been reached, and therefore extinction rates are low. Extremely high diversification rates support the previous statement (Madriñán et al. 2013)
4. The modern species ranges are representative of past ranges, a reasonable assumption given the recent divergence of sister taxa pairs.

### EVALUATING NICHE CONSERVATISM

We used leaf area as a proxy to infer the ecological niche of *Linochilus* species. To test for phylogenetic signal reflected in the evolution of the leaf area, we averaged log-transformed leaf data and calculated Pagel’s (1999) lambda λ) using the R package PHYTOOLS (Revell 2012). To calculate λ, the observed phylogeny was compared to modified trees in which the internal branches are compressed to various degrees. When λ = 0 the observed data follow a model in which internal branches are completely collapsed (star phylogeny), meaning that the trait evolves independently from the phylogeny. When λ = 1 the observed data follow a model in which the internal branches are not modified, meaning that the evolution of the trait is phylogenetically dependent (Harmon 2018; but see Revell et al. 2008).

### BIOGEOGRAPHIC ANALYSIS

To elucidate the biogeographic history of *Linochilus* species, we performed a historical biogeographic analysis. We defined the biogeographic areas based on the páramo complexes defined by Londoño et al. (2014) which was restricted to Colombian páramos, and added three complexes to completely cover the distribution of *Linochilus* (and the páramo). Contours of páramo areas were edited with QGIS 2.8Wien (QGIS Development Team 2005). Our areas were outlined as follows (areas not included by Londoño et al. (2014) are indicated with a star):

*Northern Páramos (N).* Páramos of the “Sierra Nevada de Santa Marta” and the “Serranía del Perijá.”
*Talamanca* (T).* Páramos located in the Talamanca Cordillera of Central America.
*Mérida* (T).* Páramos located in the Mérida Cordillera of Venezuela.
*Eastern Cordillera (E).* Páramos located in Eastern Cordillera of Colombia.
*Antioquia (A).* Cluster of páramos comprised by areas in the Western and Central Cordilleras of Colombia mostly located in the department of Antioquia, Colombia.
*Western Cordillera (W).* Páramos located in the Western Cordillera of Colombia with the exception of those located in the department of Antioquia, Colombia.
*Central Cordillera (C).* Páramos located in the Central Cordillera of Colombia with the exception of those located in the department of Antioquia, Colombia.
*Southern Páramos (S)*. Páramos located in the Colombian Massif and the Ecuadorian Andes.

We pruned the chronogram of Vargas et al. (2017) to include only *Linochilus* species and used BioGeoBEARS (Matzke 2013) to infer the biogeographic history of the genus. BioGeoBEARS implements different models of ancestral range inference, DEC (Dispersal-Extintion-Cladogenesis; Ree and Smith 2008), DIVALIKE (a likelihood version of Dispersal– Vicariance Analysis; Ronquist 1997), and BAYAREAALIKE (a likelihood implementation of the BAYAREA model; Landis et al. 2013), and it evaluates the addition of the J parameter (Matzke 2014) to each one of the models to account for founder-event speciation (DEC+J, DIVALIKE+J, BAYAREALIKE+J but see Ree and Sanmartín 2018). We opted not to use a constrained model (e.g. limiting the presence areas to time windows based on their inferred history) because the paleoaltitudes of the Northern and Central Andes are still debated (Luebert and Weigend 2014). We assigned areas to tips based on the same occurrence dataset used for the sister species comparisons (Appendix S1).

## RESULTS

### SISTER SPECIES COMPARISONS

Independently of the framework used to infer allopatry/sympatry and the method employed to infer sister species (phylogeny vs. phylogeny supplemented with missing sampling), our results unequivocally reveal a strong signal of geographic isolation and little ecological divergence *Linochilus* sister species (Tables 1–2, Fig 3, Appendix S5–7). When only species included in the Vargas et al. (2017) phylogeny are considered (row 1, Table 1; Appendix S5–7), nine out of 14 sister species pairs (64.3%) are allopatric, with only one of these pairs presenting different leaf areas, suggesting divergence by geographical isolation with little ecological divergence after speciation. A further three species pairs (21.4%) occur sympatrically and have evidence of ecological divergence in the leaf areas, while two cases (14.3%) present inconclusive evidence. When species not sampled in the phylogeny of Vargas et al. (2017) are considered (see Appendix S4) by pairing them to their most likely sampled sister species (row 2, Table 1–2, Fig 3) the contribution of allopatric speciation cases increase to 12 out of 15 (80%), while the number of cases for parapatric ecological divergence decreases to 1 (6.6%); the number of inconclusive cases remains about the same, 2 (13.3%). The aforementioned results are virtually the same to using grids of 0.1 and 0.05 decimal degrees to quantify range overlap (Table 3, Appendix S6). For example, range overlap calculated with 0.1 grid produces the same results as our island framework in the analysis supplemented with missing sampling (Tables 2–3). The most fine-scale analysis, using a grid of 0.05 decimal degrees, flips one of the pairs (*L. romeroi – L. saxatilis*) from sympatry to allopatry (this pair is likely to be parapatric with species occupying different vegetational belts), with this pair being the only conflicting result among the approaches.

**FIGURE 3.**
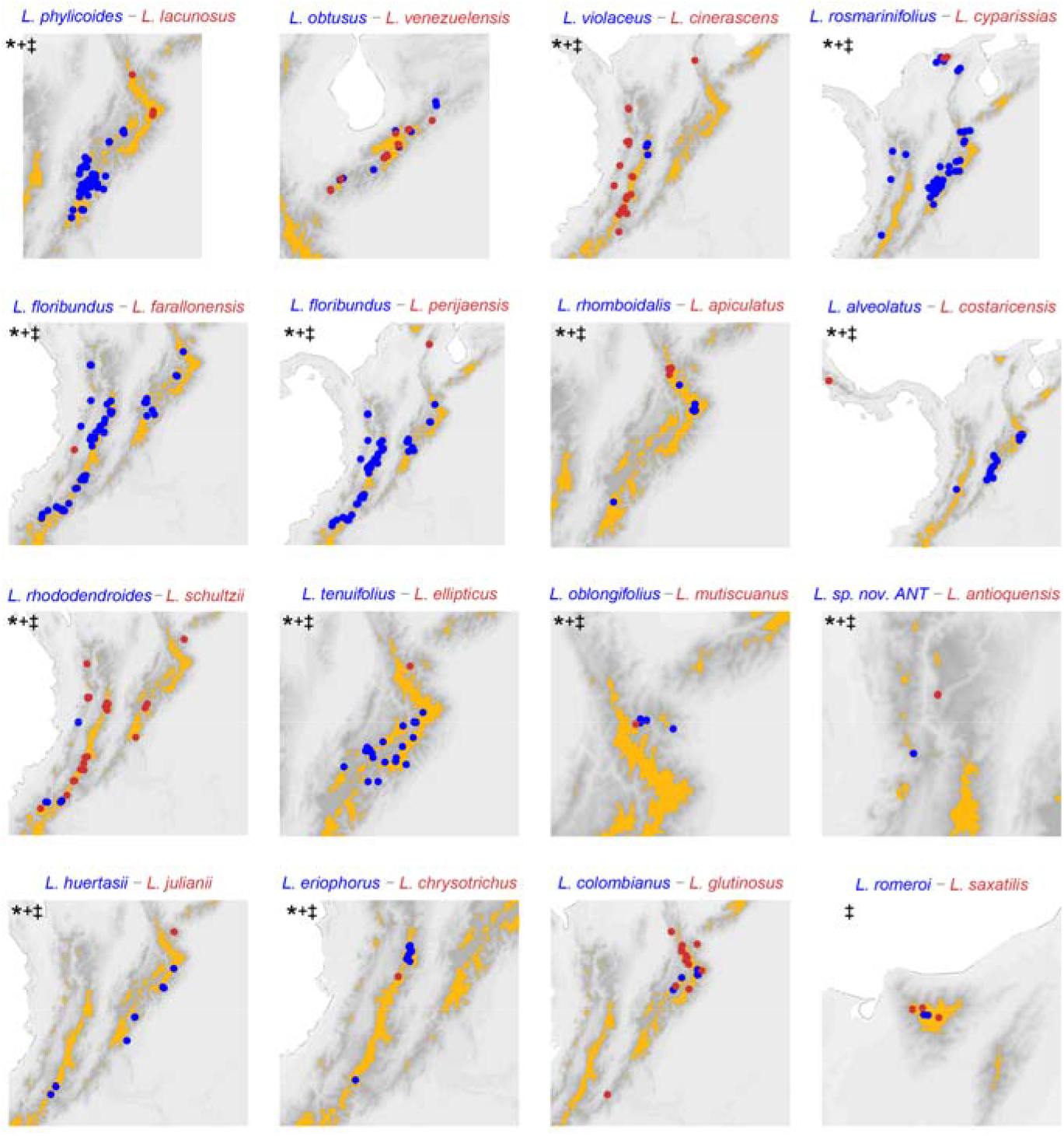
Distribution of sister species based on the phylogeny Vargas et al. (2017) supplemented with missing sampling. Darker pixels indicate higher elevations, orange polygons delineate paramo complexes. **\*** : allopatric based on páramo islands. **+** : allopatric based on 0.1 decimal degree grid analysis. ‡ : allopatric based on 0.05 decimal degree grid analysis. Pairs *L. rosmarinifolius–L. cyparissias* and *L. oblongifolius–L. mutiscuanus*, all inhabiting upper boundary of the cloud forest, are codified as allopatric in the island framework as they inhabit different slopes in the mountains they co-occur.

**TABLE 1.** Relative signal of geographical isolation and parapatric ecological divergence in *Linochilus* based on sister species comparisons.

| Sister sp. pairs based on: | Geographical isolation | Parapatric ecological divergence | Inconclusive |
| --- | --- | --- | --- |
| Phylogeny of Vargas et al. (2017) | 9 (64.3%) | 3 (21.4%) | 2 (14.3%) |
| Phylogeny supplemented with missing sampling | 12 (80%) | 1 (6.6%) | 2 (13.3%) |

**TABLE 2.**
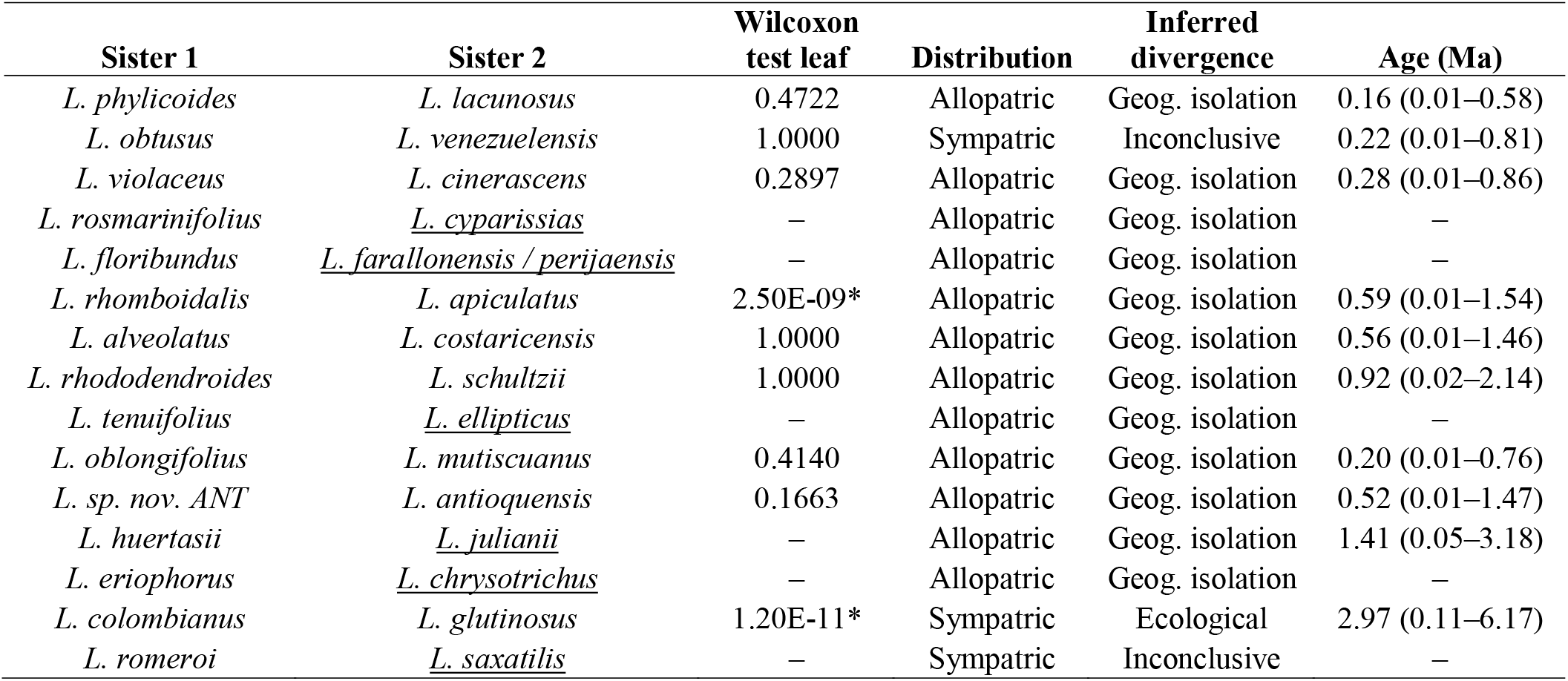
Sister species comparisons based on the phylogeny of Vargas et al. (2017) and supplemented with non-sampled species (underlined). Allopatry and sympatry is determined based on overlapping occurrences on paramo islands as defined by Londoño et al. 2014.

**TABLE 3.** Range overlap calculated using 0.1 and 0.05 decimal degrees grids. *Linochilus* sister species comparisons based on the phylogeny of Vargas et al. (2017) supplemented with non-sampled species (underlined). *L. floribundus* is found twice in the table because there are two hypothesized sisters for it in the taxonomic literature.

| Sister 1 | Sister 2 | Range overlap |  | Range asymmetry |  |
| --- | --- | --- | --- | --- | --- |
|  |  | 0.1 | 0.05 | 0.1 | 0.05 |
| *L. phyllicoides* | *L. lacunosus* | 0 | 0 | 10.7 | 16.4 |
| *L. obtusus* | *L. venezuelensis* | 0.4 | 0.2 | 1.0 | 1.1 |
| *L. violaceus* | *L. cinerascens* | 0 | 0 | 5.7 | 6.3 |
| *L. rosmarinifolius* | <u>*L. cyparissias*</u> | 0 | 0 | 21.8 | 25.3 |
| *L. floribundus* | <u>*L. farallonensis*</u> | 0 | 0 | 42.0 | 46.0 |
| *L. floribundus* | <u>*L. perijaensis*</u> | 0 | 0 | 42.5 | 46.6 |
| *L. rhomboidalis* | *L. apiculatus* | 0 | 0 | 1.7 | 1.5 |
| *L. alveolatus* | *L. costaricensis* | 0 | 0 | 15.2 | 8.1 |
| *L. rhododendroides* | *L. schultzii* | 0 | 0 | 4.0 | 5.2 |
| *L. tenuifolius* | <u>*L. ellipticus*</u> | 0 | 0 | 17.1 | 20.1 |
| *L. oblongifolius* | *L. mutiscuanus* | 0 | 0 | 3.0 | 4.0 |
| *L. sp. nov. ANT* | *L. antioquensis* | 0 | 0 | 1.0 | 1.0 |
| *L. huertasii* | <u>*L. julianii*</u> | 0 | 0 | 7.0 | 7.0 |
| *L. eriophorus* | <u>*L. chrysotrichus*</u> | 0 | 0 | 7.0 | 9.0 |
| *L. colombianus* | *L. glutinosus* | 0.2 | 0.2 | 2.6 | 2.6 |
| *L. romeroi* | <u>*L. saxatilis*</u> | 1.0 | 0 | 3.0 | 1.5 |

### EVALUATING NICHE CONSERVATISM

Our results suggest that there is a strong signal of niche conservatism in the leaf area of *Linochilus*. Graphing the boxplots of leaf areas per species in front of the phylogeny reveals a pattern in which closely related species tend to have similar leaf areas, likely occupying similar niches (Fig. 4). These observations are supported by the Pagel’s (1999) lambda (λ) of 0.98 calculated with leaf area data on the phylogeny, P= 2.8e-09 against the null hypothesis of λ=0. This Pagel’s lambda of almost one suggests that the evolution of the leaf area is highly dependent on the phylogeny, possessing a strong phylogenetic signal. The phylogenetic signal found, builds on the sister species comparisons, where only a small fraction of pairs (13.3%) presents statistically significantly different leaf areas (Table 2).

**FIGURE 4.**
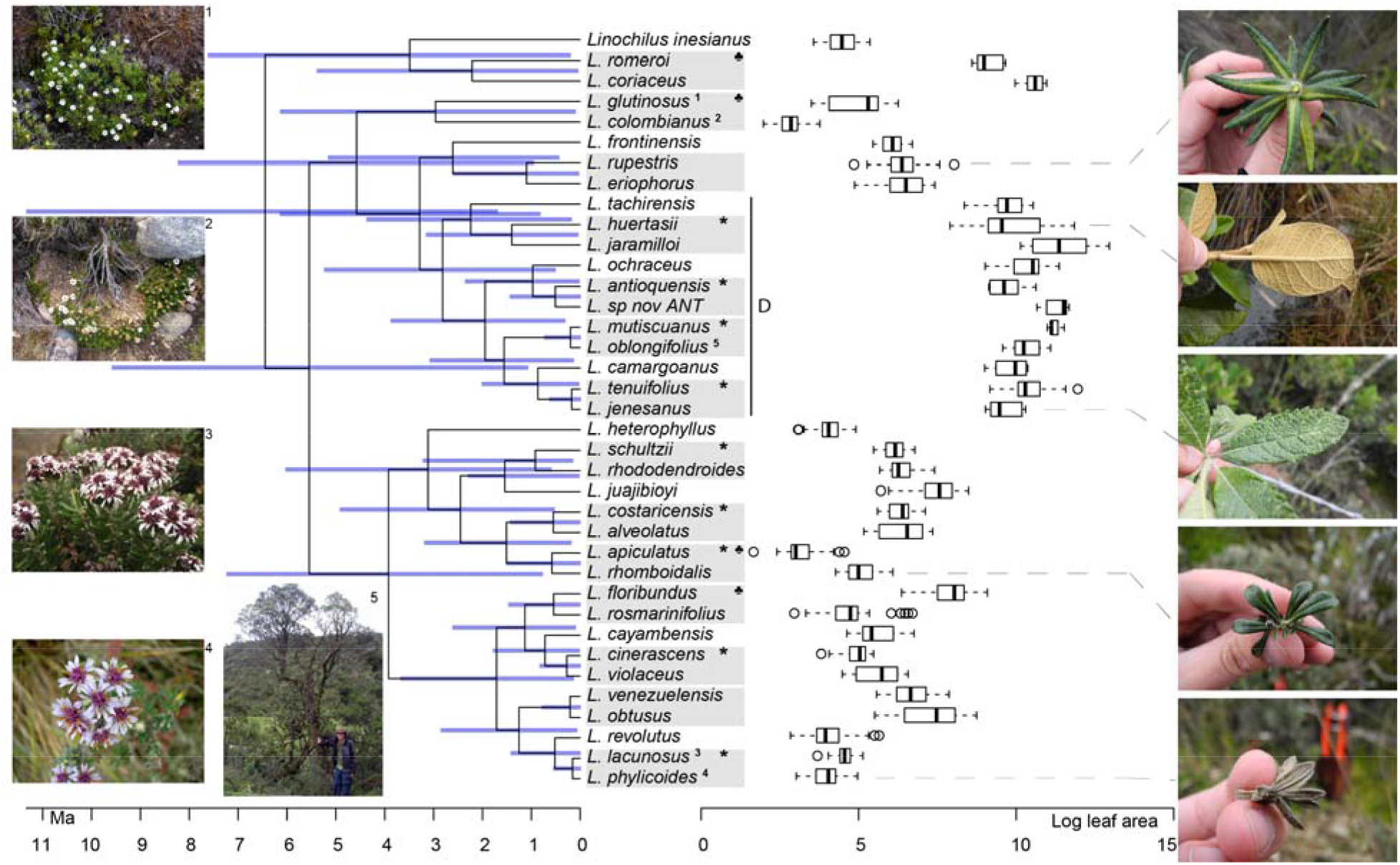
*Linochilus* phylogeny (Vargas et al. 2017) with boxplots for leaf area. Blue bars indicate confidence intervals for node ages. Photos on the left correspond to the superscript numeration on the tips of the phylogeny; photos 1 and 2 correspond to sympatric sister species with different leaf areas, while photos 3 and 4 correspond to allopatric sister species with similar leaf areas. Photos on the right correspond to the species indicated by the gray dashed line. The *Denticulata* clade is indicated by the letter D. Pairs are indicated by gray boxes, * : allopatric, ♣ : different leaf areas.

### BIOGEOGRAPHIC ANALYSES

The best scoring model in the biogeographic reconstruction for *Linochilus* was the BAYAREALIKE+J with an AICc of 226.78 followed by DEC+J with 232.62 (Table 4). The BAYAREALIKE+J reconstruction (Fig. 5) shows that the Colombian Eastern Cordillera played a major role in the diversification of *Linochilus*. This area, which contains the most species of the genus, was shown to be the ancestral range for most *Linochilus* ancestors (61%). The ancestral range for the node representing the ancestor for all *Linochilus* species is inconclusive, with two areas sharing aproximately two thirds of the relative probability: the Eastern Cordillera and the Northern Colombian Páramos (Sierra Nevada de Santa Marta + the Serranía del Perijá). The reconstruction shows that the *Denticulata* clade (clade D, Fig. 5), which downslope colonized the upper limit of the cloud forest from the páramo, originated in the Eastern Colombian Cordillera. The biogeographic analysis also shows that approximately two thirds of the species sampled are restricted to a single paramo complex (27 spp., 71%), while the other third is found in two or more complexes (11 spp. 29%). These biogeographic results evidence high endemism in the genus where most species are restricted to a few paramo islands, reinforcing the importance of geographic isolation in the diversification of the genus.

**FIGURE 5.**
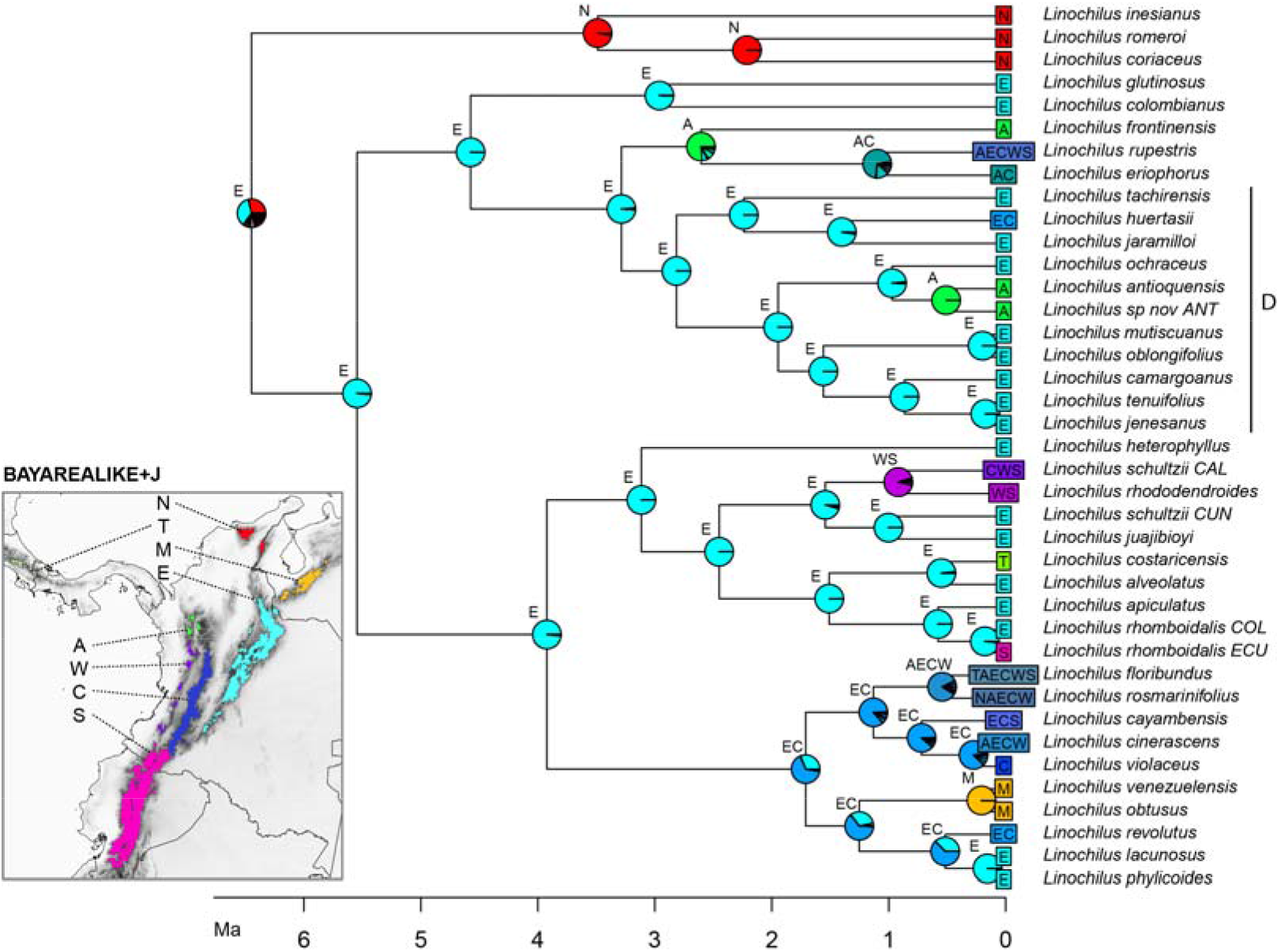
BAYAREALIKE+J biogeographical ancestral reconstruction based on the *Linochilus* phylogeny of Vargas et al. (2017) with percent probabilities of the different ancestral areas as pie charts. Letters indicate biogeographic areas considered in the analysis. N: northern páramos. T: Talamanca. M: Mérida. E: Eastern Cordillera. A: Antioquia’s páramos. W: Western Cordillera. C: Central Cordillera. S: southern páramos. Letters above each pie charts indicate the most probable area or area combination for that node.

**TABLE 4.** Comparison of the different biogeographic models for ancestral range inference evaluated by BioGeoBEARS in the phylogeny of *Linochilus* (Vargas et al. 2017).

| Model | LnL | Num. params | AIC |
| --- | --- | --- | --- |
| DEC | -115.09205 | 2 | 234.184098 |
| DEC+J | -113.31082 | 3 | 232.621645 |
| DIVALIKE | -115.63941 | 2 | 235.278824 |
| DIVALIKE+J | -115.64036 | 3 | 237.280724 |
| BAYAREALIKE | -121.933 | 2 | 247.865999 |
| BAYAREALIKE+J | -110.39081 | 3 | 226.78162 |

## DISCUSSION

In this study, we developed a framework to quantify the relative contributions of geographic isolation and parapatric ecological divergence in recent speciation events. We incorporated phylogenetics, geographic distributions, and a morpho-ecological trait. Our framework was applied to *Linochilus*, a genus of plants restricted to the páramo, which is the most species-rich tropical montane ecosystem. Because of the island-like distribution of the páramo and the within-páramo elevation gradient (Fig 2.), it has been suggested that allopatric speciation and parapatric ecological divergence are the main drivers of speciation in the high Andes (van der Hammen and Cleef 1986, Hughes and Atchison 2015). The application of our approach to *Linochilus* revealed that most recent speciation events are allopatric (Table 1), with most sister species presenting allopatry and similar leaf areas. The signal of allopatric speciation between sister species is robust independently of the method used to estimate allopatry/sympatry. 64.3% sisters are allopatric using presence/absence in páramo islands and the sister species inferred from Vargas et al. (2017). 80% sisters are allopatric when missing sampling is accounted for. Allopatry is inferred similarly (relative to presence/absence in paramo islands) for most sisters when using grids of 0.1 and 0.05 decimal degrees (Tables 1–3). Likewise, the contribution of parapatric ecological divergence to recent speciation events is low (< 25%) in both sets of results, phylogeny: 21.4% and phylogeny supplemented with missing sampling: 6.6%.

We believe that the most reliable estimate for calculating the relative contribution of allopatric speciation vs. parapatric ecological divergence is the phylogeny supplemented with missing sampling on the island framework results (second row, Table 1; Table 2). These results agree with the 0.1 grid results, are corrected for missing sampling in the phylogeny, and are more conservative at estimating allopatric pairs (relative to using a 0.05 degree grid). Therefore, *we will focus our discussion on the species pairs derived from the phylogeny and supplemented for missing sampling* (Tables 2–3).

### ALLOPATRIC VS. PARAPATRIC ECOLOGICAL SPECIATION IN THE PÁRAMO

Our results suggest that most recent speciation events (12, 80%) in *Linochilus* are driven by allopatric speciation, while only a few are driven by parapatric ecological divergence events (1, 6.6%) (Table 2, Fig. 3). Out of the 12 allopatric pairs, only one shows a signal of subsequent ecological divergence. Inconclusive divergence in pairs (2, 13.3%) was represented by a sympatric pair with similar leaf areas, and a sympatric pair with no leaf data. Independently of range overlap, only two pairs haven significantly different leaf areas. This lack of ecological divergence suggests a strong signal of niche conservatism in *Linochilus*, as does a Pagel’s λ of 0.98 (a value closer to one indicates strong phylogenetic signal in the evolution of leaf area). In the context of the páramo flora, our results are consistent with the hypothesis that páramos are island-like in promoting allopatric speciation. Our results suggest that allopatric speciation alone can explain most of the recent and rapid speciation found in other páramo genera where most species are restricted to one or few páramo islands: *Bartsia, Espeletia, Escallonia, Hypericum, Jamesonia-Eriosorus, Lachemilla*, and *Lupinus*, (Drummond et al. 2012; Zapata 2013; Sánchez-Baracaldo and Thomas 2014; Nürk et al. 2015; Uribe-Convers and Tank 2015; Diazgranados and Barber 2017; Contreras-Ortiz et al. 2018; Morales-Briones et al. 2018ab).

Although our results suggest a minor role of parapatric ecological divergence (6.6%) (Table 1), our analysis focused on sister species that diverged recently and does not consider more ancient divergent events. However, when we look at the distribution of leaf area on the phylogeny, we observe that taxa in clade D *(Linochilus* series *Denticulata)* have significantly larger leaves suited for dwelling in the upper zone of Andean forest (Fig. 3): Wilcoxon P < 2.2e-16 for both *L*. ser. *Denticulata* vs. its sister clade, and *L*. ser. *Denticulata* vs. *Linochilus’*s most species-rich clade (the clade originating with the most common ancestor of *L. heterophyllus* and *L. phylicoides*). The significantly larger leaves in *L*. ser. *Denticulata* suggests that an event of ecological shift, from microphyllous to macrophyllous leaves, took place ca. 3 Ma near the ancestor of the *Denticulata* clade leading to the evolution of at least 20 species (32% out of all species in *Linochilus*). Based on the larger leaf area found in the *Denticulata* clade, which allows *Linochilus* species to be competitive in the upper zone of the cloud forest, we hypothesize that this downslope colonization event by the *Denticulata* clade is a case ecological divergence.

Regardless of the reason for ecological divergence (after allopatric speciation or during parapatric ecological divergence), ecological divergence may *boost* allopatric speciation by allowing a lineage to colonize a new niche and rapidly speciate in it by means of allopatric speciation in an island-like system. In the specific case of *Linochilus*, the evolution of larger leaves, which happened in one single event ca. 3 Ma, allowed a lineage to colonize a lower vegetational belt and speciate. A similar pattern is found in Andean *Senecio*, where monophyletic forest and páramo clades have been documented to speciate in parallel (Dušková et al. 2017). In the context of adaptive radiations, we propose that ecological speciation happens fewer times when compared with allopatric speciation, but its effect could be significant at geological time scales; in other words, while allopatric speciation can explain most of the recent speciation, ecological divergence explains the origin of a single (or few) key adaptation(s) facilitating the colonization of new niches. *Espeletia*, whose crown origin is estimated at 2.5 Ma, shows a peak in morphological differentiation relatively deep in its phylogeny at 1.5 Ma (Pouchon et al. 2018) in addition to have a strong signal of allopatric speciation (Diazgranados and Barber 2017). Other potential diversification *boosters* are genome duplication events (Morales-Briones et al. 2018a) and pollination shifts (Lagomarsino et al. 2016).

### ALTERNATIVE HYPOTHESES AND FURTHER CONSIDERATIONS

Our framework is unable to distinguish between vicariant and peripatric speciation. Testing which kind of allopatric speciation occurs is challenging in island systems—typically peripatric speciation predicts that the founder species will be distributed in a significantly smaller area than the source lineage (Coyne and Orr 2004; Anacker and Strauss 2014; Grossenbancher et al. 2014; Skeels and Cardillo 2019), but geographically restricted patches (e.g. páramos) can confound testing for significant differences in the distributions. Additionally, páramo islands have shifted their elevational distribution due to Pleistocene climate fluctuations, connecting páramo islands during glaciation periods and disconnecting them during interglacial periods (Simpson 1974, 1975; Flantua et al. 2019). This “flickering connectivity” could be a major driver of geographic isolation in páramo plants, especially for taxa with low seed dispersal ability (i.g. *Espeletia* complex with ca. 100 spp.). In *Linochilus*, which has a fruit that easily disperses with wind, we find that 10 sisters (66.6%) present range asymmetry in which the widespread-sister range is > 3 times larger than the small-ranged sister, suggesting peripatric speciation as a lead player in the diversification of *Linochilus;* we advise to take this result with caution for the aforementioned reasons. Finally, the best fit of the BAYAREALIKE+J as the best model for the biogeographic analysis in *Linochilus* supports the idea that founder speciation events (peripatric speciation), modeled by the J parameter, are important for the speciation of *Linochilus* in the páramos.

Our sister-species framework assumes that speciation is a bifurcating process in which every speciation event produces two reciprocal monophyletic species. In the context of vicariant speciation in the páramo, a glacial-interglacial event could result in the fragmentation of one previously continually distributed population, into more than two daughter proto-species; the complex topography of the Colombian Eastern Cordillera provides a probable location for this process to occur (Fig 2, Fig 7-10 in van der Hammen and Cleef 1986). Hypothetically, parapatric and peripatric speciation can also violate a bifurcating speciation model because a widely distributed population could be the source for multiple independent parapatric and or peripatric divergent events (e.g. upslope colonization and adaptation in different mountains, multiple dispersal events, Dexter et al. 2017).

How does the result of allopatric speciation driving diversification align with recent finds of pervasive hybridization-introgression in *Linochilus* (Vargas et al. 2017) and other páramo plants (Morales-Briones et al. 2018a; Nevado et al. 2018; Pouchon et al. 2018)? High gene flow indices can be explained by three non-exclusive scenarios. First, under allopatric speciation, recent geographically isolated populations (proto-species) should be reproductively compatible for the time that it takes to reproductive barriers to develop. During that time window, gene transfer between these proto-species is likely to occur leaving a signal of introgression among sister species. Second, it is possible that, given geographic isolation, the biological pressure to enforce reproductive barriers is relaxed, making geographically isolated proto-species reproductively compatible for a long period of time; this would allow hybridization post-morphological differentiation—that is, hybridization after the time point when they are morphologically different and therefore considered by taxonomists as different species. Third, considering the small geographical ranges and likely low population numbers of many *Linochilus* species, it is possible that introgression has a positive effect by increasing genetic variability and counteracting genetic drift —making gene flow adaptive. Testing these hypotheses with phylogeographic approaches would help to elucidate the role of hybridization in the evolution of páramo plants.

Finally, it has been suggested that hybridization can cause speciation because it has the potential of producing genotypes pre-adapted to un-exploited niches (Anderson and Stebbins 1954; Seehausen 2004); this hypothesis predicts a signal of ecological divergence between the hybrid and its parental species. Taking into account that we only observed two cases of parapatric ecological divergence in sister species, our results suggest that hybrid speciation might not play a major role in the recent diversification history of *Linochilus*.

### SPATIOTEMPORAL HISTORY OF *LINOCHILUS* IN THE CONTEXT OF THE PÁRAMO

Our biogeographic reconstruction of *Linochilus* shows that the genus originated in the Northern Andes 6.46 Ma, predating the estimated origin of the páramo 2–4 Ma (van der Hammen and Cleef 1986; Gregory-Wodzicki 2000). Despite the fact that *Linochilus’*s inferred age 95% confidence interval of 1.71–11.37 can accommodate such incongruence, other primarily páramo genera like *Arcytophyllum, Brunfelsia, Jamesonia+Eriosorus, Lysipomia, Valeriana*, and *Vasconcellea* also show ages older than 4 Ma (Luebert and Weigend 2014). An explanation for this early origin could be that ancestors of these lineages inhabited the summits of middle elevation mountains (<2000 m) extant at that time. Mid-elevation tropical mountains often have open and semi-dry areas at upper elevations, which are somewhat physiologically similar to the páramo. These physiologically dry patches are caused by well-drained soils and strong winds similar to the contemporary *campos de altitude* and *campus rupestres* in Brazil (Safford 1999; Alves et al. 2014). It is possible that middle elevation mountaintops provided an early habitat for *Linochilus* ancestors before higher elevations were available at 4 Ma. A second alternative is that páramos were available before 2–4 Ma as suggested by Ehlers and Poulsen (2009). A third scenario is that *Linochilus* originated in the Sierra Nevada de Santa Marta (SNSM), a mountain range located in northern Colombia which is now separated from the main Andes Cordilleras. SNSM’s paleoelevation remains largely unstudied (Villagómez et al. 2011), but our biogeographic analysis provides some evidence for the last hypothesis because the *Linochilus* species endemic to the SNSM (*L*. *coriaceus, L. inesianus*, and *P. romeroi)* comprise a clade that that is sister to the rest of *Linochilus*, making the SNSM the second most likely area of origin for the genus after the Eastern Cordillera (Fig. 5).

Our biogeographic analysis also suggests that the Eastern Cordillera of Colombia played a major role in the diversification of the genus given the many extant and ancestral species whose distributional range include this area. The Eastern Cordillera contains the most páramo land area with discrete patches (Londoño et al. 2014) making it ideal for autochthonous allopatric speciation. Our reconstruction also indicates that the Eastern Cordillera was the source for the colonization of three other mountain ranges: the Colombian Central Cordillera, the Colombian Western Cordillera, and the Talamanca Cordillera in Central America.

Achenes of *Linochilus* are small and a have a pappus that allows for long-distance dispersal of their seeds by wind (Cuatrecasas 1969). Collectively, *Linochilus* is found on almost every páramo island, with the exception of the southernmost páramos, south of the Girón-Paute valley in Ecuador, which is a biogeographic barrier for numerous Andean genera (Jørgensen et al. 1995). Specific examples of long-distance dispersal in *Linochilus* are shown by the two species reported at the westernmost páramos in Costa Rica. *L*. *costaricensis* is endemic to Costa Rica and is probably a direct descendant of *L. alveolatus*, which inhabits the Colombian Eastern Cordillera. The second Costa Rican species, *L. floribundus*, is also reported for Colombia and Ecuador (Vargas 2011; 2018). The aforementioned examples of long-distance dispersal build on previous results about the potential dominant role of peripatric speciation in the genus and in other high montane taxa with seeds or fruits adapted to wind dispersal.

## CONCLUSIONS

Our comparative framework that incorporates phylogenetics, geographical distributions, and morpho-ecological characters unveiled a high signal of allopatric speciation, supporting it as the process driving most of the recent speciation events in *Linochilus*. The island-like distribution of the páramo is likely a primary factor of autochthonous allopatric speciation via geographic isolation, explaining the particularly high accumulation of plant species in the páramo (Simpson and Todzia 1990) and their high speciation rates (Madriñán et al. 2013). Despite the comparatively small role of parapatric ecological speciation identified in recent *Linochilus* sister taxa, we propose that ecological divergence has a role that is infrequent but potentially powerful in island-like systems. When ecological divergence does occur, it allows a lineage to colonize a new niche and then rapidly speciate by means of allopatric speciation among islands. Ecological divergence events that *boost* diversification are thus expected to be detectable at deeper geological time scales (> 1 Ma). We conclude that geographic isolation and parapatric ecological divergence are positively synergistic processes in the history of the diversification of the paramo flora, contributing significantly to the global latitudinal species gradient.

## ACKNOWLEDGEMENTS

We thank Stefani Brandt, Julia Harencar, Shelley Sianta, Kathleen Kay, and Dena Grossenbacher for helpful comments and discussion during the writing of this manuscript. This study would not have been possible without the scientific collections housed at Herbario Nacional Colombiano (COL), the Billie Turner Plant Resources Center (TEX), and the United States National Herbarium at the Smithsonian Institution (US). Financial support was provided by The University of Texas at Austin (Plant Biology Program Awards, the C. L. Lundell Chair of Systematic Botany, The Linda Escobar Award), the Garden Club of America (2012 Award in Tropical Botany), and the Smithsonian Institution (Cuatrecasas Award 2006).

## CONFLICT OF INTEREST

We declare no competing interest.

## Notes

### Competing Interest Statement

The authors have declared no competing interest.

### Summary of Updates

This version of the manuscript has been revised to update changes in taxonomic nomenclature, edit part of the text, and add maps to the manuscript.

