## Appendix for "Allopatric speciation is more prevalent than parapatric ecological divergence in a recent high-Andean diversification (Asteraceae: *Linochilus*)"

*­Evolution*

SUPPORTING INFORMATION

### Allopatric speciation is more prevalent than parapatric ecological divergence in tropical montane systems (Asteraceae: *Linochilus*)

**APPENDIX S1** Occurrences used for the speciation and historical biogeographical analyses.

| **Name** | **Identifier** | **Latitud** | **Longitud** |
| --- | --- | --- | --- |
| Linochilus alveolatus | 27015 | 3.18333 | -76.21667 |
| Linochilus alveolatus | 214 | 3.75 | -74.42 |
| Linochilus alveolatus | 2605 | 3.75 | -74.42 |
| Linochilus alveolatus | 7764 | 3.75 | -74.42 |
| Linochilus alveolatus | 7904 | 3.75 | -74.42 |
| Linochilus alveolatus | 2354 | 3.75 | -74.42 |
| Linochilus alveolatus | 1667 | 3.75 | -74.42 |
| Linochilus alveolatus | 1706 | 3.75 | -74.42 |
| Linochilus alveolatus | 5765 | 3.75 | -74.42 |
| Linochilus alveolatus | 6891 | 3.75 | -74.42 |
| Linochilus alveolatus | 360 | 3.85 | -74.05 |
| Linochilus alveolatus | 382 | 3.85 | -74.05 |
| Linochilus alveolatus | 383 | 3.85 | -74.05 |
| Linochilus alveolatus | 2859 | 4.07444 | -74.24833 |
| Linochilus alveolatus | 27023 | 4.09805556 | -74.25 |
| Linochilus alveolatus | 25871 | 4.27333 | -74.19944 |
| Linochilus alveolatus | 25778 | 4.28389 | -74.2084 |
| Linochilus alveolatus | 26496 | 4.28389 | -74.2084 |
| Linochilus alveolatus | 3725 | 4.47306 | -74.11611 |
| Linochilus alveolatus | 7001 | 4.55 | -74.006 |
| Linochilus alveolatus | 2 | 4.84250212 | -73.814522 |
| Linochilus alveolatus | 9989 | 4.93333 | -73.85 |
| Linochilus alveolatus | 6509 | 5.1267 | -74.0011 |
| Linochilus alveolatus | 806 | 5.185 | -74.018 |
| Linochilus alveolatus | 2730A | 6.082434 | -72.49749 |
| Linochilus alveolatus | 1758 | 6.28662014 | -72.511864 |
| Linochilus alveolatus | 8781 | 6.361927 | -72.334908 |
| Linochilus alveolatus | 5564 | 6.370335 | -72.331151 |
| Linochilus alveolatus | 8651 | 6.383104 | -72.310565 |
| Linochilus alveolatus | 1582 | 6.385995 | -72.344335 |
| Linochilus alveolatus | 9990 | 6.402217 | -72.36426 |
| Linochilus alveolatus | 27819 | 6.405436 | -72.372883 |
| Linochilus anactinotus | 1911 | 10.811 | -73.941 |
| Linochilus anactinotus | 24557 | 10.819 | -73.639 |
| Linochilus anactinotus | 6925 | 10.85 | -73.68 |
| Linochilus anactinotus | 6940 | 10.85 | -73.68 |
| Linochilus anactinotus | 6967 | 10.85 | -73.68 |
| Linochilus anactinotus | 6972 | 10.85 | -73.68 |
| Linochilus anactinotus | 6974 | 10.85 | -73.68 |
| Linochilus anactinotus | 390 | 11.038 | -73.619 |
| Linochilus anactinotus | 955 | 11.038 | -73.619 |
| Linochilus anactinotus | 987 | 11.038 | -73.619 |
| Linochilus andinus | 3245 | 5.54972 | -75.98583 |
| Linochilus andinus | 3284 | 5.54972 | -75.98583 |
| Linochilus antioquensis | 5292 | 6.26666667 | -75.683333 |
| Linochilus antioquensis | 5342 | 6.26666667 | -75.683333 |
| Linochilus antioquensis | 24226 | 6.27806 | -75.68333 |
| Linochilus apiculatus | 19571 | 7.223 | -72.917 |
| Linochilus apiculatus | 17499 | 7.3312 | -72.8896 |
| Linochilus apiculatus | 18430 | 7.363 | -72.929 |
| Linochilus apiculatus | 1227 | 7.38 | -72.87 |
| Linochilus bicolor | 26367 | 0.81667 | -77.4 |
| Linochilus bicolor | 26398 | 0.81667 | -77.4 |
| Linochilus bicolor | 26413 | 0.81667 | -77.4 |
| Linochilus bicolor | 11932 | 1.17 | -77.2 |
| Linochilus bicolor | 5344 | 1.17 | -77.2 |
| Linochilus bicolor | 5759 | 1.1884 | -76.9954 |
| Linochilus bicolor | 11706 | 1.18841 | -76.9159 |
| Linochilus bicolor | 222 | 1.19711 | -76.92242 |
| Linochilus bicolor | 7590 | 1.2075 | -76.979 |
| Linochilus bicolor | 1060 | 1.2217 | -77.2931 |
| Linochilus bicolor | 18567 | 1.3151 | -77.2545 |
| Linochilus bicolor | 1895 | 1.46176 | -76.93162 |
| Linochilus bicolor | 8679 | 2.3341 | -75.4873 |
| Linochilus bicolor | 14925 | 2.3955 | -75.337 |
| Linochilus bicolor | 2587 | 2.765 | -76.045 |
| Linochilus bicolor | 1465 | 2.93327 | -75.7658 |
| Linochilus bicolor | 2513 | 2.9686 | -76.1416 |
| Linochilus bicolor | 2386 | 2.98881 | -76.16885 |
| Linochilus bicolor | 11932 | 3.06441 | -76.15259 |
| Linochilus bicolor | 17920 | 3.37595 | -76.68905 |
| Linochilus bicolor | 21832 | 3.37595 | -76.68905 |
| Linochilus bicolor | 3 | 3.37595 | -76.68905 |
| Linochilus bicolor | 59 | 3.59951 | -76.49311 |
| Linochilus bicolor | 462 | 3.6736 | -75.9935 |
| Linochilus bicolor | 4351 | 3.768337 | -75.96576 |
| Linochilus bicolor | 7488 | 3.97090006 | -75.661797 |
| Linochilus bicolor | 3027 | 4.19721 | -75.74971 |
| Linochilus bicolor | 66-B | 4.20355 | -76.41281 |
| Linochilus bicolor | 8164 | 4.30249977 | -75.606102 |
| Linochilus bicolor | 8705 | 4.30249977 | -75.606102 |
| Linochilus bicolor | WV8168 | 4.30249977 | -75.606102 |
| Linochilus bicolor | 27679 | 4.45539999 | -75.573998 |
| Linochilus bicolor | 2126 | 4.7203 | -75.2517 |
| Linochilus bicolor | 27704 | 4.99905 | -75.00327 |
| Linochilus bicolor | 6000 | 4.99905 | -75.00327 |
| Linochilus bicolor | 2066 | 5.04336 | -75.36434 |
| Linochilus camargoanus | 2984 | 5.73171997 | -73.441429 |
| Linochilus camargoanus | 6309 | 5.75 | -73.43333 |
| Linochilus camargoanus | 6309 | 5.75 | -73.43333 |
| Linochilus camargoanus | 12110 | 5.77892 | -72.95843 |
| Linochilus camargoanus | 5224 | 5.8778 | -73.12951 |
| Linochilus camargoanus | 2635 | 6.10731 | -73.19771 |
| Linochilus cayambensis | 28644 | 1.13 | -77.12 |
| Linochilus cayambensis | 11782 | 1.14499 | -77.10122 |
| Linochilus cayambensis | 155 | 1.22 | -77.13 |
| Linochilus cayambensis | 1505 | 1.49998 | -76.90001 |
| Linochilus cayambensis | 1985 | 1.54719 | -76.91395 |
| Linochilus cayambensis | 2971 | 1.92 | -76.6 |
| Linochilus cayambensis | 3242 | 1.92 | -76.6 |
| Linochilus cayambensis | 4094 | 1.92 | -76.6 |
| Linochilus cayambensis | 872 | 2.41996 | -76.38476 |
| Linochilus cayambensis | 2737 | 3.86047 | -74.34874 |
| Linochilus chrysotrichus | 263 | 4.30249977 | -75.606102 |
| Linochilus chrysotrichus | 189 | 4.30999994 | -75.610001 |
| Linochilus chrysotrichus | 371 | 4.30999994 | -75.610001 |
| Linochilus cinerascens | 17851 | 3.37595 | -76.68905 |
| Linochilus cinerascens | 1839 | 5.05148 | -76.09248 |
| Linochilus cinerascens | 1798 | 5.13913 | -76.18014 |
| Linochilus cinerascens | 2428 | 5.163 | -76.095 |
| Linochilus cinerascens | 1610 | 5.16353 | -76.09478 |
| Linochilus cinerascens | 1741 | 5.16353 | -76.09478 |
| Linochilus cinerascens | 273 | 6.47 | -76.1 |
| Linochilus cinerascens | 2288 | 6.47 | -76.1 |
| Linochilus cinerascens | 2340 | 6.47 | -76.1 |
| Linochilus cinerascens | 13176 | 6.5 | -76.11667 |
| Linochilus cinerascens | 540 | 1.5 | -76.5 |
| Linochilus cinerascens | 598 | 1.5 | -76.5 |
| Linochilus cinerascens | 659 | 2.20417 | -76.49639 |
| Linochilus cinerascens | 513 | 2.24028 | -76.17028 |
| Linochilus cinerascens | 26282 | 2.32 | -76.31 |
| Linochilus cinerascens | 244 | 2.32 | -76.31 |
| Linochilus cinerascens | 134 | 2.32 | -76.31 |
| Linochilus cinerascens | 14602 | 2.4 | -76.38 |
| Linochilus cinerascens | 38572 | 2.4 | -76.38 |
| Linochilus cinerascens | 26415 | 2.49 | -76.29 |
| Linochilus cinerascens | 27371 | 2.83 | -76.2 |
| Linochilus cinerascens | 439 | 2.83 | -76.2 |
| Linochilus cinerascens | 2564 | 2.933 | -76.174 |
| Linochilus cinerascens | 1470 | 3.0193 | -76.0077 |
| Linochilus cinerascens | 1865 | 3.73 | -75.93 |
| Linochilus cinerascens | 27547 | 3.75 | -75.96667 |
| Linochilus cinerascens | 18796 | 4.19241 | -76.4595 |
| Linochilus cinerascens | 8669 | 8.45 | -73.41 |
| Linochilus colombianus | 3565 | 5.85784 | -73.15854 |
| Linochilus colombianus | 6993 | 5.98 | -73.08 |
| Linochilus colombianus | 7259 | 5.98 | -73.08 |
| Linochilus colombianus | 2376 | 5.98 | -73.08 |
| Linochilus colombianus | 924 | 6.27 | -72.87 |
| Linochilus colombianus | 1492 | 6.368452 | -72.333851 |
| Linochilus colombianus | 5620 | 6.380028 | -72.304029 |
| Linochilus colombianus | 5682 | 6.385979 | -72.327765 |
| Linochilus colombianus | 1549 | 6.549573 | -72.328398 |
| Linochilus colombianus | 1549 | 6.549573 | -72.328398 |
| Linochilus coriaceus | 4546 | 10.819 | -73.549 |
| Linochilus coriaceus | 7112 | 10.819 | -73.549 |
| Linochilus coriaceus | 7162 | 10.819 | -73.549 |
| Linochilus coriaceus | 6691 | 10.889 | -73.878 |
| Linochilus costaricensis | 1169 | 9.5553593 | -83.76378 |
| Linochilus costaricensis | 8022 | 9.5666093 | -83.749982 |
| Linochilus crassifolius | 7373 | 10.085 | -72.931 |
| Linochilus crassifolius | 25120 | 10.211 | -72.91 |
| Linochilus crassifolius | 11191 | 10.343 | -72.911 |
| Linochilus crassifolius | 10865 | 10.42738 | -72.89652 |
| Linochilus cyparissias | 6931 | 10.85 | -73.68 |
| Linochilus cyparissias | 24648 | 10.91 | -73.52 |
| Linochilus ellipticus | 12336 | 7.35 | -72.65 |
| Linochilus eriophorus | 3329 | 1.92 | -76.6 |
| Linochilus eriophorus | s.n. | 4.65992 | -75.33995 |
| Linochilus eriophorus | 8077 | 4.707 | -75.391 |
| Linochilus eriophorus | 1808 | 4.72 | -75.414 |
| Linochilus eriophorus | 5690 | 4.802385 | -75.378196 |
| Linochilus eriophorus | 6348 | 4.858 | -75.337 |
| Linochilus eriophorus | 5235 | 4.9 | -75.3 |
| Linochilus eriophorus | 2512 | 4.9 | -75.3 |
| Linochilus eriophorus | 9266 | 4.9 | -75.3 |
| Linochilus eriophorus | 3714 | 4.9 | -75.3 |
| Linochilus eriophorus | 782 | 4.9 | -75.3 |
| Linochilus eriophorus | 877 | 5.00008 | -75.35769 |
| Linochilus eriophorus | 8018 | 5.028 | -75.333 |
| Linochilus farallonensis | 17855 | 3.337 | -76.699 |
| Linochilus floribundus | 2243 | 0.7999 | -77.9166 |
| Linochilus floribundus | 12864 | 0.9546 | -77.8829 |
| Linochilus floribundus | 11822 | 1.0814 | -77.0638 |
| Linochilus floribundus | 970 | 1.083 | -77.15 |
| Linochilus floribundus | 12784 | 1.0874 | -77.6842 |
| Linochilus floribundus | 4479 | 1.13 | -77.12 |
| Linochilus floribundus | 994 | 1.159 | -77.24287 |
| Linochilus floribundus | 5004 | 1.2117 | -77.3633 |
| Linochilus floribundus | 7998 | 1.216 | -77.371 |
| Linochilus floribundus | 26939 | 1.2208 | -77.3574 |
| Linochilus floribundus | 3482 | 1.346 | -76.9102 |
| Linochilus floribundus | 5817 | 1.9 | -76.666 |
| Linochilus floribundus | 5875 | 1.9 | -76.666 |
| Linochilus floribundus | 3960 | 1.92 | -76.6 |
| Linochilus floribundus | 27382 | 2.216 | -76.467 |
| Linochilus floribundus | 27545 | 2.22 | -76.32 |
| Linochilus floribundus | 2699 | 2.3138 | -76.3952 |
| Linochilus floribundus | 26293 | 2.3592 | -76.3556 |
| Linochilus floribundus | 1186 | 3.47662 | -76.04785 |
| Linochilus floribundus | 868 | 3.7 | -76.1 |
| Linochilus floribundus | 2009 | 3.7155 | -76.1207 |
| Linochilus floribundus | 1188 | 3.7493 | -75.9464 |
| Linochilus floribundus |  | 3.8 | -76.08 |
| Linochilus floribundus | 483 | 3.8 | -76.08 |
| Linochilus floribundus | 4213 | 3.96484 | -75.91038 |
| Linochilus floribundus | 7512 | 3.97117 | -75.60107 |
| Linochilus floribundus | 7771 | 3.9879601 | -75.639107 |
| Linochilus floribundus | 20547 | 4.02892 | -75.81919 |
| Linochilus floribundus | 10239 | 4.1924 | -76.4594 |
| Linochilus floribundus | 3078 | 4.19721 | -75.74971 |
| Linochilus floribundus | 8704 | 4.30249977 | -75.606102 |
| Linochilus floribundus | 27672 | 4.45539999 | -75.573998 |
| Linochilus floribundus | sn. | 4.5798 | -74.022 |
| Linochilus floribundus | 134 | 4.6259 | -73.7442 |
| Linochilus floribundus | 2028 | 4.70087 | -75.32059 |
| Linochilus floribundus | 6189 | 4.73769999 | -75.309601 |
| Linochilus floribundus | 6288 | 4.7516 | -75.3809 |
| Linochilus floribundus | 403 | 4.78945017 | -73.822067 |
| Linochilus floribundus | 16 | 5.00686 | -75.49382 |
| Linochilus floribundus | 3692 | 5.066 | -74.036 |
| Linochilus floribundus | 26948 | 5.07 | -74.1 |
| Linochilus floribundus | 26952 | 5.07 | -74.1 |
| Linochilus floribundus | 936 | 5.0874 | -74.0531 |
| Linochilus floribundus | 9543 | 5.10777 | -74.0728 |
| Linochilus floribundus | 587 | 5.1258 | -75.35455 |
| Linochilus floribundus | 768 | 5.1258 | -75.35455 |
| Linochilus floribundus | 2030 | 5.14221 | -76.0929 |
| Linochilus floribundus | 802 | 5.2129 | -74.0284 |
| Linochilus floribundus | 816 | 5.2129 | -74.0284 |
| Linochilus floribundus | 10356 | 6.050244 | -72.900109 |
| Linochilus floribundus | 1950 | 6.0674 | -72.9441 |
| Linochilus floribundus | 591 | 6.44717 | -76.08432 |
| Linochilus floribundus | 2420 | 6.45556021 | -76.116669 |
| Linochilus floribundus | 1397 | 6.47 | -76.1 |
| Linochilus floribundus | 9961 | 6.95 | -72.68 |
| Linochilus floribundus | 1126 | 2.38815 | -76.2755 |
| Linochilus floribundus | 2884 | 4.64675 | -75.35626 |
| Linochilus fosbergii | 2620 | 3.7248 | -74.01 |
| Linochilus fosbergii | 2722 | 3.7248 | -74.01 |
| Linochilus fosbergii | 8332 | 3.75 | -74.42 |
| Linochilus fosbergii | 7856 | 3.931 | -74.0749 |
| Linochilus fosbergii | 2636 | 4.0169 | -73.9944 |
| Linochilus fosbergii | 20912 | 3.86667 | -74.24 |
| Linochilus frontinensis | 2358 | 6.47 | -76.1 |
| Linochilus frontinensis | 13135 | 6.5 | -76.116669 |
| Linochilus frontinensis | 4424 | 6.5 | -76.116669 |
| Linochilus glutinosus | 4533 | 7.18 | -72.88 |
| Linochilus glutinosus | 79 | 7.1952 | -72.8615 |
| Linochilus glutinosus | 1226 | 7.1952 | -72.8615 |
| Linochilus glutinosus | 2 | 7.1952 | -72.8615 |
| Linochilus glutinosus | 18440 | 7.3767 | -72.8533 |
| Linochilus glutinosus | 19748 | 7.8472 | -73.2253 |
| Linochilus glutinosus | 20730 | 7.8472 | -73.2253 |
| Linochilus glutinosus | 4487 | 5.88 | -72.62 |
| Linochilus glutinosus | 4592 | 5.88 | -72.62 |
| Linochilus glutinosus | 6113 | 5.98 | -73.08 |
| Linochilus glutinosus | 9195 | 6.5169 | -72.1841 |
| Linochilus glutinosus | 2490 | 6.7299 | -72.6299 |
| Linochilus glutinosus | 14.861 | 2.312 | -75.413 |
| Linochilus glutinosus | 43 | 6.8618 | -72.7325 |
| Linochilus glutinosus | 3900 | 6.96 | -72.68 |
| Linochilus glutinosus | 10393 | 6.96 | -72.68 |
| Linochilus glutinosus | 20799 | 7.0287 | -72.717 |
| Linochilus glutinosus | 17487 | 7.28257 | -72.87837 |
| Linochilus glutinosus | 145 | 7.41907 | -72.34351 |
| Linochilus grantii | 4506 | 9.8354 | -72.9447 |
| Linochilus grantii | 25153 | 10.3733 | -72.8966 |
| Linochilus grantii | 25152 | 10.39938 | -72.91173 |
| Linochilus grantii | 7301 | 10.39938 | -72.91173 |
| Linochilus grantii | 10791 | 10.42738 | -72.89652 |
| Linochilus heterophyllus | 2736 | 3.7248 | -74.01 |
| Linochilus heterophyllus | 1109 | 3.75 | -74.42 |
| Linochilus heterophyllus | 1232 | 3.75 | -74.42 |
| Linochilus heterophyllus | 8306 | 3.75 | -74.42 |
| Linochilus heterophyllus | 8356 | 3.75 | -74.42 |
| Linochilus heterophyllus | 16170 | 4.5273 | -74.1845 |
| Linochilus heterophyllus | 16186 | 4.5273 | -74.1845 |
| Linochilus heterophyllus | 10457 | 4.57 | -74.03 |
| Linochilus heterophyllus | 1264 | 4.57 | -74.03 |
| Linochilus heterophyllus | 614 | 4.6182 | -73.7471 |
| Linochilus heterophyllus | 6722 | 4.7695 | -73.7508 |
| Linochilus heterophyllus | 304 | 4.7695 | -73.7508 |
| Linochilus heterophyllus | 781 | 4.79677916 | -73.814171 |
| Linochilus heterophyllus | sn. | 4.9 | -73.78 |
| Linochilus huertasii | 3258 | 1.92 | -76.6 |
| Linochilus huertasii | 533 | 2.1814 | -76.4307 |
| Linochilus huertasii | 2723 | 3.7248 | -74.01 |
| Linochilus huertasii | 2742 | 3.7248 | -74.01 |
| Linochilus huertasii | 5951 | 4.52 | -73.75 |
| Linochilus huertasii | 54 | 4.52925015 | -73.732224 |
| Linochilus huertasii | 9425 | 5.5 | -72.7498 |
| Linochilus huertasii | 28706 | 5.5329 | -72.7917 |
| Linochilus huertasii | 4756 | 6.1649 | -72.4167 |
| Linochilus huertasii | 9917 | 6.1649 | -72.4167 |
| Linochilus inesianus | 526 | 10.7605 | -73.6335 |
| Linochilus inesianus | 7127 | 10.8824 | -73.7994 |
| Linochilus inesianus | 7140 | 10.8824 | -73.7994 |
| Linochilus inesianus | 1382 | 10.9853 | -73.868 |
| Linochilus jaramilloi | 356 | 5.67954016 | -73.479874 |
| Linochilus jaramilloi | 28667 | 5.7671 | -73.426 |
| Linochilus jaramilloi | 28667 | 5.7671 | -73.426 |
| Linochilus jaramilloi | 20262 | 5.7671 | -73.426 |
| Linochilus jenesanus | 509 | 5.21361 | -73.5725 |
| Linochilus jenesanus | 14 | 5.379427 | -73.391853 |
| Linochilus jenesanus | 7162 | 5.42384 | -73.53025 |
| Linochilus jenesanus | 7163 | 5.42384 | -73.53025 |
| Linochilus juajibioyi | 4304 | 5.88 | -72.62 |
| Linochilus juajibioyi | 9185 | 6.3909 | -72.2074 |
| Linochilus juajibioyi | 10388 | 6.95 | -72.68 |
| Linochilus juajibioyi | 7584 | 3.95665 | -74.16973 |
| Linochilus juajibioyi | 8017 | 3.95665 | -74.16973 |
| Linochilus julianii | 98616 | 7.41306 | -72.407 |
| Linochilus lacunosus | 1443 | 6.368452 | -72.333851 |
| Linochilus lacunosus | 5659 | 6.394393 | -72.324749 |
| Linochilus lacunosus | 8766 | 6.477948 | -72.316125 |
| Linochilus lacunosus | 1069 | 7.41013 | -72.85549 |
| Linochilus leiocladus | 10531 | 5.1737 | -76.09354 |
| Linochilus longilobatus | 14850 | 3.91663 | -75.9333 |
| Linochilus longilobatus | 14856 | 3.91663 | -75.9333 |
| Linochilus longilobatus | 14856 | 3.91663 | -75.9333 |
| Linochilus micradenius | 1655 | 5.1651 | -76.0829 |
| Linochilus micradenius | 10533 | 5.16652 | -76.09333 |
| Linochilus mutiscuanus | 19708 | 7.33497 | -72.70074 |
| Linochilus nevadensis | 24440 | 10.7653 | -73.5278 |
| Linochilus nevadensis | 6937 | 10.85 | -73.68 |
| Linochilus nevadensis | 6 | 10.86534 | -73.7217 |
| Linochilus oblongifolius | 13463 | 7.27917 | -72.24639 |
| Linochilus oblongifolius | 1294 | 7.35 | -72.65 |
| Linochilus oblongifolius | 1273 | 7.38 | -72.57 |
| Linochilus oblongifolius | 10238 | 7.3942 | -72.6456 |
| Linochilus obtusus | 28393 | 8.14784467 | -71.907719 |
| Linochilus obtusus | 14726 | 8.17393819 | -71.846134 |
| Linochilus obtusus | 28801 | 8.33445074 | -71.279937 |
| Linochilus obtusus | 8778 | 8.55516481 | -71.084063 |
| Linochilus obtusus | 28606 | 8.75654882 | -70.806364 |
| Linochilus obtusus | 9246 | 9.03104647 | -70.581893 |
| Linochilus obtusus | 28062 | 9.04795135 | -70.86962 |
| Linochilus obtusus | 1123 | 9.51578366 | -70.108099 |
| Linochilus obtusus | 24 | 9.58724171 | -70.120773 |
| Linochilus ocanensis | 1199 | 8.06144 | -73.01432 |
| Linochilus ochraceus | 2900 | 4.0744 | -74.2483 |
| Linochilus ochraceus | 5175 | 4.4333 | -74.3 |
| Linochilus ochraceus | 2244-10 | 4.4581 | -74.0439 |
| Linochilus ochraceus | 159 | 4.5 | -73.73333 |
| Linochilus ochraceus | 611 | 4.52435398 | -73.746429 |
| Linochilus ochraceus | 56 | 4.52925015 | -73.732224 |
| Linochilus ochraceus | 603 | 4.52925205 | -73.732224 |
| Linochilus ochraceus | 25989 | 4.5354 | -74.1713 |
| Linochilus ochraceus | 4411 | 4.5355 | -74.0331 |
| Linochilus ochraceus | 6705 | 4.543 | -74.3108 |
| Linochilus ochraceus | 939 | 4.54810286 | -73.777023 |
| Linochilus ochraceus | 5894 | 4.5623 | -73.9869 |
| Linochilus ochraceus | 413 | 4.57 | -74.03 |
| Linochilus ochraceus | 5472 | 4.5967 | -74.1431 |
| Linochilus ochraceus | 9437 | 4.6143 | -74.0413 |
| Linochilus ochraceus | 59 | 4.6159 | -74.0431 |
| Linochilus ochraceus | 156 | 4.62 | -74.07 |
| Linochilus ochraceus | 25597 | 4.6492 | -73.7483 |
| Linochilus ochraceus | 5410 | 4.6679 | -74.0402 |
| Linochilus ochraceus | 5522 | 4.6679 | -74.0402 |
| Linochilus ochraceus | 1731 | 4.71667194 | -73.764038 |
| Linochilus ochraceus | 7994 | 4.7383 | -74.0089 |
| Linochilus ochraceus | 1173 | 4.7884 | -73.7974 |
| Linochilus ochraceus | 36 | 4.81111111 | -73.849167 |
| Linochilus ochraceus | 4047 | 4.84444444 | -73.826389 |
| Linochilus ochraceus | 5715 | 4.8786 | -73.8848 |
| Linochilus ochraceus | 9461 | 4.9 | -73.78 |
| Linochilus ochraceus | 634 | 4.990418 | -74.194966 |
| Linochilus ochraceus | 9542 | 5.03 | -74 |
| Linochilus ochraceus | 4237 | 5.0484 | -74.1585 |
| Linochilus ochraceus | 231 | 5.5248 | -73.964 |
| Linochilus ochraceus | 11 | 4.59187 | -74.03081 |
| Linochilus parvifolius | 545 | 10.7972 | -73.6454 |
| Linochilus parvifolius | 7053 | 10.85 | -73.68 |
| Linochilus parvifolius | 388 | 10.86534 | -73.7217 |
| Linochilus perijaensis | 11212 | 10.0154 | -72.9588 |
| Linochilus perijaensis | 11249 | 10.0154 | -72.9588 |
| Linochilus perijaensis | 11378 | 10.0154 | -72.9588 |
| Linochilus phylicoides | 1108 | 3.75 | -74.42 |
| Linochilus phylicoides | 1517 | 3.75 | -74.42 |
| Linochilus phylicoides | 1519 | 3.75 | -74.42 |
| Linochilus phylicoides | 8368 | 3.75 | -74.42 |
| Linochilus phylicoides | 6398 | 3.94152594 | -74.382668 |
| Linochilus phylicoides | 6399 | 3.94152594 | -74.382668 |
| Linochilus phylicoides | 2555 | 3.9445 | -74.1107 |
| Linochilus phylicoides | 2411 | 3.9544 | -74.158 |
| Linochilus phylicoides | 185 | 3.9544 | -74.158 |
| Linochilus phylicoides | 4477 | 4.2824 | -74.216 |
| Linochilus phylicoides | 25758 | 4.2842 | -74.209 |
| Linochilus phylicoides | 6156 | 4.28537 | -74.20828 |
| Linochilus phylicoides | 237 | 4.28777778 | -74.214444 |
| Linochilus phylicoides | 242 | 4.28777778 | -74.214444 |
| Linochilus phylicoides | 17567 | 4.3166649 | -73.99506 |
| Linochilus phylicoides | 6231 | 4.3845 | -74.171 |
| Linochilus phylicoides | 25920 | 4.3845 | -74.171 |
| Linochilus phylicoides | 196 | 4.387 | -74.167 |
| Linochilus phylicoides | 98 | 4.43725 | -74.20745 |
| Linochilus phylicoides | 5139 | 4.4502 | -74.05 |
| Linochilus phylicoides | 1053 | 4.461 | -74.1254 |
| Linochilus phylicoides | 4483 | 4.4643792 | -74.076476 |
| Linochilus phylicoides | 21085 | 4.4660906 | -74.072163 |
| Linochilus phylicoides | 16 | 4.473 | -74.116 |
| Linochilus phylicoides | 12026 | 4.4782 | -73.595 |
| Linochilus phylicoides | 6998 | 4.4858 | -74.1753 |
| Linochilus phylicoides | 26986 | 4.52 | -73.75 |
| Linochilus phylicoides | 96 | 4.52 | -73.75 |
| Linochilus phylicoides | 6454 | 4.52 | -73.75 |
| Linochilus phylicoides | 6 | 4.52925015 | -73.732224 |
| Linochilus phylicoides | 1 | 4.5333 | -73.9333 |
| Linochilus phylicoides | 6736 | 4.53585005 | -73.756523 |
| Linochilus phylicoides | 3224 | 4.544 | -73.9353 |
| Linochilus phylicoides | 390 | 4.5462 | -73.7513 |
| Linochilus phylicoides | 22524 | 4.563 | -74.0098 |
| Linochilus phylicoides | 16184 | 4.5633 | -74.0808 |
| Linochilus phylicoides | 923 | 4.56533718 | -74.005692 |
| Linochilus phylicoides | 929 | 4.56534004 | -74.005692 |
| Linochilus phylicoides | 7765 | 4.57 | -74.03 |
| Linochilus phylicoides | 2752 | 4.57 | -74.03 |
| Linochilus phylicoides | 329 | 4.57 | -74.03 |
| Linochilus phylicoides | 188 | 4.57 | -74.03 |
| Linochilus phylicoides | 204 | 4.57 | -74.03 |
| Linochilus phylicoides | 222 | 4.57 | -74.03 |
| Linochilus phylicoides | 228 | 4.57 | -74.03 |
| Linochilus phylicoides | 231 | 4.57 | -74.03 |
| Linochilus phylicoides | 314 | 4.57 | -74.03 |
| Linochilus phylicoides | 1337 | 4.57 | -74.03 |
| Linochilus phylicoides | 1339 | 4.57 | -74.03 |
| Linochilus phylicoides | 2514 | 4.593395 | -74.05025 |
| Linochilus phylicoides | 361 | 4.5934933 | -74.060276 |
| Linochilus phylicoides | 5596 | 4.5971 | -74.1566 |
| Linochilus phylicoides | 5634 | 4.5971 | -74.1566 |
| Linochilus phylicoides | 2301-2 | 4.602 | -74.029 |
| Linochilus phylicoides | 01302A | 4.6039737 | -74.058237 |
| Linochilus phylicoides | 29 | 4.6056 | -73.9969 |
| Linochilus phylicoides | 4374 | 4.6066259 | -74.04871 |
| Linochilus phylicoides | 11922 | 4.6159 | -74.0416 |
| Linochilus phylicoides | 608 | 4.6189 | -73.7477 |
| Linochilus phylicoides | 5172 | 4.62 | -74.07 |
| Linochilus phylicoides | 12 | 4.62 | -74.07 |
| Linochilus phylicoides | 2981 | 4.6336 | -74.0548 |
| Linochilus phylicoides | 4066A | 4.657 | -74.037 |
| Linochilus phylicoides | 25613 | 4.662 | -73.887 |
| Linochilus phylicoides | 14 | 4.662 | -73.887 |
| Linochilus phylicoides | 19417 | 4.67863321 | -73.911903 |
| Linochilus phylicoides | 20501 | 4.690278 | -73.819655 |
| Linochilus phylicoides | 209 | 4.70856905 | -73.911491 |
| Linochilus phylicoides | 210 | 4.70856905 | -73.911491 |
| Linochilus phylicoides | 26917 | 4.70856905 | -73.911491 |
| Linochilus phylicoides | 3671 | 4.70856905 | -73.911491 |
| Linochilus phylicoides | 2462 | 4.71061516 | -73.907806 |
| Linochilus phylicoides | 2455 | 4.7198 | -73.978 |
| Linochilus phylicoides | 60 | 4.75833321 | -73.833054 |
| Linochilus phylicoides | 34097 | 4.77565384 | -73.826576 |
| Linochilus phylicoides | 34151 | 4.77565384 | -73.826576 |
| Linochilus phylicoides | 8114 | 4.78373623 | -73.797546 |
| Linochilus phylicoides | 439 | 4.79288387 | -73.862473 |
| Linochilus phylicoides | 784 | 4.79677916 | -73.814171 |
| Linochilus phylicoides | 29 | 4.81111111 | -73.849167 |
| Linochilus phylicoides | 4067 | 4.81111111 | -73.849167 |
| Linochilus phylicoides | 12 | 4.81111111 | -73.849167 |
| Linochilus phylicoides | 9423 | 4.8143 | -73.5467 |
| Linochilus phylicoides | 3 | 4.85117817 | -73.801628 |
| Linochilus phylicoides | 1601 | 4.8666 | -73.8833 |
| Linochilus phylicoides | 4226 | 4.87 | -74.22 |
| Linochilus phylicoides | 12063 | 4.87 | -74.22 |
| Linochilus phylicoides | 3618 | 4.99509478 | -74.180603 |
| Linochilus phylicoides | 7 | 4.99849987 | -74.189468 |
| Linochilus phylicoides | 34 | 5.00732803 | -74.19165 |
| Linochilus phylicoides | 609 | 5.04200888 | -74.153008 |
| Linochilus phylicoides | 444 | 5.04619503 | -74.02227 |
| Linochilus phylicoides | 2277 | 5.0901296 | -74.08287 |
| Linochilus phylicoides | 367 | 5.21666718 | -74.033333 |
| Linochilus phylicoides | 286 | 5.219641 | -74.00995 |
| Linochilus phylicoides | 6239 | 5.2253 | -73.9884 |
| Linochilus phylicoides | 8677 | 5.3 | -74.07 |
| Linochilus phylicoides | 20082 | 5.67 | -73.45 |
| Linochilus phylicoides | 41 | 5.68496 | -73.451799 |
| Linochilus phylicoides | 30 | 5.68496 | -73.451799 |
| Linochilus phylicoides | 9862 | 5.69000006 | -73.440277 |
| Linochilus phylicoides | 11611 | 5.90088987 | -73.067741 |
| Linochilus phylicoides | 1261 | 5.94805556 | -73.101944 |
| Linochilus phylicoides | 7156 | 5.98 | -73.08 |
| Linochilus phylicoides | 3465 | 5.98 | -73.08 |
| Linochilus phylicoides | s.n. | 4.59187 | -74.03081 |
| Linochilus pittieri | 4106 | 1.92 | -76.6 |
| Linochilus pittieri | 2690 | 2.1814 | -76.4307 |
| Linochilus pittieri | 26327 | 2.4 | -76.38 |
| Linochilus pittieri | 26365 | 2.5261 | -76.3605 |
| Linochilus pittieri | 2512 | 2.963 | -76.11 |
| Linochilus pittieri | 1156 | 3.7493 | -75.9464 |
| Linochilus rangelii | 994 | 10.8983 | -73.7493 |
| Linochilus revolutus | 8437 | 3.75 | -74.42 |
| Linochilus revolutus | 917 | 3.75 | -74.42 |
| Linochilus revolutus | 25875 | 3.75 | -74.42 |
| Linochilus revolutus | 54 | 3.75 | -74.42 |
| Linochilus revolutus | 5767 | 3.75 | -74.42 |
| Linochilus revolutus | 2499 | 3.9564 | -74.1608 |
| Linochilus revolutus | 10376 | 4.12 | -74.25 |
| Linochilus revolutus | 27049 | 4.12 | -74.25 |
| Linochilus revolutus | 6522 | 4.1738 | -75.8019 |
| Linochilus revolutus | 208 | 4.27947807 | -74.199463 |
| Linochilus revolutus | 6142 | 4.280897 | -74.220114 |
| Linochilus revolutus | 23 | 4.28168678 | -74.230339 |
| Linochilus revolutus | 4481 | 4.2847 | -74.2086 |
| Linochilus revolutus | 25732 | 4.2847 | -74.2078 |
| Linochilus revolutus | 25967 | 4.2847 | -74.2078 |
| Linochilus revolutus | 141 | 4.28927517 | -74.207314 |
| Linochilus revolutus | 409 | 4.3837 | -74.1678 |
| Linochilus revolutus | 915 | 4.521324 | -74.307575 |
| Linochilus revolutus | 3220 | 4.55 | -73.97 |
| Linochilus revolutus | 708 | 4.5608961 | -73.971848 |
| Linochilus revolutus | 2 | 4.5639 | -74.0045 |
| Linochilus revolutus | 931 | 4.56533718 | -74.005692 |
| Linochilus revolutus | 2963 | 4.57 | -74.03 |
| Linochilus revolutus | 146 | 4.66805 | -74.10004 |
| Linochilus revolutus | 4533 | 4.7948 | -75.3643 |
| Linochilus revolutus | 8061 | 4.91222222 | -75.376667 |
| Linochilus revolutus | 78 | 4.9486 | -75.4102 |
| Linochilus revolutus | 9365 | 5.5051 | -72.7435 |
| Linochilus revolutus | 255 | 5.5345 | -72.9283 |
| Linochilus revolutus | 2783 | 5.5345 | -72.9283 |
| Linochilus revolutus | 1229 | 5.94805556 | -73.101944 |
| Linochilus revolutus | 6865 | 5.98 | -73.08 |
| Linochilus revolutus | 6856 | 5.98 | -73.08 |
| Linochilus revolutus | 7289 | 5.98 | -73.08 |
| Linochilus revolutus | 10416 | 5.98 | -73.08 |
| Linochilus revolutus | 3441 | 5.98 | -73.08 |
| Linochilus revolutus | 7490 | 6.42 | -72.3 |
| Linochilus revolutus | 1369 | 6.42 | -72.3 |
| Linochilus revolutus | 274 | 6.486613 | -72.332023 |
| Linochilus revolutus | 1540 | 6.549573 | -72.328398 |
| Linochilus revolutus | 13513 | 6.95 | -72.68 |
| Linochilus revolutus | 27879 | 6.95 | -72.68 |
| Linochilus revolutus | 27889 | 6.95 | -72.68 |
| Linochilus revolutus | 9906A | 6.95 | -72.68 |
| Linochilus revolutus | 9940 | 6.95 | -72.68 |
| Linochilus revolutus | 7661 | 6.95 | -72.68 |
| Linochilus revolutus | 10294 | 7.27 | -72.88 |
| Linochilus revolutus | 28750 | 7.38 | -72.87 |
| Linochilus revolutus | 1069 | 7.41013 | -72.85549 |
| Linochilus rhododendroides | 12785 | 1.087 | -77.6631 |
| Linochilus rhododendroides | 1878A | 1.09067 | -77.71952 |
| Linochilus rhododendroides | 429 | 1.09067 | -77.71952 |
| Linochilus rhododendroides | 1020 | 1.0909 | -77.1571 |
| Linochilus rhododendroides | 28650 | 1.13 | -77.12 |
| Linochilus rhododendroides | 11755 | 1.1449 | -77.1012 |
| Linochilus rhododendroides | 1018 | 1.1449 | -77.1012 |
| Linochilus rhododendroides | 3984 | 4.1924 | -76.4594 |
| Linochilus rhomboidalis | 1450 | 6.378382 | -72.33189 |
| Linochilus rhomboidalis | 5761 | 6.385995 | -72.344335 |
| Linochilus rhomboidalis | 137 | 6.399974 | -72.311559 |
| Linochilus rhomboidalis | 8600 | 6.401111 | -72.309351 |
| Linochilus rhomboidalis | 8728 | 6.401275 | -72.337769 |
| Linochilus rhomboidalis | 8754 | 6.401275 | -72.337769 |
| Linochilus rhomboidalis | 9986 | 6.405338 | -72.373902 |
| Linochilus rhomboidalis | 27820 | 6.405436 | -72.372883 |
| Linochilus rhomboidalis | 7373 | 6.5399 | -72.3168 |
| Linochilus rhomboidalis | 1539 | 6.546737 | -72.325109 |
| Linochilus rhomboidalis | 340 | 6.98556 | -72.68306 |
| Linochilus rhomboidalis | 25904 | 4.2877 | -74.2075 |
| Linochilus ritterbushii | s.n. | 3.01667 | -76 |
| Linochilus romeroi | 6775 | 10.8151 | -73.7169 |
| Linochilus romeroi | 7115 | 10.8204 | -73.7656 |
| Linochilus rosmarinifolius | 100 | 2.49611111 | -76.630278 |
| Linochilus rosmarinifolius | 142 | 3.9555 | -74.1611 |
| Linochilus rosmarinifolius | 3479 | 4.2827 | -74.3074 |
| Linochilus rosmarinifolius | 4063 | 4.2827 | -74.3074 |
| Linochilus rosmarinifolius | 7049 | 4.2827 | -74.3074 |
| Linochilus rosmarinifolius | 34193 | 4.4646787 | -74.077506 |
| Linochilus rosmarinifolius | 651 | 4.47 | -74.1266 |
| Linochilus rosmarinifolius | 25929 | 4.5099 | -74.0478 |
| Linochilus rosmarinifolius | 9 | 4.5499 | -74.1333 |
| Linochilus rosmarinifolius | 16145 | 4.5697 | -74.0666 |
| Linochilus rosmarinifolius | 1114 | 4.5947 | -74.0504 |
| Linochilus rosmarinifolius | 1277 | 4.6039737 | -74.058237 |
| Linochilus rosmarinifolius | 38000 | 4.616 | -74.0415 |
| Linochilus rosmarinifolius | 9206 | 4.6404 | -74.0488 |
| Linochilus rosmarinifolius | 85 | 4.6645 | -74.2764 |
| Linochilus rosmarinifolius | 8069 | 4.6701 | -74.0166 |
| Linochilus rosmarinifolius | 46-406 | 4.674 | -74.036 |
| Linochilus rosmarinifolius | 5001 | 4.6772 | -74.0323 |
| Linochilus rosmarinifolius | 19415 | 4.67863321 | -73.911903 |
| Linochilus rosmarinifolius | 1763 | 4.7101 | -74.3512 |
| Linochilus rosmarinifolius | 27 | 4.7181 | -74.0077 |
| Linochilus rosmarinifolius | 5548 | 4.7181 | -74.0077 |
| Linochilus rosmarinifolius | 1297 | 4.7211 | -73.8938 |
| Linochilus rosmarinifolius | 2436 | 4.7264 | -73.9821 |
| Linochilus rosmarinifolius | 356-A | 4.7366 | -74.0167 |
| Linochilus rosmarinifolius | 54 | 4.7366 | -74.0167 |
| Linochilus rosmarinifolius | 134 | 4.7366 | -74.0167 |
| Linochilus rosmarinifolius | 22 | 4.7527 | -74.3829 |
| Linochilus rosmarinifolius | 24 | 4.78444444 | -74.023889 |
| Linochilus rosmarinifolius | 11530 | 4.80414009 | -73.805817 |
| Linochilus rosmarinifolius | 6254 | 4.815382 | -73.786163 |
| Linochilus rosmarinifolius | 3124 | 4.8392 | -73.8081 |
| Linochilus rosmarinifolius | 7732 | 4.8392 | -73.8081 |
| Linochilus rosmarinifolius | 5708 | 4.9 | -73.78 |
| Linochilus rosmarinifolius | 13381 | 4.9 | -73.78 |
| Linochilus rosmarinifolius | 221-A | 4.9341 | -73.8527 |
| Linochilus rosmarinifolius | 222 | 4.9341 | -73.8527 |
| Linochilus rosmarinifolius | 973 | 4.99916667 | -74.150278 |
| Linochilus rosmarinifolius | 13629 | 5.0666 | -73.9833 |
| Linochilus rosmarinifolius | 224 | 5.103 | -73.7069 |
| Linochilus rosmarinifolius | 1800 | 5.1391 | -76.1796 |
| Linochilus rosmarinifolius | 47 | 5.14375 | -73.9756 |
| Linochilus rosmarinifolius | 8087 | 5.446238 | -73.433715 |
| Linochilus rosmarinifolius | 125 | 5.52546 | -73.129677 |
| Linochilus rosmarinifolius | 150 | 5.5342 | -72.9283 |
| Linochilus rosmarinifolius | 28691 | 5.5342 | -72.9283 |
| Linochilus rosmarinifolius | 150 | 5.5342 | -72.9283 |
| Linochilus rosmarinifolius | 3529 | 5.7421 | -73.843 |
| Linochilus rosmarinifolius | 1459 | 5.756158 | -73.415369 |
| Linochilus rosmarinifolius | 5756 | 5.98 | -73.08 |
| Linochilus rosmarinifolius | 2069 | 6.0357 | -72.8999 |
| Linochilus rosmarinifolius | 2580 | 6.3216 | -75.4697 |
| Linochilus rosmarinifolius | 580 | 6.47 | -76.1 |
| Linochilus rosmarinifolius | 17 | 6.8558 | -72.7176 |
| Linochilus rosmarinifolius | 13529 | 6.95 | -72.68 |
| Linochilus rosmarinifolius | 10266 | 7.3406 | -72.7195 |
| Linochilus rosmarinifolius | 18080 | 7.3631 | -72.8845 |
| Linochilus rosmarinifolius | 10241 | 7.3968 | -72.6474 |
| Linochilus rosmarinifolius | 12641 | 7.42 | -72.43 |
| Linochilus rosmarinifolius | 11014 | 10.2512 | -73.01 |
| Linochilus rosmarinifolius | 11262 | 10.3589 | -72.9091 |
| Linochilus rosmarinifolius | 25048 | 10.3733 | -72.8966 |
| Linochilus rosmarinifolius | 10863 | 10.3733 | -72.8966 |
| Linochilus rosmarinifolius | 923 | 10.65 | -74.02389 |
| Linochilus rosmarinifolius | 594 | 10.7107 | -73.6707 |
| Linochilus rosmarinifolius | 1379 | 10.7801 | -73.6837 |
| Linochilus rosmarinifolius | 6604 | 10.8868 | -73.7979 |
| Linochilus rosmarinifolius | 6778 | 10.8868 | -73.7979 |
| Linochilus rupestris | 2928 | 0.812 | -77.847 |
| Linochilus rupestris | 2937 | 0.8125 | -77.8466 |
| Linochilus rupestris | 120 | 0.8671 | -77.8597 |
| Linochilus rupestris | 26296 | 2.3701 | -76.3572 |
| Linochilus rupestris | 56A | 3.3421 | -76.6922 |
| Linochilus rupestris | 140 | 3.3515 | -76.6984 |
| Linochilus rupestris | 1293 | 3.75 | -74.42 |
| Linochilus rupestris | 1331 | 3.75 | -74.42 |
| Linochilus rupestris | 2575 | 3.9343 | -74.1103 |
| Linochilus rupestris | 1895 | 4.6796 | -75.35067 |
| Linochilus rupestris | 5795 | 4.765547 | -75.439799 |
| Linochilus rupestris | 5742 | 4.782 | -75.4052 |
| Linochilus rupestris | 5764 | 4.782 | -75.4052 |
| Linochilus rupestris | 1768 | 4.8461 | -75.3703 |
| Linochilus rupestris | 9287 | 4.9 | -75.3 |
| Linochilus rupestris | s.n. | 4.9 | -75.3 |
| Linochilus rupestris | 27036 | 4.9 | -75.3 |
| Linochilus rupestris | 5956 | 4.9 | -75.3 |
| Linochilus rupestris | 3638 | 4.9516 | -75.3596 |
| Linochilus rupestris | 10635 | 5.041 | -75.3339 |
| Linochilus rupestris | 2426 | 5.1591 | -76.0994 |
| Linochilus rupestris | 2427 | 5.164 | -76.095 |
| Linochilus rupestris | 401 | 6.45555544 | -76.120552 |
| Linochilus rupestris | 268 | 6.47 | -76.1 |
| Linochilus rupestris | 2361 | 6.47 | -76.1 |
| Linochilus rupestris | 2164 | 6.5 | -76.116669 |
| Linochilus rupestris | 2882 | 4.65992 | -75.33995 |
| Linochilus santamartae | 1808 | 10.8672 | -73.6204 |
| Linochilus santamartae | 928 | 10.8726 | -73.6441 |
| Linochilus saxatilis | 24620 | 10.78769 | -73.59226 |
| Linochilus saxatilis | 6752 | 10.8792 | -73.896 |
| Linochilus saxatilis | 986 | 10.8921 | -73.7801 |
| Linochilus schultzii | 146 | 2.36712 | -76.3677 |
| Linochilus schultzii | 1891 | 4.6796 | -75.35067 |
| Linochilus schultzii | 6250 | 4.6796 | -75.35067 |
| Linochilus schultzii | 5688 | 4.7871 | -75.39607 |
| Linochilus schultzii | 5791 | 4.7906 | -75.4019 |
| Linochilus schultzii | 1184 | 4.7993 | -75.4059 |
| Linochilus schultzii | 8083 | 4.8236 | -75.3618 |
| Linochilus schultzii | 9267 | 4.9 | -75.3 |
| Linochilus schultzii | 6435 | 4.9355 | -75.3491 |
| Linochilus schultzii | 1943 | 5.1163 | -76.0854 |
| Linochilus schultzii | 1611 | 5.1497 | -76.0395 |
| Linochilus schultzii | 2425 | 5.1621 | -76.0921 |
| Linochilus schultzii | 901 | 4.65992 | -75.33995 |
| Linochilus schultzii | 114 | 0.8388 | -77.9243 |
| Linochilus schultzii | 3466 | 1.35587 | -76.90579 |
| Linochilus schultzii | 4010 | 1.9089 | -76.6527 |
| Linochilus schultzii | 5958 | 1.92 | -76.6 |
| Linochilus schultzii | 83 | 1.93259 | -76.62626 |
| Linochilus schultzii | 4893 | 2.38815 | -76.2755 |
| Linochilus schultzii | 27438 | 2.83 | -76.2 |
| Linochilus schultzii | 28636 | 2.83 | -76.2 |
| Linochilus schultzii | 2466 | 3.6144 | -74.22809 |
| Linochilus schultzii | 164 | 4.74136 | -73.86186 |
| Linochilus schultzii | 2217 | 4.9 | -73.78 |
| Linochilus schultzii | 528 | 6.46139 | -76.1275 |
| Linochilus schultzii | 26259 | 2.34113 | -76.23647 |
| Linochilus schultzii | 12809 | 2.36576 | -76.37153 |
| Linochilus schultzii | 26362 | 2.6 | -76.25 |
| Linochilus schultzii | 19041 | 2.85728 | -76.10485 |
| Linochilus schultzii | 316 | 7.41907 | -72.34351 |
| Linochilus tachirensis | 337 | 7.34453 | -72.61139 |
| Linochilus tachirensis | 57360 | 7.41847 | -72.36488 |
| Linochilus tamanus | 12710 | 7.39379 | -72.42806 |
| Linochilus tenuifolius | 5097 | 5.15972 | -73.45066 |
| Linochilus tenuifolius | 7811 | 5.16098 | -73.25905 |
| Linochilus tenuifolius | 28710 | 5.44174 | -73.9101 |
| Linochilus tenuifolius | 2729 | 5.486606 | -72.725196 |
| Linochilus tenuifolius | 254 | 5.53 | -73.13 |
| Linochilus tenuifolius | 3 | 5.54437 | -72.9258 |
| Linochilus tenuifolius | 457 | 5.61389017 | -72.913651 |
| Linochilus tenuifolius | 2858 | 5.61697 | -73.35682 |
| Linochilus tenuifolius | 6213 | 5.70446 | -73.3753 |
| Linochilus tenuifolius | 3008 | 5.72942 | -73.41051 |
| Linochilus tenuifolius | 13205 | 5.74487 | -73.49643 |
| Linochilus tenuifolius | 28666 | 5.76791 | -73.41947 |
| Linochilus tenuifolius | 20261A | 5.775515 | -73.426014 |
| Linochilus tenuifolius | 20271 | 5.81481 | -73.4686 |
| Linochilus tenuifolius | 7186 | 5.8281 | -72.79461 |
| Linochilus tenuifolius | 242 | 5.9168 | -73.1664 |
| Linochilus tenuifolius | 2816 | 5.99812 | -72.58934 |
| Linochilus tenuifolius | 863 | 6.21944 | -72.7846 |
| Linochilus tenuifolius | 1793 | 6.28435 | -72.53487 |
| Linochilus tenuifolius | 27809 | 6.289406 | -72.574213 |
| Linochilus tenuifolius | 2118 | 6.46743 | -72.403405 |
| Linochilus tergocanus | 6762 | 10.89332 | -73.88254 |
| Linochilus venezuelensis | 2559 | 7.94785611 | -72.080187 |
| Linochilus venezuelensis | 27994 | 7.9713828 | -72.081276 |
| Linochilus venezuelensis | 8133 | 8.16208861 | -71.886184 |
| Linochilus venezuelensis | 89V33 | 8.56172289 | -71.088186 |
| Linochilus venezuelensis | 9950 | 8.5914537 | -71.018935 |
| Linochilus venezuelensis | 2585 | 8.61191657 | -71.031 |
| Linochilus venezuelensis | 28126 | 8.78785332 | -70.807572 |
| Linochilus venezuelensis | 10815 | 8.7979491 | -70.825283 |
| Linochilus venezuelensis | 28040 | 8.99915812 | -70.867365 |
| Linochilus venezuelensis | 28182 | 9.06707776 | -70.634872 |
| Linochilus venezuelensis | 3447 | 9.07552859 | -70.618523 |
| Linochilus venezuelensis | 104819 | 9.23781401 | -70.185546 |
| Linochilus violaceus | s.n. | 4.63323 | -75.32717 |
| Linochilus violaceus | s.n. | 4.63323 | -75.32717 |
| Linochilus violaceus | 9229 | 4.97076 | -75.38358 |
| Linochilus violaceus | 6302 | 5.06217 | -75.32758 |

**APPENDIX S2**. Leaf area measurements.

| **Species** | **id** | **area_mm** | **area_log** |
| --- | --- | --- | --- |
| *Linochilus_alveolatus* | P_alve_0000_Layer_16 | 122.206338 | 6.93317529 |
| *Linochilus_alveolatus* | P_alve_0001_Layer_15 | 130.403487 | 7.02683863 |
| *Linochilus_alveolatus* | P_alve_0002_Layer_14 | 92.8828023 | 6.53733959 |
| *Linochilus_alveolatus* | P_alve_0005_Layer_13 | 125.437527 | 6.97082521 |
| *Linochilus_alveolatus* | P_alve_0006_Layer_12 | 52.2241188 | 5.70664434 |
| *Linochilus_alveolatus* | P_alve_0007_Layer_11 | 62.1882964 | 5.95857119 |
| *Linochilus_alveolatus* | P_alve_0008_Layer_10 | 70.1130435 | 6.13161096 |
| *Linochilus_alveolatus* | P_alve_0009_Layer_9 | 105.278884 | 6.7180723 |
| *Linochilus_alveolatus* | P_alve_0010_Layer_7 | 97.6543172 | 6.60961192 |
| *Linochilus_alveolatus* | P_alve_0012_Layer_25 | 147.76284 | 7.20713969 |
| *Linochilus_alveolatus* | P_alve_0013_Layer_24 | 126.167815 | 6.97920012 |
| *Linochilus_alveolatus* | P_alve_0015_Layer_23 | 132.019977 | 7.04461244 |
| *Linochilus_alveolatus* | P_alve_0016_Layer_22 | 36.0242656 | 5.17089711 |
| *Linochilus_alveolatus* | P_alve_0017_Layer_21 | 68.7680587 | 6.10366671 |
| *Linochilus_alveolatus* | P_alve_0018_Layer_20 | 37.9463125 | 5.24588779 |
| *Linochilus_alveolatus* | P_alve_0019_Layer_19 | 138.489524 | 7.11363304 |
| *Linochilus_alveolatus* | P_alve_0021_Layer_18 | 95.4132733 | 6.57611807 |
| *Linochilus_alveolatus* | P_alve_0022_Layer_17 | 137.721602 | 7.10561105 |
| *Linochilus_alveolatus* | P_alve_0024_Layer_32 | 75.7671766 | 6.24350108 |
| *Linochilus_alveolatus* | P_alve_0025_Layer_31 | 50.2303872 | 5.65048849 |
| *Linochilus_alveolatus* | P_alve_0026_Layer_30 | 90.6489268 | 6.50221804 |
| *Linochilus_alveolatus* | P_alve_0027_Layer_29 | 163.205524 | 7.35054607 |
| *Linochilus_alveolatus* | P_alve_0029_Layer_28 | 142.177704 | 7.15155143 |
| *Linochilus_alveolatus* | P_alve_0030_Layer_27 | 156.011065 | 7.28550454 |
| *Linochilus_alveolatus* | P_alve_0031_Layer_26 | 93.1032328 | 6.54075936 |
| *Linochilus_alveolatus* | P_alve_0033_Layer_5 | 41.060118 | 5.35966587 |
| *Linochilus_alveolatus* | P_alve_0034_Layer_4 | 35.863871 | 5.16445931 |
| *Linochilus_alveolatus* | P_alve_0035_Layer_3 | 40.6022317 | 5.34348712 |
| *Linochilus_alveolatus* | P_alve_0036_Layer_2 | 38.8253465 | 5.27892689 |
| *Linochilus_alveolatus* | P_alve_0038_Layer_1 | 49.1309226 | 5.61855943 |
| *Linochilus_antioquensis* | P_anti_0000_Layer_11 | 559.037856 | 9.12680217 |
| *Linochilus_antioquensis* | P_anti_0001_Layer_10 | 560.778003 | 9.13128595 |
| *Linochilus_antioquensis* | P_anti_0002_Layer_9 | 872.03669 | 9.76824503 |
| *Linochilus_antioquensis* | P_anti_0003_Layer_8 | 575.029824 | 9.16749297 |
| *Linochilus_antioquensis* | P_anti_0004_Layer_7 | 1066.02185 | 10.0580213 |
| *Linochilus_antioquensis* | P_anti_0007_Layer_5 | 1583.42342 | 10.6288314 |
| *Linochilus_antioquensis* | P_anti_0008_Layer_4 | 903.569011 | 9.81949098 |
| *Linochilus_antioquensis* | P_anti_0009_Layer_3 | 704.280979 | 9.46000731 |
| *Linochilus_antioquensis* | P_anti_0010_Layer_2 | 595.024488 | 9.21680523 |
| *Linochilus_antioquensis* | P_anti_0011_Layer_1 | 1265.28837 | 10.3052505 |
| *Linochilus_apiculatus* | P_apic_0000_Layer_6 | 10.379053 | 3.37560291 |
| *Linochilus_apiculatus* | P_apic_0001_Layer_5 | 7.78316969 | 2.96035781 |
| *Linochilus_apiculatus* | P_apic_0002_Layer_4 | 9.06901455 | 3.18094579 |
| *Linochilus_apiculatus* | P_apic_0003_Layer_3 | 7.30288199 | 2.86846592 |
| *Linochilus_apiculatus* | P_apic_0004_Layer_2 | 6.29302334 | 2.65375329 |
| *Linochilus_apiculatus* | P_apic_0005_Layer_25 | 5.76345239 | 2.52693326 |
| *Linochilus_apiculatus* | P_apic_0007_Layer_23 | 9.50091506 | 3.24806647 |
| *Linochilus_apiculatus* | P_apic_0008_Layer_22 | 8.28317069 | 3.05018312 |
| *Linochilus_apiculatus* | P_apic_0010_Layer_21 | 7.71775737 | 2.94818169 |
| *Linochilus_apiculatus* | P_apic_0011_Layer_20 | 5.28674893 | 2.40238081 |
| *Linochilus_apiculatus* | P_apic_0012_Layer_19 | 3.16846512 | 1.66378413 |
| *Linochilus_apiculatus* | P_apic_0013_Layer_18 | 7.33962041 | 2.87570545 |
| *Linochilus_apiculatus* | P_apic_0014_Layer_17 | 7.42474603 | 2.89234168 |
| *Linochilus_apiculatus* | P_apic_0015_Layer_16 | 8.78406775 | 3.13488918 |
| *Linochilus_apiculatus* | P_apic_0016_Layer_15 | 6.84230759 | 2.77448296 |
| *Linochilus_apiculatus* | P_apic_0017_Layer_14 | 7.25897509 | 2.85976587 |
| *Linochilus_apiculatus* | P_apic_0018_Layer_13 | 11.2903452 | 3.49701769 |
| *Linochilus_apiculatus* | P_apic_0020_Layer_12 | 11.09142 | 3.47137218 |
| *Linochilus_apiculatus* | P_apic_0021_Layer_11 | 13.9426802 | 3.80143601 |
| *Linochilus_apiculatus* | P_apic_0022_Layer_10 | 9.50270718 | 3.24833857 |
| *Linochilus_apiculatus* | P_apic_0023_Layer_9 | 18.4588183 | 4.20623829 |
| *Linochilus_apiculatus* | P_apic_0024_Layer_8 | 23.4014805 | 4.5485279 |
| *Linochilus_apiculatus* | P_apic_0026_Layer_7 | 20.9077479 | 4.38596576 |
| *Linochilus_apiculatus* | P_apic_0029_Layer_1 | 7.46417263 | 2.89998235 |
| *Linochilus_camargoanus* | P_cama_0000_Layer_8 | 1175.81597 | 10.1994466 |
| *Linochilus_camargoanus* | P_cama_0001_Layer_7 | 512.428444 | 9.00120675 |
| *Linochilus_camargoanus* | P_cama_0002_Layer_6 | 860.121792 | 9.74839715 |
| *Linochilus_camargoanus* | P_cama_0003_Layer_5 | 657.414397 | 9.36065924 |
| *Linochilus_camargoanus* | P_cama_0004_Layer_4 | 1321.71143 | 10.3681915 |
| *Linochilus_camargoanus* | P_cama_0007_Layer_2 | 1277.71044 | 10.3193452 |
| *Linochilus_cayambensis* | P_caya_0000_Layer_25 | 38.0538396 | 5.24997012 |
| *Linochilus_cayambensis* | P_caya_0001_Layer_18 | 25.2133121 | 4.65611374 |
| *Linochilus_cayambensis* | P_caya_0002_Layer_17 | 39.5171041 | 5.30440532 |
| *Linochilus_cayambensis* | P_caya_0004_Layer_16 | 43.8620949 | 5.45490281 |
| *Linochilus_cayambensis* | P_caya_0005_Layer_15 | 69.196375 | 6.11262456 |
| *Linochilus_cayambensis* | P_caya_0006_Layer_14 | 58.2922313 | 5.86523172 |
| *Linochilus_cayambensis* | P_caya_0007_Layer_13 | 39.6730184 | 5.31008626 |
| *Linochilus_cayambensis* | P_caya_0008_Layer_12 | 62.9580112 | 5.97631806 |
| *Linochilus_cayambensis* | P_caya_0010_Layer_10 | 72.9634076 | 6.1891012 |
| *Linochilus_cayambensis* | P_caya_0011_Layer_11 | 50.0673044 | 5.64579688 |
| *Linochilus_cayambensis* | P_caya_0012_Layer_21 | 35.4463075 | 5.14756344 |
| *Linochilus_cayambensis* | P_caya_0013_Layer_20 | 37.9149504 | 5.24469493 |
| *Linochilus_cayambensis* | P_caya_0014_Layer_19 | 24.6676121 | 4.62454616 |
| *Linochilus_cayambensis* | P_caya_0015_Layer_24 | 25.7303382 | 4.68539851 |
| *Linochilus_cayambensis* | P_caya_0016_Layer_31 | 43.4427392 | 5.44104317 |
| *Linochilus_cayambensis* | P_caya_0017_Layer_30 | 34.0394946 | 5.08913771 |
| *Linochilus_cayambensis* | P_caya_0018_Layer_29 | 35.0421848 | 5.13102082 |
| *Linochilus_cayambensis* | P_caya_0019_Layer_28 | 43.3925599 | 5.43937579 |
| *Linochilus_cayambensis* | P_caya_0020_Layer_27 | 36.2312553 | 5.17916288 |
| *Linochilus_cayambensis* | P_caya_0022_Layer_26 | 27.6864353 | 4.79110741 |
| *Linochilus_cayambensis* | P_caya_0023_Layer_23 | 25.7787254 | 4.68810903 |
| *Linochilus_cayambensis* | P_caya_0024_Layer_22 | 29.5430698 | 4.88474784 |
| *Linochilus_cayambensis* | P_caya_0026_Layer_9 | 96.6650679 | 6.59492273 |
| *Linochilus_cayambensis* | P_caya_0027_Layer_8 | 107.742151 | 6.75143896 |
| *Linochilus_cayambensis* | P_caya_0029_Layer_7 | 108.340719 | 6.75943175 |
| *Linochilus_cayambensis* | P_caya_0030_Layer_6 | 85.0512454 | 6.41026046 |
| *Linochilus_cayambensis* | P_caya_0031_Layer_5 | 68.6712843 | 6.10163504 |
| *Linochilus_cayambensis* | P_caya_0032_Layer_4 | 43.6103023 | 5.44659708 |
| *Linochilus_cayambensis* | P_caya_0033_Layer_3 | 85.6148666 | 6.41978943 |
| *Linochilus_cayambensis* | P_caya_0036_Layer_2 | 41.7043845 | 5.38212716 |
| *Linochilus_cinerascens* | P_cine_0000_Layer_32 | 26.2751422 | 4.71562666 |
| *Linochilus_cinerascens* | P_cine_0001_Layer_31 | 34.1129715 | 5.09224852 |
| *Linochilus_cinerascens* | P_cine_0002_Layer_30 | 22.2061376 | 4.47288658 |
| *Linochilus_cinerascens* | P_cine_0003_Layer_29 | 36.5197863 | 5.19060642 |
| *Linochilus_cinerascens* | P_cine_0004_Layer_28 | 30.5072295 | 4.93107926 |
| *Linochilus_cinerascens* | P_cine_0006_Layer_27 | 38.9212248 | 5.28248521 |
| *Linochilus_cinerascens* | P_cine_0007_Layer_26 | 37.1228341 | 5.21423495 |
| *Linochilus_cinerascens* | P_cine_0008_Layer_25 | 33.4113572 | 5.06226668 |
| *Linochilus_cinerascens* | P_cine_0009_Layer_24 | 33.1730054 | 5.05193782 |
| *Linochilus_cinerascens* | P_cine_0010_Layer_23 | 28.6102723 | 4.83846132 |
| *Linochilus_cinerascens* | P_cine_0013_Layer_21 | 39.5699716 | 5.30633413 |
| *Linochilus_cinerascens* | P_cine_0014_Layer_20 | 33.0726468 | 5.0475666 |
| *Linochilus_cinerascens* | P_cine_0015_Layer_19 | 40.9472145 | 5.35569341 |
| *Linochilus_cinerascens* | P_cine_0016_Layer_18 | 44.3764328 | 5.4717218 |
| *Linochilus_cinerascens* | P_cine_0018_Layer_17 | 38.5502563 | 5.26866855 |
| *Linochilus_cinerascens* | P_cine_0019_Layer_16 | 33.5045473 | 5.06628501 |
| *Linochilus_cinerascens* | P_cine_0020_Layer_15 | 31.1568724 | 4.96147851 |
| *Linochilus_cinerascens* | P_cine_0021_Layer_14 | 40.4920165 | 5.33956558 |
| *Linochilus_cinerascens* | P_cine_0022_Layer_13 | 26.2339234 | 4.71336168 |
| *Linochilus_cinerascens* | P_cine_0024_Layer_12 | 25.940016 | 4.69710747 |
| *Linochilus_cinerascens* | P_cine_0025_Layer_11 | 20.8378553 | 4.38113489 |
| *Linochilus_cinerascens* | P_cine_0026_Layer_10 | 29.1685171 | 4.86634014 |
| *Linochilus_cinerascens* | P_cine_0027_Layer_9 | 22.0170691 | 4.46055053 |
| *Linochilus_cinerascens* | P_cine_0028_Layer_8 | 35.7563439 | 5.16012732 |
| *Linochilus_cinerascens* | P_cine_0029_Layer_7 | 41.3370002 | 5.36936179 |
| *Linochilus_cinerascens* | P_cine_0031_Layer_6 | 14.0385585 | 3.8113229 |
| *Linochilus_cinerascens* | P_cine_0032_Layer_5 | 16.7034384 | 4.06207321 |
| *Linochilus_cinerascens* | P_cine_0033_Layer_4 | 30.5341113 | 4.93234995 |
| *Linochilus_cinerascens* | P_cine_0034_Layer_3 | 29.0959363 | 4.86274577 |
| *Linochilus_cinerascens* | P_cine_0035_Layer_2 | 32.5538286 | 5.02475532 |
| *Linochilus_colombianus* | P_colo_0000_Layer_26 | 5.44176716 | 2.44407523 |
| *Linochilus_colombianus* | P_colo_0001_Layer_25 | 5.93818392 | 2.57002178 |
| *Linochilus_colombianus* | P_colo_0002_Layer_24 | 5.77958145 | 2.53096502 |
| *Linochilus_colombianus* | P_colo_0003_Layer_23 | 6.89786326 | 2.78614953 |
| *Linochilus_colombianus* | P_colo_0004_Layer_22 | 8.79930075 | 3.13738888 |
| *Linochilus_colombianus* | P_colo_0005_Layer_21 | 7.2105879 | 2.85011689 |
| *Linochilus_colombianus* | P_colo_0007_Layer_20 | 8.08245344 | 3.01479329 |
| *Linochilus_colombianus* | P_colo_0008_Layer_19 | 8.04033866 | 3.00725627 |
| *Linochilus_colombianus* | P_colo_0009_Layer_18 | 8.11919187 | 3.02133614 |
| *Linochilus_colombianus* | P_colo_0013_Layer_17 | 11.3046821 | 3.49884852 |
| *Linochilus_colombianus* | P_colo_0014_Layer_16 | 13.6335398 | 3.76908829 |
| *Linochilus_colombianus* | P_colo_0016_Layer_15 | 11.6416003 | 3.54121749 |
| *Linochilus_colombianus* | P_colo_0017_Layer_14 | 4.61112033 | 2.20511732 |
| *Linochilus_colombianus* | P_colo_0018_Layer_13 | 7.20341943 | 2.84868191 |
| *Linochilus_colombianus* | P_colo_0019_Layer_12 | 4.82707059 | 2.27114792 |
| *Linochilus_colombianus* | P_colo_0020_Layer_11 | 6.16399082 | 2.62386471 |
| *Linochilus_colombianus* | P_colo_0021_Layer_10 | 7.23836573 | 2.855664 |
| *Linochilus_colombianus* | P_colo_0022_Layer_9 | 8.000016 | 3.00000289 |
| *Linochilus_colombianus* | P_colo_0025_Layer_7 | 5.92922333 | 2.56784314 |
| *Linochilus_colombianus* | P_colo_0026_Layer_6 | 8.3378303 | 3.05967201 |
| *Linochilus_colombianus* | P_colo_0027_Layer_5 | 3.93011539 | 1.97457167 |
| *Linochilus_colombianus* | P_colo_0028_Layer_4 | 8.34589483 | 3.06106674 |
| *Linochilus_colombianus* | P_colo_0029_Layer_3 | 7.12994258 | 2.83389046 |
| *Linochilus_colombianus* | P_colo_0030_Layer_1 | 10.5053974 | 3.39305883 |
| *Linochilus_coriaceus* | P_cori_0000_Layer_11 | 1636.50686 | 10.6764039 |
| *Linochilus_coriaceus* | P_cori_0001_Layer_10 | 1392.93021 | 10.4439073 |
| *Linochilus_coriaceus* | P_cori_0002_Layer_9 | 1479.44472 | 10.5308401 |
| *Linochilus_coriaceus* | P_cori_0003_Layer_8 | 2000.86959 | 10.9664114 |
| *Linochilus_coriaceus* | P_cori_0005_Layer_7 | 1903.51008 | 10.8944465 |
| *Linochilus_coriaceus* | P_cori_0006_Layer_6 | 1310.89241 | 10.3563336 |
| *Linochilus_coriaceus* | P_cori_0007_Layer_5 | 1830.36477 | 10.8379155 |
| *Linochilus_coriaceus* | P_cori_0008_Layer_4 | 1756.21677 | 10.7782552 |
| *Linochilus_coriaceus* | P_cori_0010_Layer_3 | 998.436585 | 9.96352699 |
| *Linochilus_coriaceus* | P_cori_0011_Layer_2 | 1292.5438 | 10.3359975 |
| *Linochilus_costaricensis* | P_cost_0000_Layer_25 | 79.4347467 | 6.31169831 |
| *Linochilus_costaricensis* | P_cost_0001_Layer_24 | 63.6381201 | 5.99181931 |
| *Linochilus_costaricensis* | P_cost_0002_Layer_23 | 87.682075 | 6.45421004 |
| *Linochilus_costaricensis* | P_cost_0003_Layer_22 | 62.3890137 | 5.9632201 |
| *Linochilus_costaricensis* | P_cost_0004_Layer_21 | 60.621985 | 5.92176919 |
| *Linochilus_costaricensis* | P_cost_0006_Layer_20 | 50.9078079 | 5.66981504 |
| *Linochilus_costaricensis* | P_cost_0007_Layer_19 | 57.6264593 | 5.84865948 |
| *Linochilus_costaricensis* | P_cost_0008_Layer_18 | 81.4177256 | 6.34727101 |
| *Linochilus_costaricensis* | P_cost_0009_Layer_17 | 80.4912004 | 6.33075917 |
| *Linochilus_costaricensis* | P_cost_0010_Layer_16 | 77.4580402 | 6.27534309 |
| *Linochilus_costaricensis* | P_cost_0011_Layer_15 | 132.243096 | 7.0470486 |
| *Linochilus_costaricensis* | P_cost_0013_Layer_14 | 86.590675 | 6.43613976 |
| *Linochilus_costaricensis* | P_cost_0014_Layer_13 | 138.847052 | 7.11735273 |
| *Linochilus_costaricensis* | P_cost_0015_Layer_12 | 83.2044675 | 6.37858909 |
| *Linochilus_costaricensis* | P_cost_0016_Layer_11 | 116.925861 | 6.86945024 |
| *Linochilus_costaricensis* | P_cost_0017_Layer_10 | 90.2376357 | 6.49565736 |
| *Linochilus_costaricensis* | P_cost_0018_Layer_9 | 105.367594 | 6.71928742 |
| *Linochilus_costaricensis* | P_cost_0020_Layer_8 | 85.2618193 | 6.41382793 |
| *Linochilus_costaricensis* | P_cost_0021_Layer_7 | 48.4167635 | 5.59743474 |
| *Linochilus_costaricensis* | P_cost_0022_Layer_6 | 97.7394428 | 6.61086897 |
| *Linochilus_costaricensis* | P_cost_0023_Layer_5 | 90.9571712 | 6.50711548 |
| *Linochilus_costaricensis* | P_cost_0024_Layer_4 | 52.7949085 | 5.7223269 |
| *Linochilus_costaricensis* | P_cost_0025_Layer_3 | 93.7869259 | 6.55131492 |
| *Linochilus_costaricensis* | P_cost_0028_Layer_2 | 101.418662 | 6.66417933 |
| *Linochilus_eriophorus* | P_erio_0000_Layer_27 | 35.2097478 | 5.13790299 |
| *Linochilus_eriophorus* | P_erio_0001_Layer_26 | 63.8755758 | 5.99719249 |
| *Linochilus_eriophorus* | P_erio_0002_Layer_25 | 55.2205406 | 5.78713311 |
| *Linochilus_eriophorus* | P_erio_0003_Layer_24 | 29.2294491 | 4.86935074 |
| *Linochilus_eriophorus* | P_erio_0004_Layer_23 | 62.6524551 | 5.96929914 |
| *Linochilus_eriophorus* | P_erio_0006_Layer_22 | 130.076425 | 7.0232157 |
| *Linochilus_eriophorus* | P_erio_0007_Layer_21 | 51.1864823 | 5.67769096 |
| *Linochilus_eriophorus* | P_erio_0008_Layer_20 | 49.1201699 | 5.61824365 |
| *Linochilus_eriophorus* | P_erio_0010_Layer_19 | 81.6381561 | 6.35117169 |
| *Linochilus_eriophorus* | P_erio_0012_Layer_17 | 83.7618163 | 6.38822082 |
| *Linochilus_eriophorus* | P_erio_0013_Layer_16 | 77.5691516 | 6.27741112 |
| *Linochilus_eriophorus* | P_erio_0014_Layer_15 | 134.642742 | 7.07299266 |
| *Linochilus_eriophorus* | P_erio_0015_Layer_14 | 88.439245 | 6.4666148 |
| *Linochilus_eriophorus* | P_erio_0016_Layer_13 | 130.496677 | 7.02786926 |
| *Linochilus_eriophorus* | P_erio_0018_Layer_12 | 138.00386 | 7.10856481 |
| *Linochilus_eriophorus* | P_erio_0019_Layer_11 | 87.3908558 | 6.44941042 |
| *Linochilus_eriophorus* | P_erio_0020_Layer_10 | 111.071907 | 6.79535015 |
| *Linochilus_eriophorus* | P_erio_0021_Layer_9 | 90.5575288 | 6.50076268 |
| *Linochilus_eriophorus* | P_erio_0023_Layer_8 | 112.12388 | 6.80894977 |
| *Linochilus_eriophorus* | P_erio_0024_Layer_7 | 115.00471 | 6.84554914 |
| *Linochilus_eriophorus* | P_erio_0025_Layer_6 | 135.666042 | 7.08391584 |
| *Linochilus_eriophorus* | P_erio_0026_Layer_5 | 106.7305 | 6.7378287 |
| *Linochilus_eriophorus* | P_erio_0027_Layer_4 | 135.849734 | 7.08586793 |
| *Linochilus_eriophorus* | P_erio_0028_Layer_3 | 171.394608 | 7.42117791 |
| *Linochilus_eriophorus* | P_erio_0029_Layer_2 | 106.470643 | 6.73431188 |
| *Linochilus_floribundus* | P_flor_0000_Layer_38 | 113.138219 | 6.82194256 |
| *Linochilus_floribundus* | P_flor_0001_Layer_37 | 248.586519 | 7.95760425 |
| *Linochilus_floribundus* | P_flor_0002_Layer_36 | 299.948629 | 8.22857163 |
| *Linochilus_floribundus* | P_flor_0003_Layer_35 | 155.946549 | 7.28490781 |
| *Linochilus_floribundus* | P_flor_0004_Layer_34 | 283.391248 | 8.14665139 |
| *Linochilus_floribundus* | P_flor_0006_Layer_33 | 254.534559 | 7.99171774 |
| *Linochilus_floribundus* | P_flor_0007_Layer_31 | 210.341819 | 7.71659189 |
| *Linochilus_floribundus* | P_flor_0008_Layer_30 | 103.993936 | 6.70035559 |
| *Linochilus_floribundus* | P_flor_0009_Layer_26 | 343.696924 | 8.42499313 |
| *Linochilus_floribundus* | P_flor_0011_Layer_29 | 149.771805 | 7.22662225 |
| *Linochilus_floribundus* | P_flor_0012_Layer_32 | 148.551372 | 7.21481812 |
| *Linochilus_floribundus* | P_flor_0013_Layer_28 | 243.211956 | 7.92607034 |
| *Linochilus_floribundus* | P_flor_0014_Layer_27 | 82.2627093 | 6.36216668 |
| *Linochilus_floribundus* | P_flor_0015_Layer_25 | 307.544522 | 8.26465147 |
| *Linochilus_floribundus* | P_flor_0016_Layer_24 | 442.152318 | 8.78839964 |
| *Linochilus_floribundus* | P_flor_0017_Layer_23 | 292.389474 | 8.19174756 |
| *Linochilus_floribundus* | P_flor_0018_Layer_22 | 315.024824 | 8.29932171 |
| *Linochilus_floribundus* | P_flor_0020_Layer_21 | 384.727472 | 8.58769304 |
| *Linochilus_floribundus* | P_flor_0021_Layer_20 | 183.864167 | 7.52249653 |
| *Linochilus_floribundus* | P_flor_0022_Layer_19 | 241.894749 | 7.91823564 |
| *Linochilus_floribundus* | P_flor_0023_Layer_18 | 195.294297 | 7.60950601 |
| *Linochilus_floribundus* | P_flor_0024_Layer_17 | 167.829189 | 7.39084984 |
| *Linochilus_floribundus* | P_flor_0025_Layer_16 | 344.549076 | 8.42856568 |
| *Linochilus_floribundus* | P_flor_0026_Layer_11 | 205.048797 | 7.67982347 |
| *Linochilus_floribundus* | P_flor_0027_Layer_10 | 332.696902 | 8.37806462 |
| *Linochilus_floribundus* | P_flor_0029_Layer_15 | 318.008701 | 8.31292243 |
| *Linochilus_floribundus* | P_flor_0031_Layer_14 | 270.21201 | 8.07794799 |
| *Linochilus_floribundus* | P_flor_0032_Layer_13 | 489.732162 | 8.93584914 |
| *Linochilus_floribundus* | P_flor_0033_Layer_12 | 344.830439 | 8.42974332 |
| *Linochilus_floribundus* | P_flor_0034_Layer_9 | 237.029148 | 7.88892067 |
| *Linochilus_floribundus* | P_flor_0035_Layer_8 | 173.589953 | 7.43953964 |
| *Linochilus_floribundus* | P_flor_0036_Layer_7 | 334.946907 | 8.38778862 |
| *Linochilus_floribundus* | P_flor_0037_Layer_6 | 275.30521 | 8.1048881 |
| *Linochilus_floribundus* | P_flor_0038_Layer_5 | 303.414585 | 8.24514663 |
| *Linochilus_floribundus* | P_flor_0039_Layer_4 | 179.977063 | 7.49166924 |
| *Linochilus_floribundus* | P_flor_0044_Layer_1 | 543.3389 | 9.08570853 |
| *Linochilus_frontinensis* | P_fron_0000_Layer_4 | 67.4194897 | 6.0750938 |
| *Linochilus_frontinensis* | P_fron_0001_Layer_14 | 50.590603 | 5.66079753 |
| *Linochilus_frontinensis* | P_fron_0002_Layer_13 | 77.000154 | 6.26678943 |
| *Linochilus_frontinensis* | P_fron_0003_Layer_12 | 81.2519546 | 6.34433061 |
| *Linochilus_frontinensis* | P_fron_0004_Layer_11 | 83.2465823 | 6.37931914 |
| *Linochilus_frontinensis* | P_fron_0005_Layer_10 | 79.1300866 | 6.30615443 |
| *Linochilus_frontinensis* | P_fron_0006_Layer_9 | 104.352359 | 6.70531941 |
| *Linochilus_frontinensis* | P_fron_0007_Layer_8 | 86.6050119 | 6.43637861 |
| *Linochilus_frontinensis* | P_fron_0009_Layer_7 | 61.4804097 | 5.94205487 |
| *Linochilus_frontinensis* | P_fron_0010_Layer_6 | 43.8522382 | 5.45457858 |
| *Linochilus_frontinensis* | P_fron_0011_Layer_5 | 54.8397154 | 5.77714918 |
| *Linochilus_frontinensis* | P_fron_0012_Layer_3 | 62.6354299 | 5.96890705 |
| *Linochilus_frontinensis* | P_fron_0016_Layer_1 | 50.7277 | 5.66470185 |
| *Linochilus_glutinosus* | P_glut_0000_Layer_19 | 56.9929455 | 5.83271145 |
| *Linochilus_glutinosus* | P_glut_0001_Layer_33 | 50.9382739 | 5.67067817 |
| *Linochilus_glutinosus* | P_glut_0002_Layer_32 | 59.940084 | 5.9054492 |
| *Linochilus_glutinosus* | P_glut_0003_Layer_30 | 59.5897249 | 5.89699168 |
| *Linochilus_glutinosus* | P_glut_0005_Layer_29 | 45.8361132 | 5.51841281 |
| *Linochilus_glutinosus* | P_glut_0006_Layer_28 | 16.3037961 | 4.027136 |
| *Linochilus_glutinosus* | P_glut_0007_Layer_27 | 21.0314041 | 4.39447326 |
| *Linochilus_glutinosus* | P_glut_0008_Layer_26 | 17.6201069 | 4.13915077 |
| *Linochilus_glutinosus* | P_glut_0009_Layer_25 | 15.5708197 | 3.96077299 |
| *Linochilus_glutinosus* | P_glut_0010_Layer_24 | 17.1523641 | 4.10033553 |
| *Linochilus_glutinosus* | P_glut_0013_Layer_22 | 76.545852 | 6.2582523 |
| *Linochilus_glutinosus* | P_glut_0014_Layer_21 | 45.6793028 | 5.51346873 |
| *Linochilus_glutinosus* | P_glut_0015_Layer_20 | 48.9293093 | 5.61262701 |
| *Linochilus_glutinosus* | P_glut_0016_Layer_18 | 51.6076301 | 5.68951248 |
| *Linochilus_glutinosus* | P_glut_0018_Layer_16 | 15.9005694 | 3.99100653 |
| *Linochilus_glutinosus* | P_glut_0019_Layer_15 | 11.3055782 | 3.49896287 |
| *Linochilus_glutinosus* | P_glut_0020_Layer_14 | 16.7043345 | 4.0621506 |
| *Linochilus_glutinosus* | P_glut_0021_Layer_13 | 13.6873034 | 3.77476633 |
| *Linochilus_glutinosus* | P_glut_0022_Layer_12 | 11.7975146 | 3.56041106 |
| *Linochilus_glutinosus* | P_glut_0024_Layer_11 | 39.4283943 | 5.30116305 |
| *Linochilus_glutinosus* | P_glut_0025_Layer_10 | 48.3208852 | 5.59457498 |
| *Linochilus_glutinosus* | P_glut_0026_Layer_9 | 46.2868309 | 5.53252989 |
| *Linochilus_glutinosus* | P_glut_0027_Layer_8 | 39.2751682 | 5.29554555 |
| *Linochilus_glutinosus* | P_glut_0029_Layer_7 | 43.8119156 | 5.45325139 |
| *Linochilus_glutinosus* | P_glut_0030_Layer_6 | 27.5681555 | 4.78493084 |
| *Linochilus_glutinosus* | P_glut_0031_Layer_5 | 54.8173139 | 5.77655973 |
| *Linochilus_glutinosus* | P_glut_0032_Layer_4 | 43.4257141 | 5.44047767 |
| *Linochilus_glutinosus* | P_glut_0033_Layer_3 | 24.9534549 | 4.64116767 |
| *Linochilus_glutinosus* | P_glut_0034_Layer_2 | 16.1577384 | 4.01415337 |
| *Linochilus_heterophyllus* | P_hete_0000_Layer_32 | 21.1550602 | 4.40293089 |
| *Linochilus_heterophyllus* | P_hete_0001_Layer_31 | 18.0045163 | 4.17028694 |
| *Linochilus_heterophyllus* | P_hete_0002_Layer_30 | 9.60933822 | 3.26443708 |
| *Linochilus_heterophyllus* | P_hete_0003_Layer_29 | 17.3575616 | 4.11749239 |
| *Linochilus_heterophyllus* | P_hete_0005_Layer_28 | 16.5251227 | 4.04658908 |
| *Linochilus_heterophyllus* | P_hete_0006_Layer_27 | 14.6425024 | 3.87209023 |
| *Linochilus_heterophyllus* | P_hete_0007_Layer_26 | 15.7043325 | 3.97309072 |
| *Linochilus_heterophyllus* | P_hete_0008_Layer_25 | 8.43729286 | 3.07678018 |
| *Linochilus_heterophyllus* | P_hete_0009_Layer_24 | 10.9704521 | 3.45555107 |
| *Linochilus_heterophyllus* | P_hete_0010_Layer_23 | 8.56274114 | 3.09807271 |
| *Linochilus_heterophyllus* | P_hete_0012_Layer_22 | 19.3853434 | 4.27689439 |
| *Linochilus_heterophyllus* | P_hete_0013_Layer_21 | 10.5573688 | 3.40017841 |
| *Linochilus_heterophyllus* | P_hete_0014_Layer_20 | 30.0475511 | 4.90917551 |
| *Linochilus_heterophyllus* | P_hete_0015_Layer_19 | 25.0332042 | 4.64577106 |
| *Linochilus_heterophyllus* | P_hete_0016_Layer_18 | 30.1559743 | 4.91437194 |
| *Linochilus_heterophyllus* | P_hete_0018_Layer_17 | 15.2679517 | 3.93243462 |
| *Linochilus_heterophyllus* | P_hete_0019_Layer_16 | 11.773321 | 3.55744943 |
| *Linochilus_heterophyllus* | P_hete_0020_Layer_15 | 15.3853354 | 3.94348399 |
| *Linochilus_heterophyllus* | P_hete_0021_Layer_14 | 16.2034374 | 4.018228 |
| *Linochilus_heterophyllus* | P_hete_0022_Layer_13 | 15.9122182 | 3.99206306 |
| *Linochilus_heterophyllus* | P_hete_0024_Layer_12 | 16.5170581 | 4.04588484 |
| *Linochilus_heterophyllus* | P_hete_0025_Layer_11 | 18.6747685 | 4.22301846 |
| *Linochilus_heterophyllus* | P_hete_0026_Layer_10 | 26.8298028 | 4.74576454 |
| *Linochilus_heterophyllus* | P_hete_0027_Layer_9 | 18.9005754 | 4.24035825 |
| *Linochilus_heterophyllus* | P_hete_0028_Layer_8 | 18.3199291 | 4.19534202 |
| *Linochilus_heterophyllus* | P_hete_0030_Layer_7 | 14.6353339 | 3.87138376 |
| *Linochilus_heterophyllus* | P_hete_0032_Layer_5 | 25.621915 | 4.6793064 |
| *Linochilus_heterophyllus* | P_hete_0033_Layer_6 | 14.2043295 | 3.82825883 |
| *Linochilus_heterophyllus* | P_hete_0034_Layer_4 | 22.5412637 | 4.49449649 |
| *Linochilus_heterophyllus* | P_hete_0035_Layer_3 | 12.0797733 | 3.59452147 |
| *Linochilus_heterophyllus* | P_hete_0038_Layer_1 | 22.1021947 | 4.46611773 |
| *Linochilus_huertasii* | P_huer_0000_Layer_25 | 1319.08239 | 10.365319 |
| *Linochilus_huertasii* | P_huer_0001_Layer_24 | 2294.96068 | 11.1642537 |
| *Linochilus_huertasii* | P_huer_0002_Layer_23 | 2901.32301 | 11.5024952 |
| *Linochilus_huertasii* | P_huer_0004_Layer_22 | 3683.46346 | 11.8468472 |
| *Linochilus_huertasii* | P_huer_0005_Layer_21 | 1727.98554 | 10.7548754 |
| *Linochilus_huertasii* | P_huer_0006_Layer_20 | 1726.16654 | 10.7533559 |
| *Linochilus_huertasii* | P_huer_0007_Layer_19 | 2844.28347 | 11.4738495 |
| *Linochilus_huertasii* | P_huer_0008_Layer_18 | 746.380525 | 9.54376753 |
| *Linochilus_huertasii* | P_huer_0009_Layer_17 | 1151.8679 | 10.1697596 |
| *Linochilus_huertasii* | P_huer_0012_Layer_9 | 3050.86811 | 11.5750041 |
| *Linochilus_huertasii* | P_huer_0013_Layer_15 | 238.420728 | 7.89736586 |
| *Linochilus_huertasii* | P_huer_0014_Layer_14 | 608.132937 | 9.24824292 |
| *Linochilus_huertasii* | P_huer_0015_Layer_13 | 607.278096 | 9.24621352 |
| *Linochilus_huertasii* | P_huer_0016_Layer_12 | 313.569624 | 8.292642 |
| *Linochilus_huertasii* | P_huer_0017_Layer_11 | 586.365868 | 9.19565732 |
| *Linochilus_huertasii* | P_huer_0020_Layer_8 | 731.609886 | 9.51493076 |
| *Linochilus_huertasii* | P_huer_0021_Layer_7 | 553.660606 | 9.11285806 |
| *Linochilus_huertasii* | P_huer_0022_Layer_6 | 437.226681 | 8.77223763 |
| *Linochilus_huertasii* | P_huer_0023_Layer_5 | 324.064268 | 8.34013615 |
| *Linochilus_huertasii* | P_huer_0024_Layer_4 | 314.201345 | 8.29554555 |
| *Linochilus_huertasii* | P_huer_0025_Layer_3 | 1194.19056 | 10.2218174 |
| *Linochilus_inesianus* | P_ines_0000_Layer_6 | 15.4211778 | 3.94684105 |
| *Linochilus_inesianus* | P_ines_0001_Layer_5 | 14.5735059 | 3.86527607 |
| *Linochilus_inesianus* | P_ines_0002_Layer_4 | 16.4650867 | 4.0413382 |
| *Linochilus_inesianus* | P_ines_0003_Layer_3 | 19.9901834 | 4.3212198 |
| *Linochilus_inesianus* | P_ines_0004_Layer_15 | 22.1496859 | 4.46921433 |
| *Linochilus_inesianus* | P_ines_0005_Layer_16 | 17.9794266 | 4.16827511 |
| *Linochilus_inesianus* | P_ines_0006_Layer_14 | 23.0815874 | 4.52867054 |
| *Linochilus_inesianus* | P_ines_0007_Layer_13 | 23.1416233 | 4.53241817 |
| *Linochilus_inesianus* | P_ines_0008_Layer_12 | 18.8360592 | 4.23542526 |
| *Linochilus_inesianus* | P_ines_0010_Layer_11 | 29.6559733 | 4.89025082 |
| *Linochilus_inesianus* | P_ines_0011_Layer_10 | 38.272478 | 5.25823541 |
| *Linochilus_inesianus* | P_ines_0012_Layer_9 | 40.9660318 | 5.35635625 |
| *Linochilus_inesianus* | P_ines_0013_Layer_8 | 28.363856 | 4.82598177 |
| *Linochilus_inesianus* | P_ines_0014_Layer_7 | 38.3764208 | 5.26214826 |
| *Linochilus_inesianus* | P_ines_0019_Layer_1 | 11.8557585 | 3.56751606 |
| *Linochilus_jaramilloi* | P_jara_0000_Layer_11 | 1126.63218 | 10.1378009 |
| *Linochilus_jaramilloi* | P_jara_0001_Layer_6 | 7106.73285 | 12.7949708 |
| *Linochilus_jaramilloi* | P_jara_0002_Layer_5 | 1621.45127 | 10.66307 |
| *Linochilus_jaramilloi* | P_jara_0003_Layer_9 | 4780.19056 | 12.2228524 |
| *Linochilus_jaramilloi* | P_jara_0004_Layer_10 | 2919.55423 | 11.5115324 |
| *Linochilus_jaramilloi* | P_jara_0005_Layer_8 | 3043.74444 | 11.5716315 |
| *Linochilus_jaramilloi* | P_jara_0006_Layer_7 | 2369.4447 | 11.2103333 |
| *Linochilus_jaramilloi* | P_jara_0008_Layer_4 | 1352.31364 | 10.4012141 |
| *Linochilus_jaramilloi* | P_jara_0009_Layer_3 | 1476.23593 | 10.5277076 |
| *Linochilus_jaramilloi* | P_jara_0010_Layer_2 | 7973.3448 | 12.9609693 |
| *Linochilus_jenesanus* | P_jene_0003_Layer_14 | 698.198529 | 9.44749351 |
| *Linochilus_jenesanus* | P_jene_0004_Layer_13 | 520.053908 | 9.02251737 |
| *Linochilus_jenesanus* | P_jene_0005_Layer_12 | 584.542388 | 9.19116383 |
| *Linochilus_jenesanus* | P_jene_0006_Layer_11 | 710.245148 | 9.47217326 |
| *Linochilus_jenesanus* | P_jene_0008_Layer_3 | 551.12207 | 9.10622809 |
| *Linochilus_jenesanus* | P_jene_0012_Layer_4 | 1163.60985 | 10.1843917 |
| *Linochilus_jenesanus* | P_jene_0014_Layer_9 | 1072.3982 | 10.066625 |
| *Linochilus_jenesanus* | P_jene_0015_Layer_8 | 1174.56507 | 10.1979109 |
| *Linochilus_jenesanus* | P_jene_0017_Layer_6 | 1257.58137 | 10.296436 |
| *Linochilus_juajibioyi* | P_juaj_0000_Layer_29 | 255.645673 | 7.9980018 |
| *Linochilus_juajibioyi* | P_juaj_0001_Layer_28 | 174.502141 | 7.44710093 |
| *Linochilus_juajibioyi* | P_juaj_0002_Layer_27 | 321.666414 | 8.3294215 |
| *Linochilus_juajibioyi* | P_juaj_0003_Layer_26 | 358.810754 | 8.48707932 |
| *Linochilus_juajibioyi* | P_juaj_0005_Layer_25 | 267.564155 | 8.06374104 |
| *Linochilus_juajibioyi* | P_juaj_0006_Layer_24 | 87.0136149 | 6.44316925 |
| *Linochilus_juajibioyi* | P_juaj_0007_Layer_23 | 188.952887 | 7.56188275 |
| *Linochilus_juajibioyi* | P_juaj_0008_Layer_22 | 157.227017 | 7.29670533 |
| *Linochilus_juajibioyi* | P_juaj_0009_Layer_21 | 156.968056 | 7.29432718 |
| *Linochilus_juajibioyi* | P_juaj_0010_Layer_20 | 163.660722 | 7.35456431 |
| *Linochilus_juajibioyi* | P_juaj_0012_Layer_19 | 137.97429 | 7.10825565 |
| *Linochilus_juajibioyi* | P_juaj_0013_Layer_4 | 250.05516 | 7.96610256 |
| *Linochilus_juajibioyi* | P_juaj_0014_Layer_15 | 191.690348 | 7.58263388 |
| *Linochilus_juajibioyi* | P_juaj_0015_Layer_16 | 124.688421 | 6.96218369 |
| *Linochilus_juajibioyi* | P_juaj_0017_Layer_18 | 263.692284 | 8.04271154 |
| *Linochilus_juajibioyi* | P_juaj_0018_Layer_17 | 246.623253 | 7.94616502 |
| *Linochilus_juajibioyi* | P_juaj_0019_Layer_14 | 61.9517368 | 5.95307282 |
| *Linochilus_juajibioyi* | P_juaj_0020_Layer_13 | 61.9535289 | 5.95311456 |
| *Linochilus_juajibioyi* | P_juaj_0021_Layer_12 | 246.23526 | 7.94389355 |
| *Linochilus_juajibioyi* | P_juaj_0022_Layer_11 | 154.578266 | 7.27219368 |
| *Linochilus_juajibioyi* | P_juaj_0023_Layer_10 | 261.447656 | 8.03037832 |
| *Linochilus_juajibioyi* | P_juaj_0026_Layer_8 | 127.995776 | 6.99995239 |
| *Linochilus_juajibioyi* | P_juaj_0028_Layer_6 | 52.1004626 | 5.70322428 |
| *Linochilus_juajibioyi* | P_juaj_0029_Layer_5 | 189.615971 | 7.56693667 |
| *Linochilus_juajibioyi* | P_juaj_0030_Layer_3 | 202.165279 | 7.65939143 |
| *Linochilus_juajibioyi* | P_juaj_0031_Layer_2 | 209.248626 | 7.70907434 |
| *Linochilus_lacunosus* | P_lacu_0000_Layer_13 | 16.4041547 | 4.03598935 |
| *Linochilus_lacunosus* | P_lacu_0001_Layer_12 | 27.848622 | 4.79953404 |
| *Linochilus_lacunosus* | P_lacu_0002_Layer_11 | 25.0009461 | 4.64391078 |
| *Linochilus_lacunosus* | P_lacu_0003_Layer_10 | 26.2751422 | 4.71562666 |
| *Linochilus_lacunosus* | P_lacu_0004_Layer_9 | 35.2464863 | 5.13940754 |
| *Linochilus_lacunosus* | P_lacu_0005_Layer_8 | 27.000054 | 4.75489039 |
| *Linochilus_lacunosus* | P_lacu_0007_Layer_7 | 20.8862425 | 4.38448106 |
| *Linochilus_lacunosus* | P_lacu_0008_Layer_6 | 21.8423376 | 4.44905536 |
| *Linochilus_lacunosus* | P_lacu_0009_Layer_5 | 21.6299716 | 4.43495986 |
| *Linochilus_lacunosus* | P_lacu_0010_Layer_4 | 22.0896499 | 4.46529865 |
| *Linochilus_lacunosus* | P_lacu_0011_Layer_3 | 12.9301334 | 3.69266525 |
| *Linochilus_lacunosus* | P_lacu_0012_Layer_2 | 25.2473623 | 4.65806077 |
| *Linochilus_mutiscuanus* | P_muti_0000_Layer_10 | 2025.17341 | 10.9838297 |
| *Linochilus_mutiscuanus* | P_muti_0001_Layer_9 | 2221.0367 | 11.1170175 |
| *Linochilus_mutiscuanus* | P_muti_0002_Layer_8 | 2205.03846 | 11.1065881 |
| *Linochilus_mutiscuanus* | P_muti_0004_Layer_7 | 2948.44765 | 11.5257399 |
| *Linochilus_mutiscuanus* | P_muti_0007_Layer_4 | 2646.68003 | 11.3699681 |
| *Linochilus_mutiscuanus* | P_muti_0008_Layer_3 | 2204.55459 | 11.1062715 |
| *Linochilus_mutiscuanus* | P_muti_0009_Layer_2 | 2485.94404 | 11.2795781 |
| *Linochilus_oblongifolius* | P_oblo_0000_Layer_8 | 1895.37207 | 10.8882654 |
| *Linochilus_oblongifolius* | P_oblo_0001_Layer_10 | 2182.73555 | 11.0919216 |
| *Linochilus_oblongifolius* | P_oblo_0002_Layer_9 | 1246.16109 | 10.2832749 |
| *Linochilus_oblongifolius* | P_oblo_0003_Layer_6 | 1532.99052 | 10.5821331 |
| *Linochilus_oblongifolius* | P_oblo_0005_Layer_5 | 891.392464 | 9.79991695 |
| *Linochilus_oblongifolius* | P_oblo_0006_Layer_4 | 760.580374 | 9.5709569 |
| *Linochilus_oblongifolius* | P_oblo_0007_Layer_3 | 1174.05074 | 10.197279 |
| *Linochilus_oblongifolius* | P_oblo_0008_Layer_2 | 1116.53987 | 10.124819 |
| *Linochilus_obtusus* | P_obtu_0000_Layer_31 | 147.515528 | 7.20472302 |
| *Linochilus_obtusus* | P_obtu_0001_Layer_30 | 177.632076 | 7.47274831 |
| *Linochilus_obtusus* | P_obtu_0002_Layer_29 | 210.073001 | 7.71474694 |
| *Linochilus_obtusus* | P_obtu_0003_Layer_28 | 61.8020949 | 5.94958384 |
| *Linochilus_obtusus* | P_obtu_0004_Layer_27 | 96.3344221 | 6.58997949 |
| *Linochilus_obtusus* | P_obtu_0006_Layer_26 | 227.662642 | 7.83075376 |
| *Linochilus_obtusus* | P_obtu_0007_Layer_25 | 176.720784 | 7.46532791 |
| *Linochilus_obtusus* | P_obtu_0008_Layer_24 | 377.959537 | 8.56208798 |
| *Linochilus_obtusus* | P_obtu_0009_Layer_23 | 286.660968 | 8.16320167 |
| *Linochilus_obtusus* | P_obtu_0010_Layer_22 | 428.858385 | 8.74435752 |
| *Linochilus_obtusus* | P_obtu_0011_Layer_21 | 413.684519 | 8.69238716 |
| *Linochilus_obtusus* | P_obtu_0012_Layer_20 | 266.889423 | 8.06009832 |
| *Linochilus_obtusus* | P_obtu_0013_Layer_19 | 84.969704 | 6.40887663 |
| *Linochilus_obtusus* | P_obtu_0014_Layer_18 | 117.884644 | 6.881232 |
| *Linochilus_obtusus* | P_obtu_0015_Layer_17 | 228.599024 | 7.83667543 |
| *Linochilus_obtusus* | P_obtu_0017_Layer_16 | 407.638808 | 8.6711476 |
| *Linochilus_obtusus* | P_obtu_0019_Layer_15 | 277.470985 | 8.11619311 |
| *Linochilus_obtusus* | P_obtu_0020_Layer_13 | 87.6166627 | 6.45313336 |
| *Linochilus_obtusus* | P_obtu_0021_Layer_14 | 97.2708039 | 6.60393494 |
| *Linochilus_obtusus* | P_obtu_0022_Layer_12 | 72.9177086 | 6.18819732 |
| *Linochilus_obtusus* | P_obtu_0023_Layer_11 | 83.7331424 | 6.38772686 |
| *Linochilus_obtusus* | P_obtu_0024_Layer_10 | 49.2814606 | 5.62297311 |
| *Linochilus_obtusus* | P_obtu_0026_Layer_9 | 45.5807363 | 5.51035233 |
| *Linochilus_obtusus* | P_obtu_0028_Layer_7 | 200.257569 | 7.64571296 |
| *Linochilus_obtusus* | P_obtu_0029_Layer_6 | 219.753128 | 7.77973989 |
| *Linochilus_obtusus* | P_obtu_0030_Layer_5 | 200.254881 | 7.6456936 |
| *Linochilus_obtusus* | P_obtu_0031_Layer_4 | 96.3675763 | 6.59047592 |
| *Linochilus_obtusus* | P_obtu_0032_Layer_3 | 85.2035754 | 6.41284207 |
| *Linochilus_obtusus* | P_obtu_0033_Layer_2 | 320.872506 | 8.32585636 |
| *Linochilus_ochraceus* | P_ochr_0000_Layer_22 | 1497.92683 | 10.5487514 |
| *Linochilus_ochraceus* | P_ochr_0001_Layer_21 | 1537.95738 | 10.5867998 |
| *Linochilus_ochraceus* | P_ochr_0002_Layer_20 | 976.480448 | 9.93144735 |
| *Linochilus_ochraceus* | P_ochr_0004_Layer_19 | 517.328096 | 9.01493573 |
| *Linochilus_ochraceus* | P_ochr_0005_Layer_18 | 621.54963 | 9.27972578 |
| *Linochilus_ochraceus* | P_ochr_0006_Layer_17 | 954.814634 | 9.89907687 |
| *Linochilus_ochraceus* | P_ochr_0007_Layer_16 | 815.837653 | 9.67213828 |
| *Linochilus_ochraceus* | P_ochr_0009_Layer_12 | 1456.78965 | 10.5085769 |
| *Linochilus_ochraceus* | P_ochr_0010_Layer_11 | 1429.1525 | 10.4809442 |
| *Linochilus_ochraceus* | P_ochr_0012_Layer_15 | 1750.29651 | 10.7733836 |
| *Linochilus_ochraceus* | P_ochr_0013_Layer_14 | 1965.77454 | 10.9408821 |
| *Linochilus_ochraceus* | P_ochr_0015_Layer_13 | 2638.53754 | 11.3655228 |
| *Linochilus_ochraceus* | P_ochr_0016_Layer_10 | 1766.35568 | 10.7865602 |
| *Linochilus_ochraceus* | P_ochr_0018_Layer_7 | 1607.3473 | 10.650466 |
| *Linochilus_ochraceus* | P_ochr_0019_Layer_8 | 1121.66712 | 10.1314289 |
| *Linochilus_ochraceus* | P_ochr_0020_Layer_6 | 976.016289 | 9.93076142 |
| *Linochilus_ochraceus* | P_ochr_0021_Layer_5 | 1872.38099 | 10.8706583 |
| *Linochilus_ochraceus* | P_ochr_0023_Layer_4 | 1612.23441 | 10.6548458 |
| *Linochilus_ochraceus* | P_ochr_0024_Layer_3 | 1508.40355 | 10.5588067 |
| *Linochilus_ochraceus* | P_ochr_0025_Layer_2 | 984.172219 | 9.94276698 |
| *Linochilus_phylicoides* | P_phyl_0001_Layer_34 | 12.5170501 | 3.6458227 |
| *Linochilus_phylicoides* | P_phyl_0002_Layer_33 | 12.7383767 | 3.67110954 |
| *Linochilus_phylicoides* | P_phyl_0003_Layer_32 | 8.57438991 | 3.10003402 |
| *Linochilus_phylicoides* | P_phyl_0004_Layer_31 | 11.7715289 | 3.55722981 |
| *Linochilus_phylicoides* | P_phyl_0006_Layer_30 | 15.196267 | 3.92564506 |
| *Linochilus_phylicoides* | P_phyl_0007_Layer_29 | 17.2097118 | 4.10515104 |
| *Linochilus_phylicoides* | P_phyl_0008_Layer_28 | 18.6272774 | 4.21934492 |
| *Linochilus_phylicoides* | P_phyl_0009_Layer_27 | 23.1452076 | 4.5326416 |
| *Linochilus_phylicoides* | P_phyl_0010_Layer_26 | 18.0081005 | 4.17057411 |
| *Linochilus_phylicoides* | P_phyl_0011_Layer_25 | 15.7392788 | 3.97629753 |
| *Linochilus_phylicoides* | P_phyl_0014_Layer_23 | 13.6953679 | 3.77561612 |
| *Linochilus_phylicoides* | P_phyl_0015_Layer_22 | 23.4901904 | 4.5539865 |
| *Linochilus_phylicoides* | P_phyl_0016_Layer_21 | 18.8817582 | 4.2389212 |
| *Linochilus_phylicoides* | P_phyl_0017_Layer_20 | 19.2374937 | 4.26584895 |
| *Linochilus_phylicoides* | P_phyl_0019_Layer_19 | 20.5950233 | 4.36422385 |
| *Linochilus_phylicoides* | P_phyl_0020_Layer_18 | 10.3396264 | 3.37011215 |
| *Linochilus_phylicoides* | P_phyl_0021_Layer_17 | 8.10127068 | 3.01814821 |
| *Linochilus_phylicoides* | P_phyl_0022_Layer_16 | 11.2894491 | 3.49690318 |
| *Linochilus_phylicoides* | P_phyl_0023_Layer_15 | 11.2822806 | 3.49598682 |
| *Linochilus_phylicoides* | P_phyl_0024_Layer_14 | 9.5878328 | 3.26120475 |
| *Linochilus_phylicoides* | P_phyl_0025_Layer_13 | 16.5206424 | 4.04619788 |
| *Linochilus_phylicoides* | P_phyl_0026_Layer_12 | 16.3772729 | 4.03362324 |
| *Linochilus_phylicoides* | P_phyl_0027_Layer_11 | 12.530491 | 3.64737104 |
| *Linochilus_phylicoides* | P_phyl_0028_Layer_10 | 15.2186684 | 3.92777023 |
| *Linochilus_phylicoides* | P_phyl_0030_Layer_9 | 30.21153 | 4.91702734 |
| *Linochilus_phylicoides* | P_phyl_0033_Layer_6 | 19.4606124 | 4.28248521 |
| *Linochilus_phylicoides* | P_phyl_0034_Layer_5 | 17.7106089 | 4.14654191 |
| *Linochilus_phylicoides* | P_phyl_0035_Layer_4 | 31.2670876 | 4.96657294 |
| *Linochilus_phylicoides* | P_phyl_0036_Layer_3 | 18.4149114 | 4.20280255 |
| *Linochilus_phylicoides* | P_phyl_0037_Layer_2 | 21.6720864 | 4.43776614 |
| *Linochilus_revolutus* | P_revo_0000_Layer_32 | 13.2061196 | 3.72313471 |
| *Linochilus_revolutus* | P_revo_0001_Layer_31 | 16.9677759 | 4.08472556 |
| *Linochilus_revolutus* | P_revo_0002_Layer_30 | 15.3199231 | 3.93733715 |
| *Linochilus_revolutus* | P_revo_0003_Layer_29 | 13.545726 | 3.75976582 |
| *Linochilus_revolutus* | P_revo_0005_Layer_28 | 14.3279857 | 3.84076389 |
| *Linochilus_revolutus* | P_revo_0006_Layer_27 | 12.3423186 | 3.62554154 |
| *Linochilus_revolutus* | P_revo_0007_Layer_26 | 12.7052225 | 3.66734974 |
| *Linochilus_revolutus* | P_revo_0008_Layer_25 | 12.8145418 | 3.67970998 |
| *Linochilus_revolutus* | P_revo_0009_Layer_24 | 14.6487748 | 3.8727081 |
| *Linochilus_revolutus* | P_revo_0010_Layer_23 | 12.2831787 | 3.61861205 |
| *Linochilus_revolutus* | P_revo_0011_Layer_20 | 14.3969822 | 3.84769453 |
| *Linochilus_revolutus* | P_revo_0013_Layer_22 | 19.0170631 | 4.24922256 |
| *Linochilus_revolutus* | P_revo_0014_Layer_21 | 16.424764 | 4.03780074 |
| *Linochilus_revolutus* | P_revo_0015_Layer_19 | 15.5367694 | 3.95761465 |
| *Linochilus_revolutus* | P_revo_0017_Layer_18 | 20.6030878 | 4.36478867 |
| *Linochilus_revolutus* | P_revo_0018_Layer_17 | 36.8611848 | 5.20403054 |
| *Linochilus_revolutus* | P_revo_0019_Layer_16 | 50.2348675 | 5.65061717 |
| *Linochilus_revolutus* | P_revo_0020_Layer_15 | 40.0081445 | 5.32222182 |
| *Linochilus_revolutus* | P_revo_0021_Layer_14 | 45.4069009 | 5.50483967 |
| *Linochilus_revolutus* | P_revo_0023_Layer_13 | 20.6846292 | 4.37048719 |
| *Linochilus_revolutus* | P_revo_0024_Layer_12 | 17.4803217 | 4.12765983 |
| *Linochilus_revolutus* | P_revo_0025_Layer_11 | 20.1407213 | 4.33204345 |
| *Linochilus_revolutus* | P_revo_0026_Layer_10 | 31.8835763 | 4.99474156 |
| *Linochilus_revolutus* | P_revo_0027_Layer_9 | 23.9014815 | 4.57902814 |
| *Linochilus_revolutus* | P_revo_0029_Layer_8 | 27.1783698 | 4.76438702 |
| *Linochilus_revolutus* | P_revo_0031_Layer_6 | 11.1039649 | 3.473003 |
| *Linochilus_revolutus* | P_revo_0032_Layer_5 | 9.16937318 | 3.19682311 |
| *Linochilus_revolutus* | P_revo_0033_Layer_4 | 8.90593179 | 3.15476656 |
| *Linochilus_revolutus* | P_revo_0034_Layer_3 | 7.13621499 | 2.83515908 |
| *Linochilus_revolutus* | P_revo_0035_Layer_2 | 12.3951861 | 3.63170802 |
| *Linochilus_rhododendroides* | P_rhod_0000_Layer_31 | 94.9634516 | 6.56930047 |
| *Linochilus_rhododendroides* | P_rhod_0001_Layer_30 | 77.4974668 | 6.27607725 |
| *Linochilus_rhododendroides* | P_rhod_0002_Layer_29 | 68.8065892 | 6.10447483 |
| *Linochilus_rhododendroides* | P_rhod_0003_Layer_28 | 53.1524361 | 5.73206391 |
| *Linochilus_rhododendroides* | P_rhod_0004_Layer_27 | 66.8254025 | 6.06232472 |
| *Linochilus_rhododendroides* | P_rhod_0006_Layer_26 | 66.8442197 | 6.0627309 |
| *Linochilus_rhododendroides* | P_rhod_0007_Layer_25 | 99.3389084 | 6.63428699 |
| *Linochilus_rhododendroides* | P_rhod_0008_Layer_24 | 76.0198653 | 6.24830456 |
| *Linochilus_rhododendroides* | P_rhod_0009_Layer_23 | 68.257305 | 6.09291155 |
| *Linochilus_rhododendroides* | P_rhod_0011_Layer_22 | 98.7967926 | 6.6263923 |
| *Linochilus_rhododendroides* | P_rhod_0012_Layer_21 | 59.9481486 | 5.90564329 |
| *Linochilus_rhododendroides* | P_rhod_0013_Layer_20 | 62.3083683 | 5.96135403 |
| *Linochilus_rhododendroides* | P_rhod_0014_Layer_19 | 75.4571402 | 6.23758552 |
| *Linochilus_rhododendroides* | P_rhod_0015_Layer_18 | 55.1479598 | 5.78523561 |
| *Linochilus_rhododendroides* | P_rhod_0017_Layer_17 | 106.345195 | 6.73261104 |
| *Linochilus_rhododendroides* | P_rhod_0018_Layer_16 | 65.379163 | 6.030759 |
| *Linochilus_rhododendroides* | P_rhod_0019_Layer_15 | 72.7698588 | 6.18526911 |
| *Linochilus_rhododendroides* | P_rhod_0020_Layer_14 | 72.3254135 | 6.17643076 |
| *Linochilus_rhododendroides* | P_rhod_0021_Layer_13 | 85.6229311 | 6.41992532 |
| *Linochilus_rhododendroides* | P_rhod_0023_Layer_12 | 78.4374329 | 6.29347041 |
| *Linochilus_rhododendroides* | P_rhod_0024_Layer_11 | 145.146348 | 7.18136446 |
| *Linochilus_rhododendroides* | P_rhod_0025_Layer_10 | 117.993964 | 6.88256925 |
| *Linochilus_rhododendroides* | P_rhod_0026_Layer_9 | 113.025316 | 6.82050214 |
| *Linochilus_rhododendroides* | P_rhod_0027_Layer_8 | 161.253907 | 7.3331903 |
| *Linochilus_rhododendroides* | P_rhod_0029_Layer_7 | 170.141021 | 7.41058721 |
| *Linochilus_rhododendroides* | P_rhod_0030_Layer_6 | 84.3845774 | 6.39890744 |
| *Linochilus_rhododendroides* | P_rhod_0031_Layer_5 | 50.8173059 | 5.66724799 |
| *Linochilus_rhododendroides* | P_rhod_0032_Layer_4 | 87.8559105 | 6.45706744 |
| *Linochilus_rhododendroides* | P_rhod_0033_Layer_3 | 87.8657671 | 6.45722929 |
| *Linochilus_rhododendroides* | P_rhod_0034_Layer_2 | 75.6049899 | 6.24040955 |
| *Linochilus_rhomboidalis_COL* | P_rhom_c_0000_Layer_31 | 21.4229819 | 4.4210874 |
| *Linochilus_rhomboidalis_COL* | P_rhom_c_0001_Layer_30 | 19.2706479 | 4.26833317 |
| *Linochilus_rhomboidalis_COL* | P_rhom_c_0002_Layer_29 | 20.469575 | 4.35540924 |
| *Linochilus_rhomboidalis_COL* | P_rhom_c_0003_Layer_28 | 22.6075721 | 4.49873416 |
| *Linochilus_rhomboidalis_COL* | P_rhom_c_0004_Layer_27 | 22.3002238 | 4.47898628 |
| *Linochilus_rhomboidalis_COL* | P_rhom_c_0006_Layer_26 | 47.7250058 | 5.57667347 |
| *Linochilus_rhomboidalis_COL* | P_rhom_c_0007_Layer_25 | 43.1022367 | 5.42969083 |
| *Linochilus_rhomboidalis_COL* | P_rhom_c_0008_Layer_24 | 51.4615725 | 5.68542364 |
| *Linochilus_rhomboidalis_COL* | P_rhom_c_0009_Layer_23 | 42.5896909 | 5.41243235 |
| *Linochilus_rhomboidalis_COL* | P_rhom_c_0010_Layer_22 | 68.0010321 | 6.08748474 |
| *Linochilus_rhomboidalis_COL* | P_rhom_c_0012_Layer_21 | 30.8405635 | 4.94675722 |
| *Linochilus_rhomboidalis_COL* | P_rhom_c_0013_Layer_20 | 32.2034694 | 5.00914422 |
| *Linochilus_rhomboidalis_COL* | P_rhom_c_0014_Layer_19 | 27.8548944 | 4.79985894 |
| *Linochilus_rhomboidalis_COL* | P_rhom_c_0015_Layer_18 | 30.2536447 | 4.91903705 |
| *Linochilus_rhomboidalis_COL* | P_rhom_c_0016_Layer_17 | 23.3898317 | 4.54780958 |
| *Linochilus_rhomboidalis_COL* | P_rhom_c_0018_Layer_6 | 38.8656691 | 5.28042445 |
| *Linochilus_rhomboidalis_COL* | P_rhom_c_0019_Layer_5 | 53.9956277 | 5.75477068 |
| *Linochilus_rhomboidalis_COL* | P_rhom_c_0020_Layer_16 | 27.6470087 | 4.78905149 |
| *Linochilus_rhomboidalis_COL* | P_rhom_c_0021_Layer_15 | 24.030514 | 4.5867956 |
| *Linochilus_rhomboidalis_COL* | P_rhom_c_0022_Layer_14 | 19.3871356 | 4.27702776 |
| *Linochilus_rhomboidalis_COL* | P_rhom_c_0023_Layer_13 | 31.6470167 | 4.9839976 |
| *Linochilus_rhomboidalis_COL* | P_rhom_c_0025_Layer_12 | 24.4265722 | 4.61037952 |
| *Linochilus_rhomboidalis_COL* | P_rhom_c_0026_Layer_11 | 45.863891 | 5.51928685 |
| *Linochilus_rhomboidalis_COL* | P_rhom_c_0027_Layer_10 | 44.5063614 | 5.47593965 |
| *Linochilus_rhomboidalis_COL* | P_rhom_c_0028_Layer_9 | 55.5260967 | 5.79509408 |
| *Linochilus_rhomboidalis_COL* | P_rhom_c_0029_Layer_8 | 50.39347 | 5.65516489 |
| *Linochilus_rhomboidalis_COL* | P_rhom_c_0031_Layer_7 | 49.8343291 | 5.639068 |
| *Linochilus_rhomboidalis_COL* | P_rhom_c_0032_Layer_4 | 27.257223 | 4.76856668 |
| *Linochilus_rhomboidalis_COL* | P_rhom_c_0033_Layer_3 | 31.1765857 | 4.96239103 |
| *Linochilus_rhomboidalis_COL* | P_rhom_c_0034_Layer_2 | 37.893445 | 5.2438764 |
| *Linochilus_rhomboidalis_ECU* | P_rhom_e_0000_Layer_31 | 39.3656701 | 5.29886613 |
| *Linochilus_rhomboidalis_ECU* | P_rhom_e_0001_Layer_30 | 43.4534919 | 5.44140021 |
| *Linochilus_rhomboidalis_ECU* | P_rhom_e_0002_Layer_29 | 29.8387694 | 4.89911613 |
| *Linochilus_rhomboidalis_ECU* | P_rhom_e_0003_Layer_28 | 21.000042 | 4.39232031 |
| *Linochilus_rhomboidalis_ECU* | P_rhom_e_0005_Layer_27 | 28.3988023 | 4.82775818 |
| *Linochilus_rhomboidalis_ECU* | P_rhom_e_0006_Layer_26 | 46.7993768 | 5.54841741 |
| *Linochilus_rhomboidalis_ECU* | P_rhom_e_0007_Layer_25 | 50.591499 | 5.66082308 |
| *Linochilus_rhomboidalis_ECU* | P_rhom_e_0008_Layer_24 | 53.6757346 | 5.74619813 |
| *Linochilus_rhomboidalis_ECU* | P_rhom_e_0009_Layer_23 | 32.1049029 | 5.00472173 |
| *Linochilus_rhomboidalis_ECU* | P_rhom_e_0010_Layer_22 | 45.3692664 | 5.50364343 |
| *Linochilus_rhomboidalis_ECU* | P_rhom_e_0012_Layer_21 | 20.1120474 | 4.32998805 |
| *Linochilus_rhomboidalis_ECU* | P_rhom_e_0013_Layer_20 | 30.1855442 | 4.91578591 |
| *Linochilus_rhomboidalis_ECU* | P_rhom_e_0014_Layer_19 | 20.424772 | 4.35224807 |
| *Linochilus_rhomboidalis_ECU* | P_rhom_e_0015_Layer_18 | 35.060106 | 5.13175845 |
| *Linochilus_rhomboidalis_ECU* | P_rhom_e_0016_Layer_17 | 28.3844654 | 4.82702966 |
| *Linochilus_rhomboidalis_ECU* | P_rhom_e_0018_Layer_16 | 20.8235184 | 4.38014194 |
| *Linochilus_rhomboidalis_ECU* | P_rhom_e_0019_Layer_15 | 22.9919815 | 4.5230589 |
| *Linochilus_rhomboidalis_ECU* | P_rhom_e_0020_Layer_14 | 23.560979 | 4.55832758 |
| *Linochilus_rhomboidalis_ECU* | P_rhom_e_0021_Layer_13 | 30.7921763 | 4.94449193 |
| *Linochilus_rhomboidalis_ECU* | P_rhom_e_0023_Layer_12 | 31.2894891 | 4.9676062 |
| *Linochilus_rhomboidalis_ECU* | P_rhom_e_0024_Layer_11 | 41.5072515 | 5.3752915 |
| *Linochilus_rhomboidalis_ECU* | P_rhom_e_0025_Layer_10 | 33.4194217 | 5.06261486 |
| *Linochilus_rhomboidalis_ECU* | P_rhom_e_0026_Layer_9 | 32.379097 | 5.01699085 |
| *Linochilus_rhomboidalis_ECU* | P_rhom_e_0027_Layer_8 | 30.6174448 | 4.93628198 |
| *Linochilus_rhomboidalis_ECU* | P_rhom_e_0029_Layer_7 | 32.2536487 | 5.01139047 |
| *Linochilus_rhomboidalis_ECU* | P_rhom_e_0030_Layer_6 | 37.9740903 | 5.2469435 |
| *Linochilus_rhomboidalis_ECU* | P_rhom_e_0031_Layer_5 | 28.1183358 | 4.81343931 |
| *Linochilus_rhomboidalis_ECU* | P_rhom_e_0032_Layer_4 | 41.4346707 | 5.37276655 |
| *Linochilus_rhomboidalis_ECU* | P_rhom_e_0033_Layer_3 | 43.5816284 | 5.4456482 |
| *Linochilus_rhomboidalis_ECU* | P_rhom_e_0034_Layer_2 | 33.5851926 | 5.0697534 |
| *Linochilus_romeroi* | P_rome_0000_Layer_7 | 386.986437 | 8.59613919 |
| *Linochilus_romeroi* | P_rome_0001_Layer_6 | 447.644265 | 8.80620889 |
| *Linochilus_romeroi* | P_rome_0002_Layer_5 | 811.533881 | 9.66450752 |
| *Linochilus_romeroi* | P_rome_0003_Layer_4 | 767.363542 | 9.58376641 |
| *Linochilus_romeroi* | P_rome_0004_Layer_3 | 439.009839 | 8.77810946 |
| *Linochilus_romeroi* | P_rome_0005_Layer_2 | 569.277124 | 9.15298732 |
| *Linochilus_rosmarinifolius* | P_rosm_0000_Layer_31 | 21.3109745 | 4.41352466 |
| *Linochilus_rosmarinifolius* | P_rosm_0001_Layer_30 | 19.3817592 | 4.27662762 |
| *Linochilus_rosmarinifolius* | P_rosm_0002_Layer_29 | 14.696266 | 3.87737773 |
| *Linochilus_rosmarinifolius* | P_rosm_0004_Layer_28 | 29.1792698 | 4.86687188 |
| *Linochilus_rosmarinifolius* | P_rosm_0005_Layer_27 | 18.4265602 | 4.20371487 |
| *Linochilus_rosmarinifolius* | P_rosm_0006_Layer_25 | 30.3047201 | 4.92147061 |
| *Linochilus_rosmarinifolius* | P_rosm_0007_Layer_26 | 25.9875072 | 4.69974635 |
| *Linochilus_rosmarinifolius* | P_rosm_0008_Layer_24 | 30.6398462 | 4.93733715 |
| *Linochilus_rosmarinifolius* | P_rosm_0009_Layer_23 | 32.8728256 | 5.03882356 |
| *Linochilus_rosmarinifolius* | P_rosm_0010_Layer_22 | 24.5520204 | 4.61776985 |
| *Linochilus_rosmarinifolius* | P_rosm_0012_Layer_21 | 30.5896669 | 4.93497249 |
| *Linochilus_rosmarinifolius* | P_rosm_0013_Layer_20 | 26.2805185 | 4.71592183 |
| *Linochilus_rosmarinifolius* | P_rosm_0014_Layer_19 | 23.4220899 | 4.5497979 |
| *Linochilus_rosmarinifolius* | P_rosm_0015_Layer_18 | 21.681943 | 4.43842214 |
| *Linochilus_rosmarinifolius* | P_rosm_0016_Layer_17 | 25.7321303 | 4.685499 |
| *Linochilus_rosmarinifolius* | P_rosm_0019_Layer_15 | 10.9059358 | 3.44704166 |
| *Linochilus_rosmarinifolius* | P_rosm_0020_Layer_16 | 11.362926 | 3.50626247 |
| *Linochilus_rosmarinifolius* | P_rosm_0021_Layer_14 | 10.0107727 | 3.32348143 |
| *Linochilus_rosmarinifolius* | P_rosm_0022_Layer_13 | 7.76883274 | 2.95769785 |
| *Linochilus_rosmarinifolius* | P_rosm_0024_Layer_12 | 10.3987663 | 3.37834048 |
| *Linochilus_rosmarinifolius* | P_rosm_0025_Layer_11 | 65.5422458 | 6.0343532 |
| *Linochilus_rosmarinifolius* | P_rosm_0026_Layer_10 | 88.4732952 | 6.46717015 |
| *Linochilus_rosmarinifolius* | P_rosm_0027_Layer_9 | 95.0046703 | 6.56992653 |
| *Linochilus_rosmarinifolius* | P_rosm_0028_Layer_8 | 80.6139605 | 6.3329578 |
| *Linochilus_rosmarinifolius* | P_rosm_0029_Layer_7 | 104.530675 | 6.70778256 |
| *Linochilus_rosmarinifolius* | P_rosm_0030_Layer_6 | 31.2715679 | 4.96677965 |
| *Linochilus_rosmarinifolius* | P_rosm_0031_Layer_5 | 27.0574018 | 4.7579514 |
| *Linochilus_rosmarinifolius* | P_rosm_0032_Layer_4 | 27.2464703 | 4.76799744 |
| *Linochilus_rosmarinifolius* | P_rosm_0033_Layer_3 | 28.060988 | 4.8104939 |
| *Linochilus_rosmarinifolius* | P_rosm_0034_Layer_2 | 40.4687189 | 5.33873527 |
| *Linochilus_rupestris* | P_rupe_0000_Layer_26 | 95.0342403 | 6.5703755 |
| *Linochilus_rupestris* | P_rupe_0001_Layer_25 | 156.503897 | 7.29005477 |
| *Linochilus_rupestris* | P_rupe_0002_Layer_24 | 190.377621 | 7.57272009 |
| *Linochilus_rupestris* | P_rupe_0003_Layer_23 | 99.8559345 | 6.64177627 |
| *Linochilus_rupestris* | P_rupe_0004_Layer_31 | 73.3612578 | 6.19694647 |
| *Linochilus_rupestris* | P_rupe_0005_Layer_30 | 70.8585647 | 6.14687034 |
| *Linochilus_rupestris* | P_rupe_0006_Layer_29 | 103.421354 | 6.69239028 |
| *Linochilus_rupestris* | P_rupe_0007_Layer_28 | 89.1471317 | 6.47811647 |
| *Linochilus_rupestris* | P_rupe_0009_Layer_27 | 66.8872306 | 6.06365891 |
| *Linochilus_rupestris* | P_rupe_0010_Layer_22 | 143.922331 | 7.16914665 |
| *Linochilus_rupestris* | P_rupe_0012_Layer_21 | 28.87909 | 4.85195338 |
| *Linochilus_rupestris* | P_rupe_0013_Layer_20 | 97.6901595 | 6.61014134 |
| *Linochilus_rupestris* | P_rupe_0014_Layer_19 | 44.5690856 | 5.47797146 |
| *Linochilus_rupestris* | P_rupe_0015_Layer_18 | 45.5198043 | 5.50842245 |
| *Linochilus_rupestris* | P_rupe_0016_Layer_17 | 94.2098658 | 6.55780624 |
| *Linochilus_rupestris* | P_rupe_0019_Layer_16 | 150.727004 | 7.2357941 |
| *Linochilus_rupestris* | P_rupe_0020_Layer_15 | 261.530093 | 8.03083315 |
| *Linochilus_rupestris* | P_rupe_0021_Layer_14 | 162.198353 | 7.34161536 |
| *Linochilus_rupestris* | P_rupe_0022_Layer_13 | 59.5691156 | 5.89649263 |
| *Linochilus_rupestris* | P_rupe_0023_Layer_12 | 73.4580322 | 6.19884834 |
| *Linochilus_rupestris* | P_rupe_0025_Layer_11 | 38.5108297 | 5.2671923 |
| *Linochilus_rupestris* | P_rupe_0026_Layer_10 | 51.5529705 | 5.68798365 |
| *Linochilus_rupestris* | P_rupe_0027_Layer_9 | 62.636326 | 5.96892769 |
| *Linochilus_rupestris* | P_rupe_0028_Layer_8 | 65.3388404 | 6.02986894 |
| *Linochilus_rupestris* | P_rupe_0029_Layer_7 | 74.9920855 | 6.22866644 |
| *Linochilus_rupestris* | P_rupe_0031_Layer_6 | 90.1121874 | 6.49365033 |
| *Linochilus_rupestris* | P_rupe_0032_Layer_5 | 76.8988993 | 6.26489104 |
| *Linochilus_rupestris* | P_rupe_0033_Layer_4 | 113.312951 | 6.82416895 |
| *Linochilus_rupestris* | P_rupe_0034_Layer_3 | 105.887309 | 6.72638587 |
| *Linochilus_rupestris* | P_rupe_0036_Layer_1 | 74.9168165 | 6.22721769 |
| *Linochilus_schultzii_CAL* | P_schu_0000_Layer_29 | 90.5539446 | 6.50070558 |
| *Linochilus_schultzii_CAL* | P_schu_0002_Layer_31 | 88.9589593 | 6.47506801 |
| *Linochilus_schultzii_CAL* | P_schu_0003_Layer_32 | 73.0180672 | 6.19018158 |
| *Linochilus_schultzii_CAL* | P_schu_0004_Layer_30 | 97.5360374 | 6.60786346 |
| *Linochilus_schultzii_CAL* | P_schu_0005_Layer_28 | 89.8155918 | 6.48889401 |
| *Linochilus_schultzii_CAL* | P_schu_0006_Layer_27 | 48.4938246 | 5.59972913 |
| *Linochilus_schultzii_CAL* | P_schu_0007_Layer_26 | 48.8325349 | 5.60977077 |
| *Linochilus_schultzii_CAL* | P_schu_0008_Layer_25 | 49.7187374 | 5.63571776 |
| *Linochilus_schultzii_CAL* | P_schu_0009_Layer_24 | 44.787724 | 5.48503145 |
| *Linochilus_schultzii_CAL* | P_schu_0011_Layer_23 | 63.5270088 | 5.98929818 |
| *Linochilus_schultzii_CAL* | P_schu_0012_Layer_22 | 88.3308218 | 6.46484503 |
| *Linochilus_schultzii_CAL* | P_schu_0013_Layer_21 | 54.5377435 | 5.7691831 |
| *Linochilus_schultzii_CAL* | P_schu_0014_Layer_20 | 62.4606984 | 5.9648768 |
| *Linochilus_schultzii_CAL* | P_schu_0015_Layer_19 | 110.071009 | 6.78229072 |
| *Linochilus_schultzii_CAL* | P_schu_0017_Layer_18 | 73.7152012 | 6.20389025 |
| *Linochilus_schultzii_CAL* | P_schu_0018_Layer_17 | 58.4230559 | 5.86846592 |
| *Linochilus_schultzii_CAL* | P_schu_0019_Layer_16 | 83.9213148 | 6.39096538 |
| *Linochilus_schultzii_CAL* | P_schu_0020_Layer_15 | 64.1004866 | 6.0022634 |
| *Linochilus_schultzii_CAL* | P_schu_0021_Layer_14 | 70.197273 | 6.13334308 |
| *Linochilus_schultzii_CAL* | P_schu_0023_Layer_13 | 44.3262535 | 5.47008953 |
| *Linochilus_schultzii_CAL* | P_schu_0024_Layer_12 | 81.3576896 | 6.3462068 |
| *Linochilus_schultzii_CAL* | P_schu_0025_Layer_11 | 81.7528517 | 6.35319715 |
| *Linochilus_schultzii_CAL* | P_schu_0026_Layer_10 | 84.7609222 | 6.40532738 |
| *Linochilus_schultzii_CAL* | P_schu_0027_Layer_9 | 75.5745239 | 6.23982808 |
| *Linochilus_schultzii_CAL* | P_schu_0029_Layer_8 | 61.1318427 | 5.93385215 |
| *Linochilus_schultzii_CAL* | P_schu_0031_Layer_6 | 49.2205286 | 5.62118825 |
| *Linochilus_schultzii_CAL* | P_schu_0032_Layer_5 | 64.5377635 | 6.01207168 |
| *Linochilus_schultzii_CAL* | P_schu_0033_Layer_4 | 95.6274314 | 6.57935262 |
| *Linochilus_schultzii_CAL* | P_schu_0034_Layer_3 | 75.0951323 | 6.23064749 |
| *Linochilus_schultzii_CAL* | P_schu_0035_Layer_2 | 68.0467311 | 6.08845395 |
| *Linochilus_sp_nov* | P_spno_0000_Layer_10 | 2929.74511 | 11.5165594 |
| *Linochilus_sp_nov* | P_spno_0001_Layer_8 | 2898.20741 | 11.5009451 |
| *Linochilus_sp_nov* | P_spno_0002_Layer_12 | 1616.58746 | 10.6587358 |
| *Linochilus_sp_nov* | P_spno_0003_Layer_11 | 2002.42515 | 10.9675326 |
| *Linochilus_sp_nov* | P_spno_0005_Layer_9 | 3282.10692 | 11.6804065 |
| *Linochilus_sp_nov* | P_spno_0006_Layer_7 | 3087.19614 | 11.5920814 |
| *Linochilus_tachirensis* | P_tach_0000_Layer_12 | 560.469759 | 9.13049272 |
| *Linochilus_tachirensis* | P_tach_0001_Layer_11 | 550.061136 | 9.10344816 |
| *Linochilus_tachirensis* | P_tach_0002_Layer_15 | 840.087702 | 9.71439614 |
| *Linochilus_tachirensis* | P_tach_0003_Layer_14 | 749.993436 | 9.55073416 |
| *Linochilus_tachirensis* | P_tach_0004_Layer_13 | 775.057106 | 9.5981588 |
| *Linochilus_tachirensis* | P_tach_0005_Layer_10 | 978.923105 | 9.93505173 |
| *Linochilus_tachirensis* | P_tach_0007_Layer_20 | 1170.2837 | 10.1926426 |
| *Linochilus_tachirensis* | P_tach_0008_Layer_19 | 1240.93438 | 10.2772111 |
| *Linochilus_tachirensis* | P_tach_0009_Layer_18 | 1487.30584 | 10.5384856 |
| *Linochilus_tachirensis* | P_tach_0011_Layer_9 | 637.253963 | 9.31572463 |
| *Linochilus_tachirensis* | P_tach_0012_Layer_8 | 806.581362 | 9.65567626 |
| *Linochilus_tachirensis* | P_tach_0013_Layer_7 | 830.162055 | 9.69724918 |
| *Linochilus_tachirensis* | P_tach_0014_Layer_6 | 1048.11052 | 10.0335751 |
| *Linochilus_tachirensis* | P_tach_0016_Layer_5 | 1142.85444 | 10.1584259 |
| *Linochilus_tachirensis* | P_tach_0017_Layer_4 | 668.27553 | 9.38429924 |
| *Linochilus_tachirensis* | P_tach_0018_Layer_3 | 871.992783 | 9.76817239 |
| *Linochilus_tachirensis* | P_tach_0019_Layer_2 | 1435.20807 | 10.4870442 |
| *Linochilus_tachirensis* | P_tach_0021_Layer_17 | 1420.7394 | 10.4724262 |
| *Linochilus_tachirensis* | P_tach_0023_Layer_16 | 1096.02728 | 10.098068 |
| *Linochilus_tachirensis* | P_tach_0025_Layer_23 | 585.461744 | 9.19343109 |
| *Linochilus_tachirensis* | P_tach_0027_Layer_26 | 780.592063 | 9.60842498 |
| *Linochilus_tachirensis* | P_tach_0028_Layer_25 | 1251.91738 | 10.2899236 |
| *Linochilus_tachirensis* | P_tach_0029_Layer_24 | 326.734524 | 8.35197509 |
| *Linochilus_tachirensis* | P_tach_0030_Layer_22 | 702.598179 | 9.45655603 |
| *Linochilus_tenuifolius* | P_tenu_0000_Layer_29 | 794.092986 | 9.63316414 |
| *Linochilus_tenuifolius* | P_tenu_0001_Layer_28 | 1566.97715 | 10.6137684 |
| *Linochilus_tenuifolius* | P_tenu_0002_Layer_27 | 1019.70634 | 9.99393802 |
| *Linochilus_tenuifolius* | P_tenu_0003_Layer_26 | 1234.67003 | 10.2699098 |
| *Linochilus_tenuifolius* | P_tenu_0005_Layer_25 | 1392.01712 | 10.4429612 |
| *Linochilus_tenuifolius* | P_tenu_0006_Layer_24 | 1853.33166 | 10.8559054 |
| *Linochilus_tenuifolius* | P_tenu_0007_Layer_23 | 1655.45403 | 10.6930112 |
| *Linochilus_tenuifolius* | P_tenu_0008_Layer_22 | 1446.61042 | 10.4984607 |
| *Linochilus_tenuifolius* | P_tenu_0009_Layer_21 | 711.280096 | 9.47427398 |
| *Linochilus_tenuifolius* | P_tenu_0010_Layer_19 | 1208.59292 | 10.2391127 |
| *Linochilus_tenuifolius* | P_tenu_0011_Layer_18 | 3962.75434 | 11.9522878 |
| *Linochilus_tenuifolius* | P_tenu_0012_Layer_17 | 3045.18082 | 11.5723122 |
| *Linochilus_tenuifolius* | P_tenu_0013_Layer_16 | 1803.44716 | 10.8165414 |
| *Linochilus_tenuifolius* | P_tenu_0014_Layer_15 | 1952.51287 | 10.9311163 |
| *Linochilus_tenuifolius* | P_tenu_0015_Layer_14 | 2386.40083 | 11.2206207 |
| *Linochilus_tenuifolius* | P_tenu_0018_Layer_20 | 1269.4443 | 10.3099814 |
| *Linochilus_tenuifolius* | P_tenu_0020_Layer_9 | 678.959243 | 9.40718116 |
| *Linochilus_tenuifolius* | P_tenu_0021_Layer_13 | 1323.23741 | 10.3698562 |
| *Linochilus_tenuifolius* | P_tenu_0022_Layer_12 | 1353.68909 | 10.4026807 |
| *Linochilus_tenuifolius* | P_tenu_0023_Layer_11 | 1174.76669 | 10.1981585 |
| *Linochilus_tenuifolius* | P_tenu_0024_Layer_10 | 930.653295 | 9.8621 |
| *Linochilus_tenuifolius* | P_tenu_0025_Layer_8 | 1180.93516 | 10.205714 |
| *Linochilus_tenuifolius* | P_tenu_0026_Layer_6 | 1116.54435 | 10.1248248 |
| *Linochilus_tenuifolius* | P_tenu_0027_Layer_7 | 1116.56496 | 10.1248515 |
| *Linochilus_tenuifolius* | P_tenu_0029_Layer_5 | 1193.80436 | 10.2213507 |
| *Linochilus_tenuifolius* | P_tenu_0030_Layer_4 | 573.852402 | 9.16453591 |
| *Linochilus_tenuifolius* | P_tenu_0031_Layer_3 | 608.557669 | 9.24925017 |
| *Linochilus_tenuifolius* | P_tenu_0032_Layer_2 | 1931.42232 | 10.9154479 |
| *Linochilus_venezuelensis* | P_vene_0000_Layer_32 | 73.7994308 | 6.20553778 |
| *Linochilus_venezuelensis* | P_vene_0001_Layer_31 | 111.924059 | 6.80637638 |
| *Linochilus_venezuelensis* | P_vene_0002_Layer_30 | 139.437555 | 7.12347537 |
| *Linochilus_venezuelensis* | P_vene_0003_Layer_29 | 140.827343 | 7.13778366 |
| *Linochilus_venezuelensis* | P_vene_0004_Layer_28 | 65.8379453 | 6.04084741 |
| *Linochilus_venezuelensis* | P_vene_0006_Layer_27 | 88.9598553 | 6.47508254 |
| *Linochilus_venezuelensis* | P_vene_0007_Layer_26 | 104.355944 | 6.70536896 |
| *Linochilus_venezuelensis* | P_vene_0008_Layer_25 | 75.18205 | 6.23231635 |
| *Linochilus_venezuelensis* | P_vene_0009_Layer_24 | 78.2205866 | 6.28947645 |
| *Linochilus_venezuelensis* | P_vene_0010_Layer_23 | 73.0816874 | 6.19143804 |
| *Linochilus_venezuelensis* | P_vene_0012_Layer_22 | 153.795111 | 7.26486583 |
| *Linochilus_venezuelensis* | P_vene_0013_Layer_21 | 137.330024 | 7.10150326 |
| *Linochilus_venezuelensis* | P_vene_0014_Layer_20 | 156.098879 | 7.28631636 |
| *Linochilus_venezuelensis* | P_vene_0015_Layer_19 | 81.1130655 | 6.34186241 |
| *Linochilus_venezuelensis* | P_vene_0016_Layer_18 | 182.283519 | 7.51004031 |
| *Linochilus_venezuelensis* | P_vene_0017_Layer_16 | 47.4947187 | 5.56969519 |
| *Linochilus_venezuelensis* | P_vene_0019_Layer_17 | 49.2850448 | 5.62307803 |
| *Linochilus_venezuelensis* | P_vene_0020_Layer_14 | 92.1659549 | 6.52616203 |
| *Linochilus_venezuelensis* | P_vene_0021_Layer_13 | 118.189305 | 6.88495568 |
| *Linochilus_venezuelensis* | P_vene_0022_Layer_12 | 71.4571322 | 6.15900611 |
| *Linochilus_venezuelensis* | P_vene_0026_Layer_11 | 55.0789632 | 5.7834295 |
| *Linochilus_venezuelensis* | P_vene_0027_Layer_10 | 78.8603728 | 6.30122862 |
| *Linochilus_venezuelensis* | P_vene_0028_Layer_9 | 104.298596 | 6.70457592 |
| *Linochilus_venezuelensis* | P_vene_0029_Layer_8 | 69.9535449 | 6.12832526 |
| *Linochilus_venezuelensis* | P_vene_0030_Layer_7 | 96.7179354 | 6.59571154 |
| *Linochilus_venezuelensis* | P_vene_0032_Layer_6 | 222.908152 | 7.80030557 |
| *Linochilus_venezuelensis* | P_vene_0033_Layer_5 | 233.909966 | 7.86980952 |
| *Linochilus_venezuelensis* | P_vene_0034_Layer_4 | 193.032644 | 7.59270103 |
| *Linochilus_venezuelensis* | P_vene_0035_Layer_3 | 211.552394 | 7.7248712 |
| *Linochilus_venezuelensis* | P_vene_0036_Layer_2 | 121.006514 | 6.91894091 |
| *Linochilus_violaceus* | P_viol_0001_Layer_28 | 75.2178924 | 6.23300398 |
| *Linochilus_violaceus* | P_viol_0002_Layer_27 | 24.4077549 | 4.60926769 |
| *Linochilus_violaceus* | P_viol_0003_Layer_26 | 30.3602758 | 4.92411299 |
| *Linochilus_violaceus* | P_viol_0004_Layer_25 | 22.3020159 | 4.47910222 |
| *Linochilus_violaceus* | P_viol_0005_Layer_24 | 23.0367844 | 4.52586745 |
| *Linochilus_violaceus* | P_viol_0007_Layer_23 | 27.4462915 | 4.77853932 |
| *Linochilus_violaceus* | P_viol_0008_Layer_22 | 22.78768 | 4.51018215 |
| *Linochilus_violaceus* | P_viol_0009_Layer_21 | 27.544858 | 4.78371112 |
| *Linochilus_violaceus* | P_viol_0010_Layer_20 | 44.0547476 | 5.4612256 |
| *Linochilus_violaceus* | P_viol_0012_Layer_19 | 42.1353889 | 5.39696054 |
| *Linochilus_violaceus* | P_viol_0015_Layer_17 | 38.2653095 | 5.25796516 |
| *Linochilus_violaceus* | P_viol_0016_Layer_16 | 53.3343361 | 5.73699272 |
| *Linochilus_violaceus* | P_viol_0017_Layer_15 | 46.6694482 | 5.5444065 |
| *Linochilus_violaceus* | P_viol_0018_Layer_14 | 50.3925739 | 5.65513924 |
| *Linochilus_violaceus* | P_viol_0019_Layer_13 | 64.5709177 | 6.01281263 |
| *Linochilus_violaceus* | P_viol_0021_Layer_11 | 77.2331294 | 6.27114792 |
| *Linochilus_violaceus* | P_viol_0022_Layer_10 | 66.0879458 | 6.04631525 |
| *Linochilus_violaceus* | P_viol_0023_Layer_9 | 94.9491146 | 6.56908264 |
| *Linochilus_violaceus* | P_viol_0024_Layer_8 | 76.9186126 | 6.26526084 |
| *Linochilus_violaceus* | P_viol_0025_Layer_7 | 85.726874 | 6.42167563 |
| *Linochilus_violaceus* | P_viol_0026_Layer_6 | 64.8675133 | 6.01942423 |
| *Linochilus_violaceus* | P_viol_0027_Layer_5 | 68.9965538 | 6.1084524 |
| *Linochilus_violaceus* | P_viol_0028_Layer_4 | 76.6148486 | 6.25955212 |
| *Linochilus_violaceus* | P_viol_0029_Layer_3 | 58.6381101 | 5.8737667 |
| *Linochilus_violaceus* | P_viol_0030_Layer_2 | 81.7080487 | 6.35240629 |

**APPENDIX S3** Number of individuals and total number of leaves measured for every species.

| **Species** | **n individuals** | **total leaves** |
| --- | --- | --- |
| *Linochilus alveolatus* | 6 | 30 |
| *Linochilus antioquensis* | 2 | 10 |
| *Linochilus apiculatus* | 4 | 24 |
| *Linochilus camargoanus* | 1 | 6 |
| *Linochilus cayambensis* | 6 | 30 |
| *Linochilus cinerascens* | 6 | 30 |
| *Linochilus colombianus* | 4 | 24 |
| *Linochilus coriaceus* | 3 | 10 |
| *Linochilus costaricensis* | 4 | 24 |
| *Linochilus eriophorus* | 5 | 25 |
| *Linochilus floribundus* | 6 | 36 |
| *Linochilus frontinensis* | 2 | 13 |
| *Linochilus glutinosus* | 6 | 29 |
| *Linochilus heterophyllus* | 6 | 31 |
| *Linochilus huertasii* | 4 | 21 |
| *Linochilus inesianus* | 3 | 15 |
| *Linochilus jaramilloi* | 2 | 10 |
| *Linochilus jenesanus* | 3 | 9 |
| *Linochilus juajibioyi* | 5 | 26 |
| *Linochilus lacunosus* | 2 | 12 |
| *Linochilus mutiscuanus* | 2 | 7 |
| *Linochilus oblongifolius* | 2 | 8 |
| *Linochilus obtusus* | 5 | 29 |
| *Linochilus ochraceus* | 6 | 20 |
| *Linochilus phylicoides* | 6 | 30 |
| *Linochilus revolutus* | 6 | 30 |
| *Linochilus rhododendroides* | 6 | 30 |
| *Linochilus rhomboidalis COL* | 6 | 30 |
| *Linochilus rhomboidalis ECU* | 6 | 30 |
| *Linochilus romeroi* | 1 | 6 |
| *Linochilus rosmarinifolius* | 6 | 30 |
| *Linochilus rupestris* | 6 | 30 |
| *Linochilus schultzii CAL* | 6 | 30 |
| *Linochilus sp nov* | 2 | 6 |
| *Linochilus tachirensis* | 6 | 24 |
| *Linochilus tenuifolius* | 6 | 28 |
| *Linochilus venezuelensis* | 6 | 30 |
| *Linochilus violaceus* | 5 | 25 |

**APPENDIX S4** Missing species (bold) from the Vargas et al. (2017) phylogeny and their putative sisters (underlined when present in the sister pair analysis based solely on the phylogeny of Vargas et al. 2017). * Putative additional pair based solely on taxonomy

| Missing species | Most similar species | Distribution | Reference |
| --- | --- | --- | --- |
| **L. anactinotus** | L. inesianus | Sympatric (but likely the same species) * | Cuatrecasas 1969 |
| **L. bicolor** | L. ochraceus | Allopatric * | Cuatrecasas 1969 |
| **L. chrysotrichus** | L. eriophorus | Allopatric | Díaz-Piedrahita and Restrepo 1994 |
| **L. crassifolius** | Unknown |  |  |
| **L. cyparissias** | L. rosmarinifolius | Allopatric | Cuatrecasas 1969 |
| **L. ellipticus** | L. tenuifolius | Allopatric | Cuatrecasas 1969 |
| **L. farallonensis** | L. floribundus | Allopatric |  |
| **L. fosbergii** | Unknown |  |  |
| **L. grantii** | **L. santamartae** | Allopatric * | Cuatrecasas 1982 |
| **L. julianii** | L. huertasii | Allopatric | Cuatrecasas 1969 |
| **L. leiocladus** | Unknown |  |  |
| **L. micradenius** | L. frontinensis | Allopatric * | Cuatrecasas 1991 |
| **L. nevadensis** | Unknown |  |  |
| **L. ocanensis** | Unknown |  |  |
| **L. parvifolius** | Unknown |  |  |
| **L. perijaensis** | L. floribundus | Allopatric | Díaz-Piedrahita and Méndez-Ramirez, 1997; Vargas 2018 |
| **L. pittieri** | Unknown |  |  |
| **L. rangelii** | Unknown |  |  |
| **L. ritterbushii** | Unknown |  |  |
| **L. saxatilis** | L. romeroi | Sympatric | Cuatrecasas 1969 |
| **L. tamanus** | Unknown |  |  |
| **L. tergocanus** | Unknown |  | Cuatrecasas 1969 |
| **L. weddellii** | Unknown |  |  |

**APPENDIX S5**. Sister species comparisons based solely on the phylogeny of Vargas et al. (2017).

| **Sister1** | **Sister 2** | **Wilcox test leaf** | **Distribution** | **Divergence** | **Age** |
| --- | --- | --- | --- | --- | --- |
| L. phylicoides | L. lacunosus | 0.4722 | Allopatric | Geog. isolation | 0.16 (0.01–0.58) |
| L. obtusus | L. venezuelensis | 1.0000 | Sympatric | Inconclusive | 0.22 (0.01–0.81) |
| L. violaceus | L. cinerascens | 0.2897 | Allopatric | Geog. isolation | 0.28 (0.01–0.86) |
| L. rosmarinifolius | L. floribundus | 2.90E-15* | Sympatric | Ecological | 0.55 (0.01–1.49) |
| L. rhomboidalis | L. apiculatus | 2.50E-09* | Allopatric | Geog. isolation | 0.59 (0.01–1.54) |
| L. alveolatus | L. costaricensis | 1.0000 | Allopatric | Geog. isolation | 0.56 (0.01–1.46) |
| L. rhododendroides | L. schultzii | 1.0000 | Allopatric | Geog. isolation | 0.92 (0.02–2.14) |
| L. jenesanus | L. tenuifolius | 1.0000 | Allopatric | Geog. isolation | 0.18 (0.01–0.66) |
| L. oblongifolius | L. mutiscuanus | 0.4140 | Allopatric | Geog. isolation | 0.20 (0.01–0.76) |
| L. sp. nov. ANT | L. antioquensis | 0.1663 | Allopatric | Geog. isolation | 0.52 (0.01–1.47) |
| L. jaramilloi | L. huertasii | 1.0000 | Allopatric | Geog. isolation | 1.41 (0.05–3.18) |
| L. eriophorus | L. rupestris | 1.0000 | Sympatric | Inconclusive | 1.11 (0.04–2.63) |
| L. colombianus | L. glutinosus | 1.20E-11* | Sympatric | Ecological | 2.97 (0.11–6.17) |
| L. coriaceus | L. romeroi | 2.24E-03* | Sympatric | Ecological | 2.22 (0.06–5.42) |

**APPENDIX S6** Range overlap calculated using 0.1 and 0.05 decimal degrees grids. *Linochilus* sister species comparisons based on the phylogeny of Vargas et al. (2017).

|  |  | **Range overlap** | | **Range Asymmetry** | |
| --- | --- | --- | --- | --- | --- |
| **Sister 1** | **Sister 2** | **0.1** | **0.05** | **0.1** | **0.05** |
| *L. phylicoides* | *L. lacunosus* | 0 | 0 | 10.704 | 16.391 |
| *L. obtusus* | *L. venezuelensis* | 0.375 | 0.222 | 1 | 1.112 |
| *L. violaceus* | *L. cinerascens* | 0 | 0 | 5.671 | 6.339 |
| *L. rosmarinifolius* | *L. floribundus* | 0.095 | 0.043 | 1.02 | 1.083 |
| *L. rhomboidalis* | *L. apiculatus* | 0 | 0 | 1.671 | 1.503 |
| *L. alveolatus* | *L. costaricensis* | 0 | 0 | 15.154 | 8.082 |
| *L. rhododendroides* | *L. schultzii* | 0 | 0 | 3.992 | 5.19 |
| *L. jenesanus* | *L. tenuifolius* | 0 | 0 | 5.663 | 6.662 |
| *L. oblongifolius* | *L. mutiscuanus* | 0 | 0 | 3 | 4 |
| *L. sp. nov. ANT* | *L. antioquensis* | 0 | 0 | 1.001 | 1.001 |
| *L. jaramilloi* | *L. huertasii* | 0 | 0 | 3.507 | 3.507 |
| *L. eriophorus* | *L. rupestris* | 0.714 | 0.444 | 2 | 1.889 |
| *L. colombianus* | *L. glutinosus* | 0.2 | 0.2 | 2.597 | 2.597 |
| *L. coriaceus* | *L. romeroi* | 0 | 0 | 2 | 1 |

**APPENDIX S7** Distribution of sister species based on the phylogeny Vargas et al. (2017). Darker pixels indicate higher elevations, orange polygons delineate paramo complexes. ***** : allopatric based on páramo islands. **+** : allopatric based on 0.1 decimal degree grid analysis.
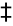
 : allopatric based on 0.05 decimal degree grid analysis. Pair *L. oblongifolius–L. mutiscuanus* is codified as allopatric in the island framework as they inhabit different slopes in the mountain they co-occur.

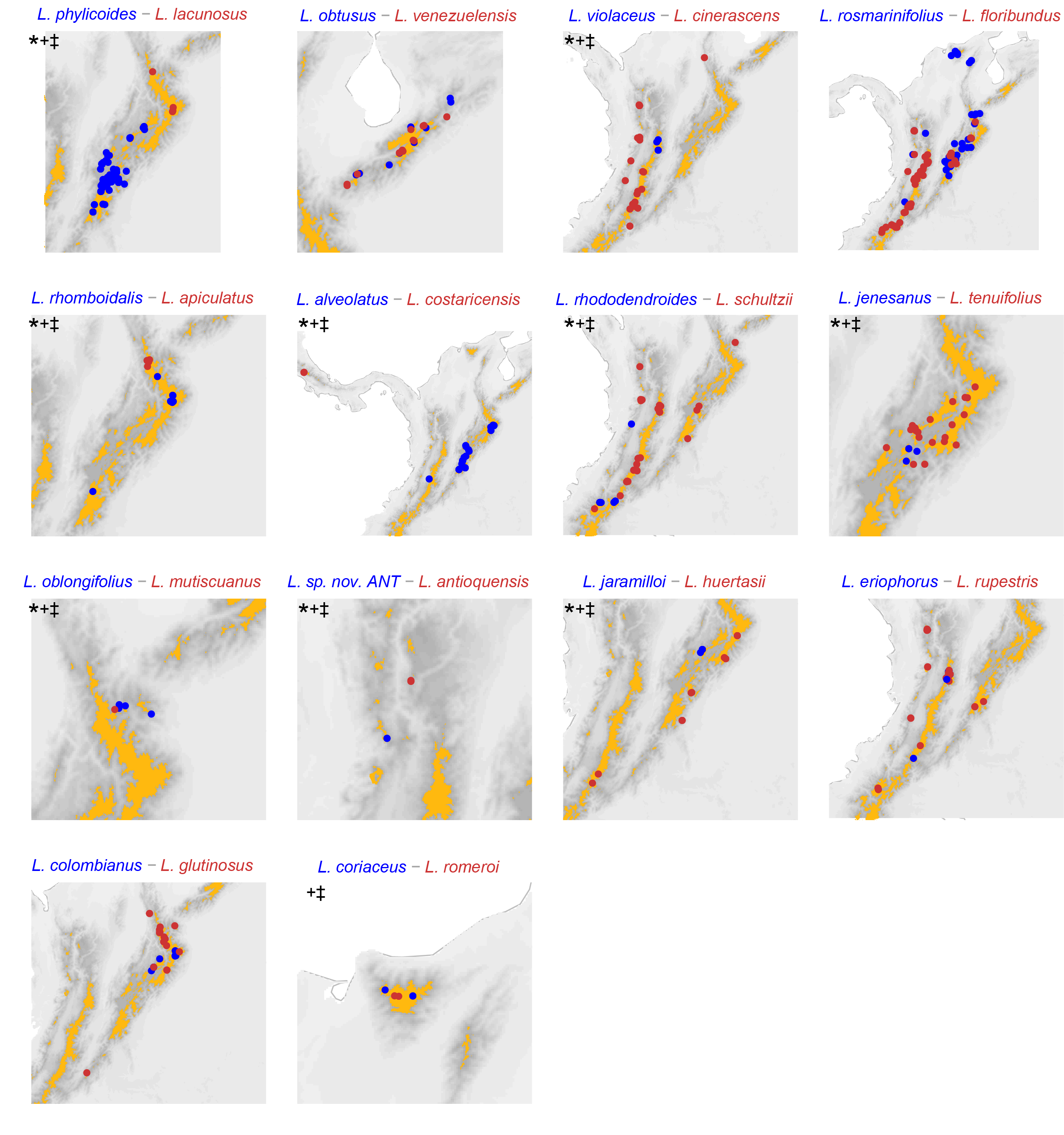
